# Cell-autonomous and non-cell-autonomous effect of TRIM72 on ALS disease progression

**DOI:** 10.1101/2025.01.09.632080

**Authors:** Xue Zhang, Ji He, Ning Song, Xipeng Li, Weizhe Wang, Yueting Hu, Qing Li, Yiqing Hu, Chunmu Yin, Peng Zhao, Xinyi Jiang, Xuejiao Piao, Dawei Meng, Hongmei Cheng, Zhaohui Chen, Xusheng Huang, Pan Yu, Xinming Qi, Jin Ren, Li Luo, Lin Peng, Wei Guo, Tong Wang, Haipeng Gong, Dongsheng Fan, Yichang Jia

**Affiliations:** Tsinghua-Peking Joint Center for Life Sciences; School of Life Sciences, Tsinghua University; School of Basic Medical Sciences, Medical Science Building, Room D204, Tsinghua University, Beijing, China, 100084; IDG/McGovern Institute for Brain Research at Tsinghua; Neurological Department of Peking University Third Hospital; School of Life Science and Technology, Shanghai Tech University, Shanghai, China; Xingtai People’s Hosptial; SineuGene Therapeutics Co., ltd.; Tsinghua Laboratory of Brain and Intelligence; Neurology Department, The First Medical Center, Chinese PLA General Hospital.; Center for Drug Safety Evaluation and Research (CDSER)Shanghai Institute of Materia Medica, Chinese Academy of Sciences.

**Keywords:** Amyotrophic Lateral Sclerosis (ALS), RNA-binding protein, FUS, TRIM72

## Abstract

Dysfunction of RNA-binding proteins, including TDP-43 and FUS, has been associated with amyotrophic lateral sclerosis (ALS); however, the underlying mechanisms are largely unknown. Here, we reported that a neuronal upregulation of TRIM72 (Tripartite Motif Containing 72) in FUS mutation knockin ALS models slows disease progression. TRIM72 interacts with Commander, a protein complex for recycling of membrane proteins, facilitating membrane repair and antioxidation. Exosomal TRIM72 is detected in ALS patient cerebrospinal fluid (CSF) and extracellular application of exosomal TRIM72 protects cell from membrane damage. In a sporadic ALS cohort, CSF TRIM72 level associates ALS disease progression. AAV-mediated neuronal expression of TRIM72 slows down the disease progressions in ALS models and in an ALS patient without adverse effects over a year treatment course. Taken together, our results suggest a universal neuronal protection of a TRIM family protein in cell-autonomous and non-cell-autonomous manners in ALS.

## INTRODUCTION

Amyotrophic lateral sclerosis (ALS) is a rare neurodegenerative disorder marked by motor neuron degeneration and death in the CNS and the spinal cord (Taylor et al., 2016). RNA-binding protein (RBP) dysfunction recently emerged as a common feature of ALS pathology. Many ALS-associated genes encode RBPs, such as TDP-43, FUS and TIA1, and ALS-associated variants in these genes were found to be associated with or causative of ALS pathogenesis (Ling et al., 2013; Taylor et al., 2016). However, the mechanisms underlying their contribution to ALS pathology remain elusive. We have previously established a FUS p.R521C knock-in mouse model that carries an ALS patient FUS mutation and late-onset motor neuron loss (Zhang et al., 2020). However, the FUS-R521C knock-in mouse does not display cytosolic FUS mislocalization and ALS-like symptoms, such as paralysis, muscle weakness, and body sweight loss, which are often seen in patients.

The TRIM (Tripartite Motif) protein family comprises over 80 evolutionarily conserved proteins among the metazoan kingdom with common E3 ubiquitin ligase activities (Hatakeyama, 2017; Ozato et al., 2008). Several well-characterized domains are commonly shared between TRIM family proteins and define the TRIM motif, or otherwise known as RBCC (RING B-box Coiled-Coil) motif: 1) RING domain associates with chaperones and attributes to flexibility of the TRIM protein, 2) one or two B-box domain(s) are zinc-binding motifs structurally similar to RING domains, and 3) Coiled-Coil domain often self-associates to form oligomers. Recently, deficits in TRIM family members, including TRIM11 and TRIM21, have been associated with neurodegenerative diseases. TRIM11 acts as a chaperone to prevent aggregate formation, confers disaggregase activity and degrades aberrant proteins by conferring SUMO ligase activities in PD (Parkinson’s disease) (Zhu et al., 2020) and AD (Alzheimer’s disease) pathogenesis (Zhang et al., 2023). TRIM21 neutralizes tau seeding in AD via its cytosolic antibody receptors in a VCP- and proteasomal-dependent manner (McEwan et al., 2017; Mukadam et al., 2023). While the majority of known functions in neurodegenerative diseases of TRIM11 and TRIM21 are E3 ligase activity-dependent, other TRIM family protein contributions and their E3-independent roles to neurodegenerative disease pathology remain unknown.

TRIM72 (Tripartite Motif Containing 72), also known as MG53, is a member of this family and is encoded by the *Mg53* gene located at the 16p11.2 locus (Cai et al., 2009). TRIM72 was reported to play a critical role in mediating acute membrane repair in skeletal muscle cells, and was also reported to mediate many physiological cellular functions (Balendra and Isaacs, 2018; Cai et al., 2009; Song et al., 2013; Wu et al., 2019; Zhong et al., 2021). Like TRIM11 and TRIM21, TRIM72 belongs to TRIM C-IV subfamily and consists of multiple domains (Hatakeyama, 2017; Ozato et al., 2008). A flexible RING domain confers E3 ligase activity (Nguyen et al., 2014; Song, 2013) and a conserved B-box domain is required for stress-induced oligomerization (Ma et al., 2023). A coiled-coil domain contains leucine zipper motif 1 (LZ1) which is necessary for dimerization (Hwang et al., 2011) and mediates intracellular vesicle tethering (Ma et al., 2023). PRY/SPRY domain at the C-terminus is involved in calcium regulation, lipid headgroup binding, and vesicle-vesicle and vesicle-plasma membrane fusion (Ahn, 2016; Ma et al., 2023).

Plasma membrane integrity is crucial to maintaining cellular homeostasis and functions (Ammendolia et al., 2021; McNeil and Kirchhausen, 2005). When subjected to cellular stress and membrane damage, cells preserve the integrity of the plasma membrane by activating intrinsic pathways to promote the recycling of lipid membrane components and facilitate repair. Deficits in membrane repair responses have been associated with muscle dystrophy and heart failure, which are diseases commonly associated with the defective membrane repair of muscle cells (Ammendolia et al., 2021; Bansal and Campbell, 2004; Doherty and McNally, 2003; McNeil and Kirchhausen, 2005; Zhong et al., 2021). However, whether membrane repair deficits contribute to the pathogenesis of ALS and other neurodegenerative diseases remains elusive. Notably, TRIM72 is activated when plasma membrane integrity is compromised and initiates a coordinated series of events to promote membrane repair (Cai et al., 2009). However, the mechanisms underlying TRIM72-mediated membrane repair and vesicle recruitment are still largely unknown.

The mammalian Retriever complex plays a critical role in endosomal trafficking, membrane trafficking and recycling. It is functionally similar to Retromer in mediating endosome-to-Golgi and endosome-to-plasma membrane receptor trafficking and endosomal recycling of integral membrane proteins (Chen et al., 2019; McNally et al., 2017; Wang et al., 2018b). Retriever consists of C16orf62 (VPS35L), DSCR3 (VPS26C), and VPS29 and interacts with the CCC (COMMD/CCDC22/CCDC93) complex to assemble into the Commander complex, a multi-protein complex that plays essential roles in endosomal trafficking similar to the Retromer cargo recognition complex (Chen et al., 2019; Healy et al., 2023; Mallam and Marcotte, 2017; McNally et al., 2017; Wang et al., 2018b). The CCC complex consists of 12 subunits, including 10 COMMD (copper metabolism MURR1 domain) proteins, from COMMD1 to COMMD10, and two CCDC (coiled-coil domain-containing) proteins, CCDC22 and CCDC93 (Healy et al., 2023). The COMMD proteins form a decametric ring structure that interacts with CCDC22 and CCDC93 proteins which stabilize the COMMD ring structure and serve as a bridge to link the COMMD ring structure to Retriever (Healy et al., 2023). In addition, deficits in Retromer-associated retrograde trafficking have been previously associated with neurodegenerative diseases, such as AD and PD (Gershlick and Lucas, 2017; Hu et al., 2015; Schreij et al., 2016). Given the functional and structural similarities between Retromer and Retriever, whether the Retriever plays a role in neurodegenerative diseases remains to be explored.

Here, we discovered that loss-of-function of TRIM72 significantly accelerates the ALS-like phenotypes in FUS-R521C KI mice we generated. In addition, loss-of-function of TRIM72 induced the mislocalization of mutant FUS in the FUS-R521C KI mouse spinal cord, which is a hallmark, like mislocalized TDP-43, shown in patients in ALS and other degenerative diseases. Furthermore, we have demonstrated that Retriever/Commander complex promotes membrane repair function mediated by TRIM72. We disclosed that the Coiled-coil and the PRY/SPRY domains of TRIM72 are indispensable for TRIM72-mediated membrane repair and anti-oxidation functions. Notably, the membrane repair functions of TRIM72 are independent of its RING domain responsible for E3 ligase activity. Application of AAV-based neuronal expression of TRIM72 in ALS models with SOD1 and TDP-43 mutations promotes animal survival, suggesting a pan neuronal protection in ALS. An ALS patient with R521C mutation received a dose of AAV9-TRIM72 displayed slowdown of disease progression in an Investigator Initiated Trial (IIT) study. Therefore, a better understanding of the functions of TRIM family members and the roles of the Retriever/Commander complex in the disease progression may unveil therapeutic targets for ALS and other neurodegenerative diseases.

## RESULTS

### Neuronal upregulation of TRIM72 in FUS-R521C knockin rodent ALS models

To study the disease mechanisms underlying neurodegeneration caused by the mutant FUS in ALS, we isolated spinal cords from wildtype and FUS-R521C KI mice at two age points, 1.5 and 7 months, and applied for RNA-seq. Compared to wildtype controls, only a few genes were differentially expressed in the mutant FUS spinal cords. Among them, *Trim72* was the only differentially expressed gene at both time points, with about 8-10 times upregulation in the mutant spinal cord (Figure 1A). To examine the upregulation of *Trim72* in motor neurons, we cultured the motor neurons from wildtype and FUS-R521C mutant animals and confirmed the motor neuron upregulation (Figure 1B).

**Figure 1.**
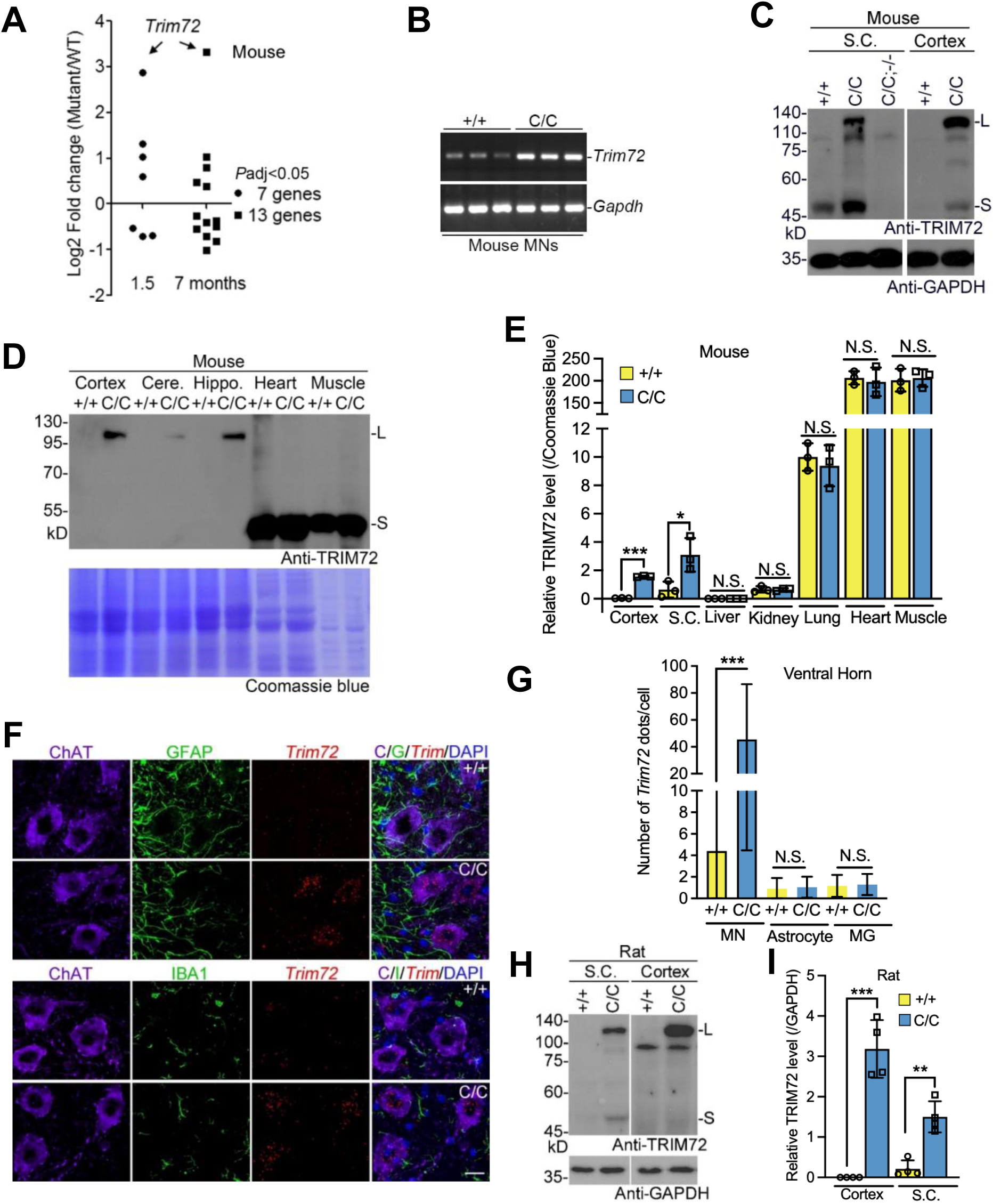
Neuronal upregulation of TRIM72 in FUS-R521C knockin rodent ALS models. (A) RNA-sequencing identified differentially expressed genes detected in the spinal cords at 1.5 and 7 months of age in wildtype (+/+) and FUS-R521C (C/C) mice (*P*adj < 0.05, n = 3). (B) *Trim72* was upregulated in C/C mice but not in +/+ mice-derived motor neurons (MNs, 3 DIV). RT-PCR, 3 biological replicates, *Gapdh* as loading control. (C) TRIM72 was upregulated in the spinal cord (S.C.) and cortex of C/C mutant mouse but not in the S.C. and cortex of +/+ mouse. GAPDH was used as loading control. *Trim72* knockout mouse with FUS-R521C mutation (C/C;−/−) served as a negative control. (**D** and **E**) TRIM72 was upregulated in neural but not non-neural tissues in mice with the indicated genotypes. Coomassie blue staining served as a loading control. Cere., cerebellum; Hippo., hippocampus. Data summary was shown in **E**. (**F** and **G**) *Trim72* RNA *in situ* hybridization was performed in the spinal cord ventral horns with the indicated genotypes. In the spinal cord, motor neurons (MN), astrocytes, and microglial (MG) were marked by ChAT, GFAP, and Iba1, respectively. In **F**, scale bar, 20 μm. Mice, 3 months of age (n = 3). Data summaries were shown in **G**. (**H** and **I**) TRIM72 expression was documented in wildtype (+/+) and FUS-R521C (C/C) knockin mutant rats (PMID: 30273830; 30338063). GAPDH was used as loading control. Data summary was shown in **I**. In **E**, **G**, and **I**, values were presented as mean ± SD. \**p* <0.05; \*\*\**p* <0.001; N.S., no statistical significance (T-Test, SPSS).

To study whether the upregulation of *Trim72* is involved in the disease pathogenesis, we generated *Trim72*^−/−^ lines in both wildtype and FUS-R521C mutant backgrounds, because of a close chromosomal distance between *Trim72* and *Fus* (Figure S1). Indeed, a significant upregulation of TRIM72 in the mutant neural tissues, including spinal cord, motor cortex, cerebellum, and hippocampus, was evidenced by western blot in comparison with that of wildtype (Figure 1C-1E). However, the TRIM72 band was absent in the *Trim72* knockout animals (Figure 1C and Figure S2A), and TRIM72 expression was not detected in livers in both wildtype and FUS-R521C mutant mice (Figure S2A. Although TRIM72 expressions vary a lot in non-neural tissues we examined, comparable expressions between wildtype and FUS-R521C mutant animals in those non-neural tissues, including heart, muscle, lung, and kidney we examined (Figure 1D, 1E, and S2). We noticed that both small (∼55kD) and large (∼110kD) molecular weight TRIM72 appeared in the FUS-R521C mutant neural tissues (Figure 1C and 1D). However, the ∼110kD band was not detected in those neural tissues in wild type animals as well as in non-neural tissues in both genotypes, suggesting that the appearance of high molecular weight TRIM72 in the mutant animals is neural tissue-specific and dependent on the FUS mutation. Unlike homo-oligomers of TRIM72 shown in our muscle tissue lysate (a lysate condition without DDT) that was sensitive to as low as 10 mM DTT (Figure S3A, the ∼110kD band in the mutant neural tissue is insensitive to DTT, as well as urea and β-ME (2-mercaptoethanol), suggesting that the formation of large TRIM72 complex or dimer does not depend on disulfide bonds (Figure S3B. We also estimated the amount of TRIM72 in the mutant spinal cord, which is as much as 1.5% of that shown in muscle (607.07±107.08 ng/mg muscle, mean±SD).

To further characterize the upregulation of *Trim72* in which cell types in the mutant neural tissues, we performed *Trim72* RNA *in situ* hybridization (ISH) in the mutant spinal cord and motor cortex. Ten-time more *Trim72* dots were detected by ISH in the ChAT-positive motor neurons in the mutant spinal cord than that of wildtype (Figurer 1F and 1G). However, such upregulation is not present in the mutant GFAP-positive astrocytes and IBA1-positive microglia in the mutant spinal cord. A significant upregulation of *Trim72* was also evidenced in *Vglut1*-positive excitatory and *Vgat*-positive inhibitory neurons but not in astrocytes and microglia in the mutant motor cortex (Figure S4). A previously reported FUS-R521C knockin rat ALS model, generation of which was independent from ours, showed no obvious ALS-like phenotypes (Zhang et al., 2018), like our FUS-R521C knockin mouse. However, a significant upregulation of TRIM72 was detected in neural but not non-neural tissues in those FUS-R521C knockin rats (Figure 1H, 1I, and S2B). In addition, we also noticed that large molecular weight TRIM72 appeared in the neural but not non-neural tissues in the rats. Our data also supported that both small and large molecular weight TRIM72 are upregulated in the mutant neuronal tissues (Figure S5). Taken together, we demonstrated that neuronal upregulation of TRIM72 appears in two FUS-R521C knockin rodent ALS models, independently.

### Loss-of-function of *Trim72* induces ALS-like phenotypes

To examine whether the neuronal upregulation of TRIM72 modifies motor functions in our FUS-R521C mouse model, we analyzed the motor performance in wild-type (+/+), FUS-R521C (C/C), FUS-R521C with *Trim72* knockout (C/C;−/−), and *Trim72* knockout alone (−/−) animals. A more severe decline of motor ability was observed in the 4-month-old C/C;−/− animals compared to that of the other three genotypes in the rotarod performance (Figure 2A). Electromyography (EMG) of hind limb supported muscle weakness shown in C/C;−/− mice with significantly increased amplitude and duration, compared to the other three genotypes (Figure 2B-2D). In agreement with impaired motor performance and function, a significant motor neuron loss was observed in C/C;- /- animals compared to that of the other three genotypes at one year of age (Figure 2E and 2F). In contrast, the *Trim72* knockout mice did not display motor neuron loss compared to +/+ controls (Figure 2F). The survival curve of these four genotypes agreed that C/C;−/− animals have a shorter lifespan (∼40% of C/C;−/− animals died at one year of age) compared to that of the other three genotypes (Figure 2G). The rest of aged C/C;−/− animals appeared paralysis, abnormal gait performance, and hindlimb pullback, which we had never observed in +/+, C/C, and −/− mice (Figure 2H, 2I, S6, and Movie S1). Therefore, we concluded that loss-of-function of *Trim72* accelerates ALS-like disease progression shown in the FUS-R521C knockin ALS model.

**Figure 2.**
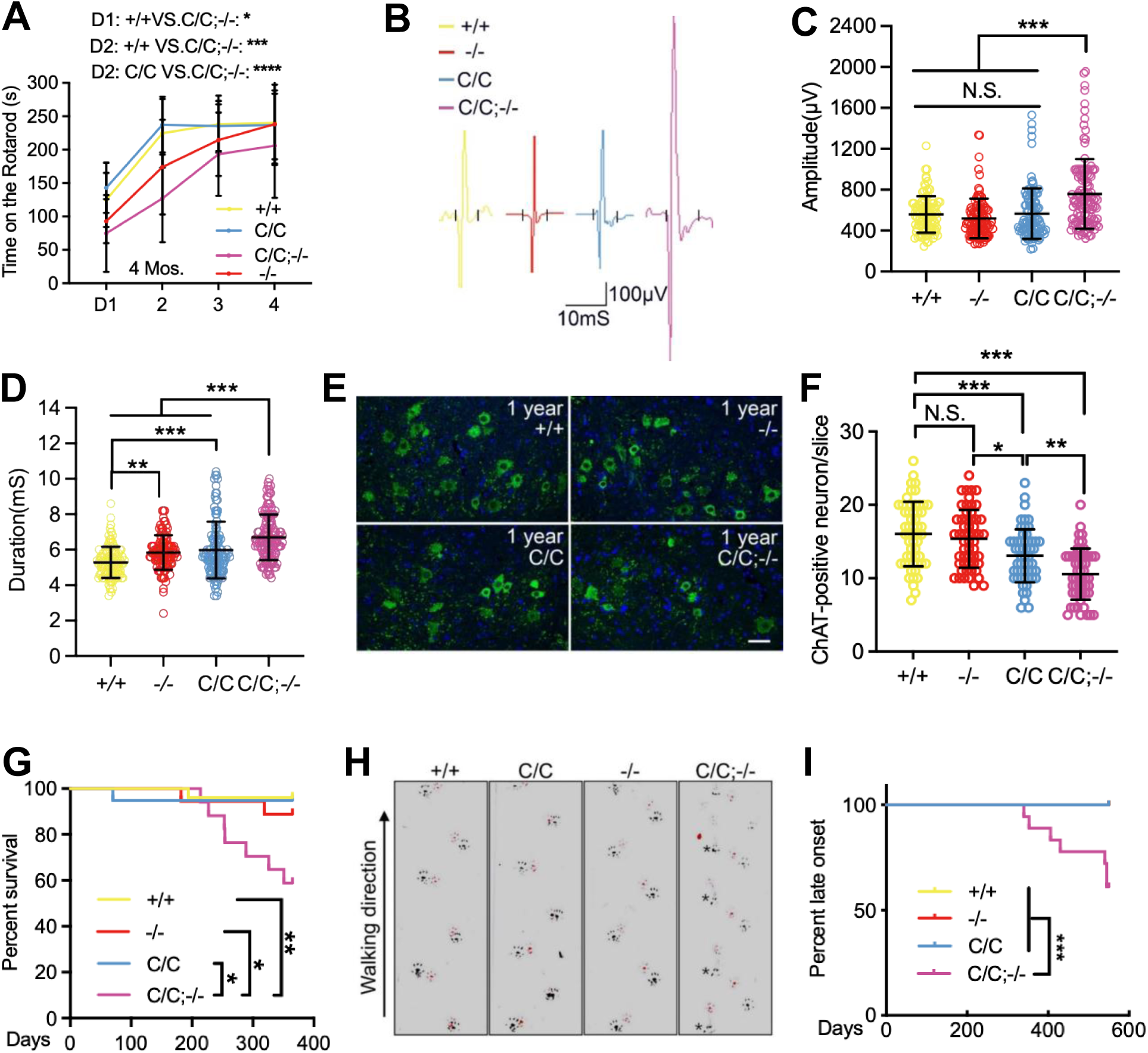
*Trim72* loss-of-function accelerates disease progress shown in FUS-R521C ALS mouse model. (**A**) Rotarod test was performed at 4 months of age. Mice, male, n = 10 (+/+), 10 (C/C), 12 (C/C;−/−), and 9 (−/−). (**B**-**D**) Compound muscle action potentials were recorded from hind limb of indicated genotypes. Amplitude and duration were summarized in **C** and **D**, respectively. Mice, 14-month-old males, n ≥ 5. (**E** and **F**) Number of ChAT-positive motor neurons in the ventral horns at 1 year of age (n ≥ 3). ChAT-positive motor neurons in cross-sections of lumbar 4-5 spinal cords. Scale bar in **E**, 50 μm. Data summary was shown in **F**. (**G**) Survival curve of indicated genotypes. *Trim72* loss-of-function significantly shortened the life span of FUS-R521C knockin mice. Mice, male, n = 25 (+/+), 19 (C/C), 17 (C/C;−/−), and 18 (−/−). (**H** and **I**) Representative gait pattern of the indicated genotypes (**H**). Mice, male, 1.5 year of age. Percentage of abnormal gait onset **(I**). Mice, male, n = 16 (+/+), 15 (C/C), 18 (C/C;−/−), and 15 (−/−). In **A**, **C**, **D**, and **F**, values were presented as mean ± SD. \**p* <0.05, \*\**p* <0.01, \*\*\**p* <0.001, and \*\*\*\**p* <0.0001 (ANOVA, SPSS). N.S., no statistical significance. In **G** and **I**, statistic analysis, log-rank test.

### Loss-of-function of *Trim72* does not affect FUS-R521C protein level but increases oxidative stress in FUS-R521C knockin ALS model

As an E3 ligase, TRIM72 may target mutant FUS for degradation; therefore, loss-of-function of *Trim72* would result in mutant FUS protein accumulation to accelerate the disease progression. To test it, we analyzed the FUS protein level in the spinal cord in these mutant mice (Figure 3A). Although upregulation of mutant FUS was documented in C/C and C/C;−/− spinal cords, we did not observe mutant or wildtype FUS protein level change when *Trim72* was knocked out.

**Figure 3.**
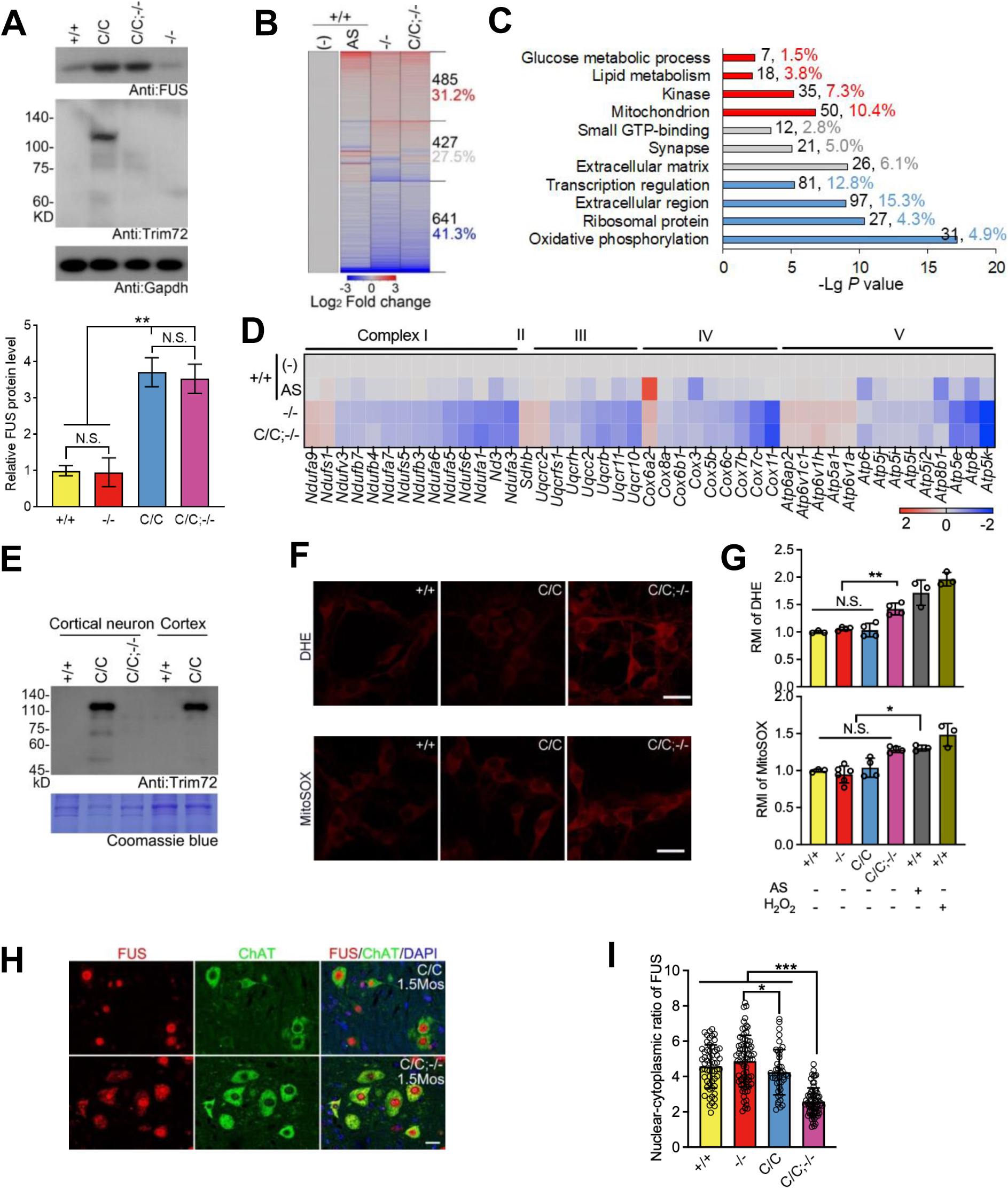
Loss-of-function of *Trim72* induces oxidative stress and mislocalization of mutant FUS in spinal cord motor neuron. (A) *Trim72*_−/−_ did not change the expression level of wildtype and mutant FUS. The data summary shown in (lower panel). GAPDH served as loading control. (B) The spinal cord mRNAs from wildtype (+/+), wildtype treated with AS for 3 hours, *Trim72*_−/−_, and homozygous FUS-R521C and *Trim72* knock-out (C/C;−/−) mice were applied for RNA-seq. Differentially expressed genes (DEGs) were identified by a multi-group comparison (*P*adj < 0.05). AS treatment, intragastric administration, 6 μg/g body weight. Mice, male, 1 month of age, n = 3. (C) DAVID functional analysis of DEGs in the indicated genotypes shown in (**B**). Red: upregulated genes; blue, downregulated genes; gray, the rest. (D) DEGs encode mitochondria complexes I, II, III, IV and V. (E) Upregulation of TRIM72 in FUS-R521C cultured cortical neurons (14DIVs) but not in that of +/+ and C/C;−/−. Coomassie blue staining served as a loading control. (**F** and **G**) DHE and MitoSOX staining in the cultured cortical neurons (12DIVs) with the indicated genotypes (**F**). Neurons treated with AS (50 mM, 30 mins) and H_2_O_2_ (100 μm, 30 minutes) were served as positive controls for the staining. The data summary shown in **G**. RMI, relative mean intensity. (**H** and **I**) Mislocalization of mutant FUS was documented in the ChAT-positive motor neuron in C/C;−/− mice (**H**). Mean FUS fluorescent intensity in the nucleus and cytoplasm were measured and summarized in (**I**). Sections of lumbar 4-5 spinal cords. Scale bar, 20 μm. In **A**, **G**, and **I**, values were presented as mean ± SD; in **A**, n = 3; in **G**, n = 3-4; in **I**, n = 4. N.S., no statistical significance; \**p* < 0.05; \*\**p* < 0.01; \*\*\**p* < 0.001; ANOVA, SPSS.

To study the mechanisms underlying acceleration of the disease progression caused by loss-of-function of *Trim72*, we extracted mRNAs from +/+, −/−, and C/C;−/− spinal cords at one month of age and sent them for RNA-seq (Figure 3B). Only small amount of differentially expressed genes (DEGs, 86 upregulated and 53 downregulated genes) appeared in C/C;−/− spinal cords compared to that of −/− (Figure S7), similar to what we observed in the comparison between C/C and +/+ animals (Figure 1A). However, over a thousand genes were differentially expressed among +/+, −/−, and C/C;−/− groups (Figure 3B). Functional GO analysis revealed those DEGs either encode mitochondrion proteins or are functionally involved in oxidative phosphorylation (Figure 3C), suggesting that loss-of-function of *Trim72* damages mitochondrion functions in the mutant spinal cord. To test our hypothesis, we treated +/+ mice with arsenite (AS), an oxidative stress inducer, and sent their spinal cords for RNA-seq (Figure 3B). Interestingly, the spinal cord DEGs from +/+ treated with AS, −/−, and C/C;−/− mice had ∼72.5% overlap (Figure 3B), suggesting that loss-of-function of *Trim72* generates oxidative stress in the mutant spinal cord. Indeed, the similar expression patterns of DEGs encoding mitochondrion complex I, II, III, IV and V were found in spinal cords among these three genotypes (Figure 3D).

To confirm oxidative stress induced by loss-of-function of *Trim72*, we cultured cortical neurons from FUS-R521C KI mice and observed TRIM72 upregulation, which was undetectable in both +/+ and C/C;−/− cultures (Figure 3E), similar to what we observed in the mutant motor cortex (Figure 1D and 1E). We then employed dihydroethidium (DHE), a superoxide indicator (Saiki et al., 1986), to measure the level of reactive oxygen species (ROS) in these primary neuron cultures. As positive controls, wildtype cortical neuron cultures treated with AS or H_2_O_2_ displayed significantly higher RMI (relative mean intensity) of DHE staining compared to that of the naïve control group (Figure 3F and 3G). Although the RMI values were comparable among +/+, −/−, and C/C cultured neurons, the values were significantly higher in cortical neurons cultured from C/C;−/− animals (Figure 3F and 3G), indicating that loss-of-function of *Trim72* increases ROS level in these FUS-R521C mutant neurons. The loss-of-function of *Trim72*-induced oxidative stress in the FUS-R521C mutant neurons was further confirmed by MitoSOX, a mitochondrial superoxide indicator (Robinson et al., 2006) (Figure 3F and 3G).

Mislocalized TDP-43 or FUS from nucleus to cytoplasm is a hallmark of ALS, and chronic oxidative stress (AS treatments) induced the mislocalization of R521C mutant but not the wildtype FUS in the spinal cord motor neurons (Zhang et al., 2020). In addition, the chronic oxidative stress induced upregulation of stress granule proteins, including G3BP1/2 and EIF3η, in the mutant spinal cord (Zhang et al., 2020). If loss-of-function of *Trim72* increases oxidative stress *in vivo* (Figure 3B-3D), we speculated that loss-of-function of *Trim72* leads to mislocalization of mutant FUS and upregulation of stress granule proteins in the C/C;−/− spinal cord. Indeed, without treatment of chronic stress, we observed significantly more mislocalized mutant FUS in ChAT-positive motor neurons in the C/C;−/− spinal cords, compared to that of +/+, −/−, and C/C mice (Figure 3H and 3I, Figure S8). In addition, upregulation of SG proteins, including G3BP2 and eIF3η, were documented in the C/C;−/− spinal cord, compared to that of C/C mice (Figure S9A. Similar upregulation of SG proteins was also evidenced in the C/C;−/− motor cortex (Figure S9B and S9C).

If loss-of-function of *Trim72* increases oxidative stress in the FUS mutant CNS, we speculated that C/C;−/− mouse would more sensitize to oxidative stress challenge, like AS treatment, which previously we used to challenge the FUS-R521C mutant animals (Zhang et al., 2020). To this end, we challenged +/+, C/C, C/C;−/−, and −/− animals with 4 doses of AS for a week and examined the treatment on their survival in 15 days (Figure S10). Compared to small percentage of animal death in +/+, C/C, and −/− groups during our experimental procedure, we observed significantly more animal death in the C/C;−/− group at the end of our 15-day observation. Taken together, we concluded that loss-of-function of *Trim72* significantly increases oxidation in FUS-R521C mutant CNS and renders the mutant mouse more sensitive to oxidative stress challenge.

### Loss-of-function of *Trim72* worsens axonal mutant FUS condensates that in turn impairs axonal mitochondrion dynamics

To study how loss-of-function of *Trim72* induces oxidation in the mutant neurons, we examined association of mutant FUS with mitochondria in the C/C;−/− spinal cord (Figure S11A. Cytosol mislocalized mutant FUS was seen in C/C;−/− motor neuron, which was not associated with mitochondria evidenced by bare localization between mutant FUS and ATP5A, a mitochondrial marker. In mitochondrial fractionation separated from C/C cortices, we failed to demonstrate association between TRIM72 and mitochondria (Figure S11B. Instead, in cytosol fractionation, we did see TRIM72 appearance.

We then examined the impact of *Trim72* depletion on axonal trafficking of mitochondria. Specifically, we assessed mitochondrial mobility in axons of cortical neurons derived from +/+, C/C, and C/C;−/− mice. Using a microfluidic device-based assay (Figure S12) (Wang et al., 2016), we successfully labeled axonal mitochondria in cortical neurons from +/+, C/C, and C/C;−/− mice (Figure S13A. Employing an automatic tracking method, we observed a reduction in axonal mitochondrial trafficking speed in C/C;−/− mice, compared to +/+ and C/C counterparts (Figure S13B. To gain deeper insights into mitochondrial mobility, we classified these mitochondrial tracks into stationary and mobile fractions, based on the average speeds of trajectories. Consistently, we observed a significant increase in the proportion of stationary mitochondria from 36.65% ± 13.57% in C/C neurons to 47.45% ± 8.538% in C/C;−/− neurons, accompanied by a significant reduction in the mobile fraction, compared to +/+ and C/C counterparts (Figure S13C. These findings indicate that C/C;−/− cortical neurons exhibit hindered axonal mitochondrial transport compared to that of +/+ and C/C, emphasizing the impeded trafficking dynamics caused by loss-of-function of *Trim72* in presence of mutant FUS.

To study how loss-of-function of *Trim72* impedes mitochondrial mobility in FUS mutant cortical neurons, we measured immunoreactive FUS signals in the axons of cortical neurons derived from +/+, C/C, and C/C;−/− mice (Figure S13D. Compared to even distribution of immunoreactive FUS signals shown in +/+ and C/C axons, large and round FUS-positive puncta were formed in C/C;−/− axons, resembling FUS condensates observed under stress conditions (Zhang et al., 2020). To quantify these FUS-positive condensates, we employed an automatic feature extraction method based on the size and shape and found a significantly increased number of FUS-positive condensates in C/C;−/− axons compared to those of +/+ and C/C (Figure S13E. Additionally, the size of these FUS-positive condensates in C/C;−/− axons were significantly increased compared to that of +/+. Similar size increase was seen between C/C;−/− and C/C groups although with no significant difference (Figure S13F. Taken together, these data indicate that loss-of-function of *Trim72* significantly induces the formation of enlarged mutant FUS condensates in the axons, which probably in turn impairs axonal mitochondrion dynamics.

### A pan neuronal protective effect of TRIM72 in ALS mouse models

To examine whether expression of TRIM72 is sufficient to protect cell from oxidative stress, we stably expressed TRIM72 with lentivirus in Hela cells, in which we barely detected TRIM72 expression as well as in commonly used U2Os and 293FT cells (Figure S14A. To induce oxidative stress in these cells, we treated the cells expressing TRIM72 with various concentrations of AS and then measured cell dehydrogenase activity by CCK-8 assay within 2 hours. Because our CCK-8 measurement took place within 2 hours after AS treatment, it limits the effects of cell proliferation on the measurement. To avoid batch to batch variation of cell number and status between our individual experiment and accurately reflect cell dehydrogenase activity, in a batch of experiments we included both AS-treated and untreated groups and normalized CCK-8 values of AS-treated to that of untreated ones (Figure 4A). Indeed, with the increase of AS concentration, the CCK-8 values of AS-treated cells without expression of TRIM72 were reduced in a dosage-dependent manner (Figure 4B). However, no matter how the AS concentration increased, the expression of TRIM72 significantly protected the cells from oxidative stress compared to that of control cells treated with the same concentration of AS. The protective effect of TRIM72 from oxidative stress was further confirmed in the C/C;−/− cultured neurons expressing TRIM72 by lentivirus (Figure 4C, Figure S14B and S14C). Therefore, we concluded that expression of TRIM72 is sufficient to protect cell from oxidative stress.

**Figure 4.**
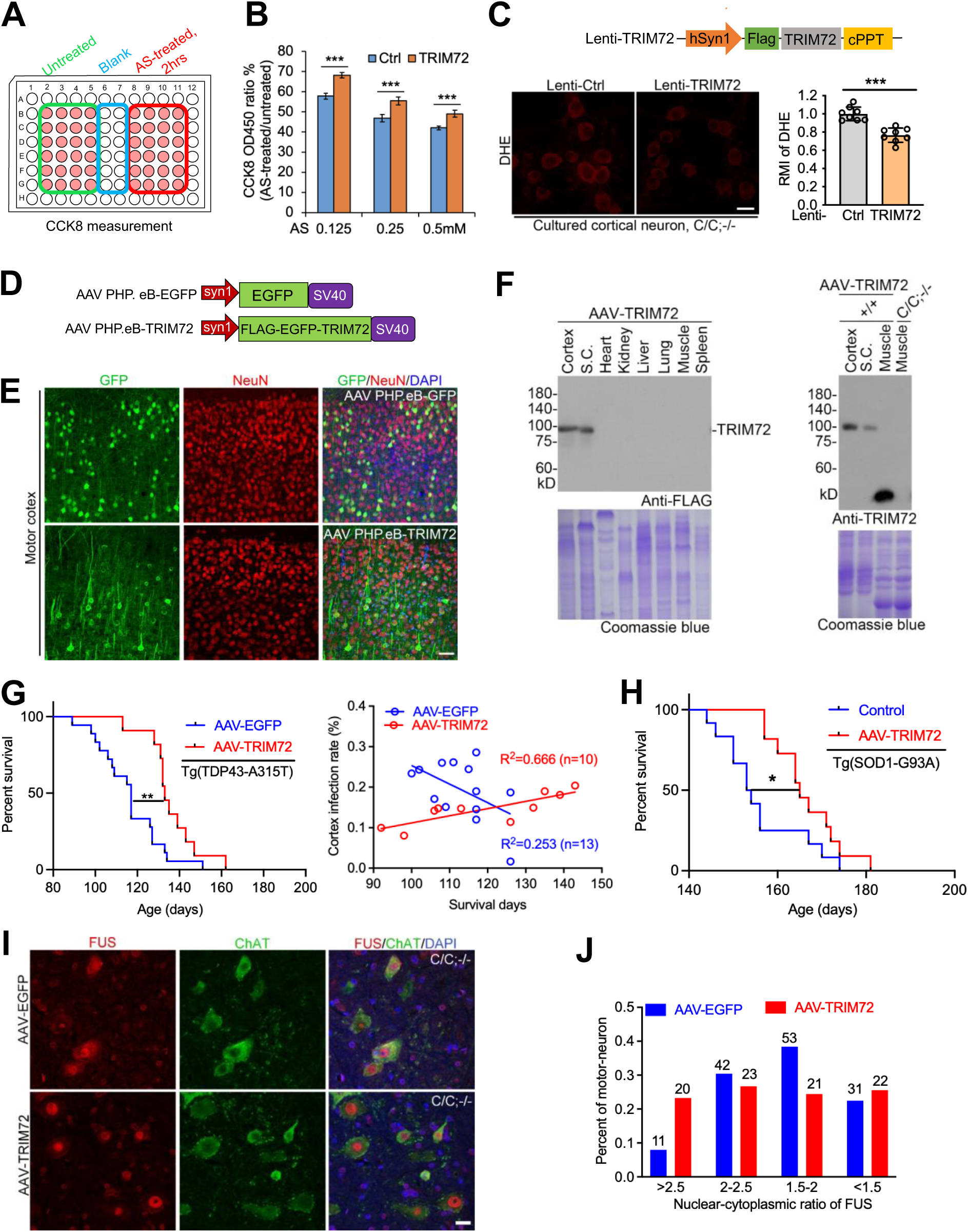
TRIM72 expression protects cells from oxidative stress and extends the life span of ALS mouse models. (**A** and **B**) Hela cells were stably expressed with and without TRIM72 by lentivirus and subsequently treated with AS at the indicated concentrations for 2 hours. After AS treatment, cell dehydrogenase activity was measured by CCK-8 assay within 2 hours. Ctrl, empty vector. Experimental procedure was illustrated I (**A**). Data summary was shown in (**B**). (A) DHE staining in cultured cortical neurons (12DIVs) with and without TRIM72 expression. Lentiviral expression cassette was showed in the upper panel. Lenti-Ctrl, empty lentiviral particles. Data summary was shown in bottom right. (B) AAV (AAV PHP.eB) design of TRIM72 expression in CNS. TRIM72 was tagged with FLAG and EGFP at the N-terminus, and expressed under a Syn1 neuron-specific promoter. AAVs were retro-orbitally injected into adult mice (1 × E11 vg/mouse). AAV PHP.eB-EGFP served as control. (C) Cortical expression of TRIM72 by AAVs. GFP fluorescence was assessed 1 month after injection. NeuN, neuron marker. EGFP-TRIM72 showed cytosol and neurite localization in infected cortical neurons. (D) TRIM72 was expressed in neural, but not non-neural tissues, detected by immunoblot 1 month post-AAV injection. Coomassie blue staining served as a loading control. AAV-TRIM72, AAV PHP.eB-TRIM72. (E) Survival curve of Tg (TDP-43-A315T) mice infected with AAV-TRIM72 or AAV-EGFP (Left). Cortical infection rate of AAV-TRIM72, but not AAV-EGFP, was positively correlated with survival days of the given individuals (Right). For survival curve, mice, AAV-TRIM72 (n = 11), AAV-EGFP (n = 14). For correlation between infection rates and survival days, mice, AAV-TRIM72 (n = 10), AAV-EGFP (n = 13). AAV-EGFP, AAV PHP.eB-EGFP. (F) Survival curve of Tg (SOD1-G93A) mice infected with AAV-TRIM72 (n = 11). Ctrl, saline-treated mice (n = 12). (**I** and **J**) Mislocalization of mutant FUS in C/C;−/− ChAT-positive motor neuron was restored by TRIM72 expression (**I**). Mean FUS fluorescent intensities in the nucleus and cytoplasm were measured and summarized in (**J**). In **I**, sections of lumbar 4-5 spinal cords (mouse n = 3). Scale bar, 20 μm. In **B** and **C**, values were presented as mean ± SD; in **B**, n = 8-10; in **C**, n = 8; in **J,** n = 3. N.S., no statistical significance; \**p* < 0.01; \*\**p* < 0.01; \*\*\**p* < 0.001; T-test, SPSS. In **G** and **H**, log-rank test.

To examine neuronal protective effect of TRIM72 *in vivo*, we employed a neuronal promoter, Syn1, to drive the expression of TRIM72 and an AAV serotype, AAV-PHP.eB, which has been shown with a high neuronal infection rate in mouse brain by retro-orbital injection (Chan et al., 2017) (Figure 4D). Indeed, neuronal expression of GFP-tagged TRIM72 was documented one month after the AAV injection (1 × E11 vg/mouse) in the adult mouse motor cortex and spinal cord with a cytosol localization, compared to a diffused expression of GFP control (Figure 4E and Figure S15). The neural specificity of TRIM72 expression was further documented in motor cortex and spinal cord but not non-neural tissues by western blot with both Flag- and TRIM72-antibodies in the AAV-TRIM72-infected animals (Figure 4F). We then injected AAV-GFP and AAV-TRIM72 in a transgenic ALS (TDP-43-A315T) mouse model (Wegorzewska et al., 2009) at two months of age, respectively. Injection of AAV-TRIM72 significantly increased the lifespans of these TDP-43-A315T mice, compared to that of AAV-GFP injection control (Figure 4G). In addition, the mutant animals with higher AAV-TRIM72 infection rate in motor cortex displayed longer lifespans (R^2^ = 0.666, n = 10), which was not seen in that of AAV-GFP controls (R^2^ = 0.253, n = 13) (Figure 4G). Expression of TRIM72 by AAV-PHP.eB also significantly extended lifespans of SOD1-G93A mice, a conventional ALS mouse model (Gurney et al., 1994) (Figure 4H). In addition, the expression of TRIM72 by AAV-PHP.eB recovered cytosolic mislocalized mutant FUS in the C/C;−/− spinal cord (Figure 4I and 4J), one of the key pathological features shown in C/C;−/− mouse (Figure 3E and 3F). Taken together, we concluded that neuronal expression of TRIM72 extends lifespan of two ALS mouse models, and restore mutant FUS mislocalization shown in FUS-R521C mutant knockin spinal cords, indicating a pan neuronal protective effect of TRIM72 in two different ALS mouse models.

### Monomer and dimmer of TRIM72 interact with CCC complex

To tackle the underlying mechanism of TRIM72-mediated protection, we stably expressed Flag-TRIM72 in Hela cells and employed Flag-IP/MS to identify TRIM72 interacting partners (Figure S16), which we previously employed for a protein functional study (Meng et al., 2023). Except for TRIM72, the top hits of our IP/MS belong to a protein complex, COMMD/CCDC22/CCDC93 (CCC) complex (Figure S16 and Table 1), which together with a trimeric protein complex Retriever (comprising VPS35L, VPS26C, and VPS29) are responsible for recycling of membrane proteins from endosome to the cell surface (Healy et al., 2023). The interaction between TRIM72 and CCDC22 was validated in 293FT cells by expressions of N-terminal Flag-tagged TRIM72 (Flag-TRIM72) and C-terminal Myc-tagged CCDC22 (CCDC22-Myc), which is independent of oxidation induced by AS treatment (Figure 5A).

**Figure 5.**
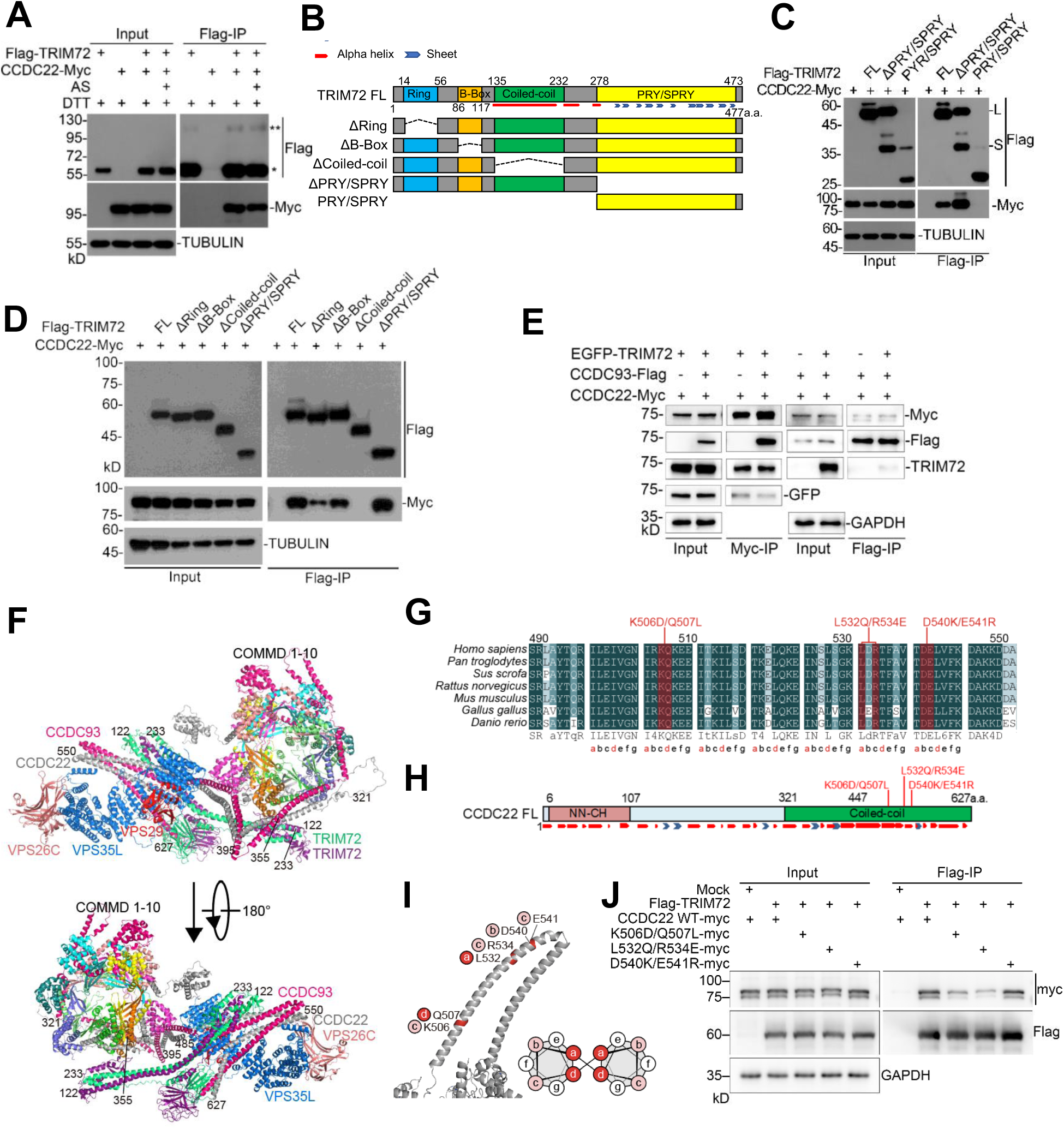
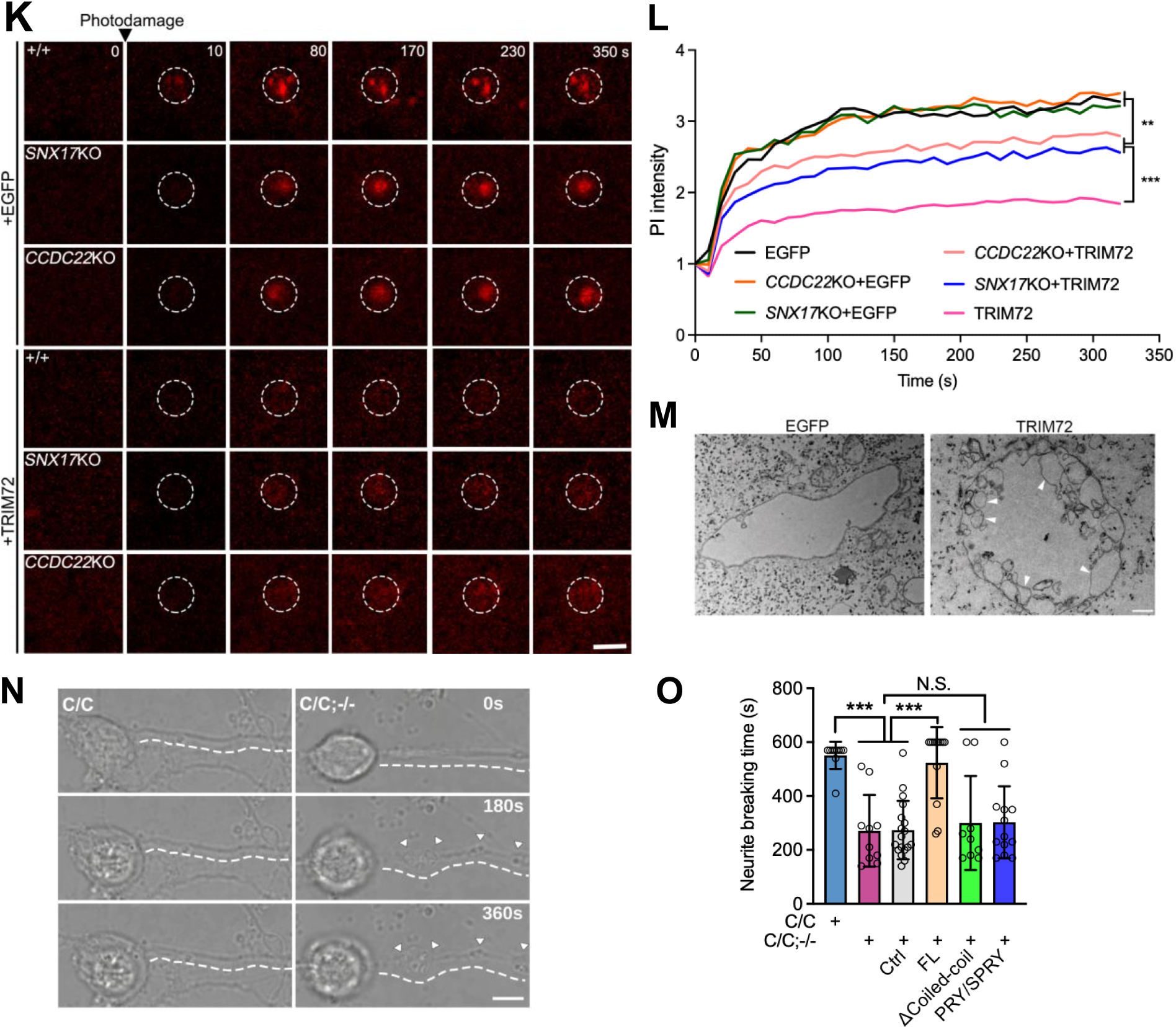
CCC and Retriever protein components are required for TRIM72-mediated anti-oxidation and membrane repair. (A) Flag-TRIM72 interacted with CCDC22-Myc in 293FT cells with or without AS treatment. TRIM72 large molecular weight band (**) was insensitive to DTT. (B) Protein domain and secondary structure features of TRIM72. FL, full length. (C) PRY/SPRY domain of TRIM72 is not required for its interaction with CCDC22. Note a strong large molecular weight band (L) appeared in the Input and Flag-IP in ΔPRY/SPRY group, which was insensitive to DTT, urea, and β-ME (**Figure S17**). (D) TRIM72 interacted with the coiled-coil domain of CCDC22. (E) EGFP-TRIM72, CCDC93-Flag, and/or CCDC22-Myc were expressed in 293FT cells. TRIM72 expression does not disrupt CCDC22/CCDC93 interaction. (F) Protein complex of TRIM72 and Commander illustrated with AlphaFold-3. CCDC22 interaction with CCDC93 was independent from CCDC22 interaction with TRIM72 (also see the **Movie S5**). (G) Conservation analysis of CCDC22 480-627a.a. The module (abcdefg) of heptad repeat (a seven-residue periodicity) of CCDC22 Coiled-Coil domain was labeled. (**H** and **I**) The potential key residues crucial for interaction between CCDC22 and TRIM72 in primary (**H**) and secondary structure (**I**) of CCDC22. (**J**) K506D/Q507L and L532Q/R534E mutations affect the interaction between CCDC22 and TRIM72. (**K** and **L**) EGFP or EGFP-TRIM72 was expressed in U2OS cells with target gene KO by lentivirus (**Figure S20**) and cells were photodamaged and monitored by time-lapse imaging. Circles indicate the damaged sites. Normalized PI intensities were summarized (**L**), which represented damage severity. (**M**) Vesicles (arrowhead labeled) are evident by CLEM in U2OS cells expressing EGFP-TRIM72, but not in those expressing EGFP alone, following membrane damage. Cells were subjected to photodamage and then fixed 10 minutes after the damage. The resulting cells were sectioned and subsequently observed using an electron microscope. Scale bar, 500 nm. (**N** and **O**) Cultured cortical neurons with the indicated genotypes were photodamaged at soma and monitored by time-lapse imaging (**N**). Note that broken neurites were observed in C/C;−/− but not C/C neuron as early as 170s after photodamage. Data summary of the neurite breaking time (**O**). In **L,** and **O**, values were presented as mean ± SD; in **L**, n = 12∼17; in **O**, n = 9∼18;. N.S., no statistical significance, \**p* < 0.01, \*\**p* < 0.01, \*\*\**p* < 0.001,one-way ANOVA, SPSS. Scale bar, in **K**, 2 µm; in **N**, 10 µm.

**Table 1.** Identification of TRIM72 interacting partners by IP/MS.

| Name | Flag-Trim72 |  |  | Control |  |  | Functions | PMID |
| --- | --- | --- | --- | --- | --- | --- | --- | --- |
|  | 1 | 2 | 3 | 1 | 2 | 3 |  |  |
| TRIM72 | 799.4 | 2084.1 | 2433.2 | 25.8 | 0.0 | 0.0 | Membrane repair | 19043407 |
| CCDC22 | 55.4 | 131.4 | 123.7 | 0.0 | 0.0 | 0.0 | Membrane protein trafficking | 26965651;25355947 |
| COMMD3 | 15.5 | 33.6 | 35.6 | 0.0 | 0.0 | 0.0 | Membrane protein trafficking | 26965651;25355947 |
| COMMD4 | 2.2 | 18.7 | 54.2 | 0.0 | 0.0 | 0.0 | Membrane protein trafficking | 26965651;25355947 |
| COMMD2 | 2.1 | 23.2 | 41.4 | 0.0 | 0.0 | 0.0 | Membrane protein trafficking | 26965651;25355947 |
| COMMD8 | 10.3 | 12.4 | 18.1 | 0.0 | 0.0 | 0.0 | Membrane protein trafficking | 26965651;25355947 |
| COMMD1 | 4.6 | 9.9 | 23.7 | 0.0 | 0.0 | 0.0 | Membrane protein trafficking | 26965651;25355947 |
| FLOT1 | 3.1 | 4.8 | 17.0 | 0.0 | 0.0 | 0.0 | Lipid raft protein promte<br>receptor surface presentation | 21399631;22732732 |
| PPFIA1 | 1.9 | 8.3 | 8.9 | 0.0 | 0.0 | 0.0 | Receptor surface presentation | 15750591 |
| HDDC2 | 3.0 | 5.5 | 8.4 | 0.0 | 0.0 | 0.0 | N.A. | N.A. |
| WNK1 | 1.8 | 2.1 | 7.6 | 0.0 | 0.0 | 0.0 | Regulate NCC membrane<br>abundance | 12671053;11498583 |
| IPO4 | 4.6 | 13.0 | 27.7 | 0.0 | 2.2 | 2.3 | Histone binding and histone<br>nuclear import | 30177573 |
| CD2AP | 1.8 | 12.5 | 13.4 | 0.0 | 2.2 | 0.0 | Facilitate the trafficking of<br>internalized growth factor<br>receptors. Risk factor of<br>Alzheimer Disease. | 12672817;21460841 |

Note: Flag-IP/MS was performed in Hela cells stably expressing Flag-TRIM72. Control, naïve Hela cells.

In 293FT cells, we noticed that a large molecular weight TRIM72 (∼110kD) was detected by Flag-antibody (Figure 5A), which was insensitive to DDT, reminiscent of the large molecular weight TRIM72 (∼110kD) shown in the FUS-R521C mutant neural tissues, although it was much weaker than that shown in the mutant neural tissues (Figure 1 and Figure S3B. To examine which part(s) of TRIM72 is required for the interaction between TRIM72 and CCDC22, we expressed Flag-tagged ΔPRY/SPAY and PRY/SPAY in 293FT cells together with CCDC22-Myc (Figure 5B and 5C). Our Flag-IP demonstrated that ΔPRY/SPAY but not PRY/SPAY is sufficient for the interaction with CCDC22. In addition, ΔPRY/SPAY but not PRY/SPAY displayed a small and a large molecular weight band, latter of which we speculated could be a dimer and was insensitive to DTT (10 mM), urea (8 M), and β-ME (200 mM) (Figure S17A, reminiscent of the large molecular weight TRIM72 (∼110kD) shown in the FUS-R521C mutant neural tissues (Figure S3B. To test it, we immunoprecipitated ΔPRY/SPAY with anti-Flag M2-beads (Figure S17B. Compared to naive control, two additional bands stained with Coomassie blue appeared only in the Flag-IP products with corresponding molecular weight. Our Flag-IP/MS of these two bands suggest that the large band is a dimmer of ΔPRY/SPAY monomer, because except for TRIM72 no other protein was identified in the large molecular weight bands by our IP/MS (Figure S17B. In addition, our data suggested that both monomer and dimer of TRIM72 are able to interact with CCDC22 and the presence of PRY/SPAY domain inhibits the dimer formation of TRIM72.

### Expression of TRIM72 does not bother the interaction between CCDC22 and CCDC93

To further identify which domain of TRIM72 is required for the interaction with CCDC22, we generated various of truncated TRIM72 (Figure 5B and 5D). Although each truncated TRIM72 had reasonable expression in 293FT cells, only the ΔCoiled-coil was not able to interact with CCDC22, suggesting that TRIM72 interacts with CCDC22 through its Coiled-coil domain. To examine whether the interaction is involved in the protective effect of TRIM72 against oxidative stress, we stably expressed full-length (FL), ΔRing, ΔB-box, ΔCoiled-coil, and ΔPRY/SPAY in 293FT cells and measured cell dehydrogenase activity by CCK-8 assay (Figure S18). In comparison with empty vector control, expression of FL significantly protected the cells from AS-induced oxidation, as well as expressions of ΔRing and ΔB-box, indicating that Ring and B-box domains are not necessary for the protection (Figure S18). However, expressions of ΔCoiled-coil and ΔPRY/SPAY showed no effect on the protection, indicating that both Coiled-coil and PRY/SPAY domains are required for the protection effect of TRIM72 full-length.

We next asked whether expression of TRIM72 affects interaction between CCDC22 and CCDC93. To this end, CCDC22-Myc was co-expressed with EGFP-TRIM72 or CCDC93-Flag in 293FT cells and the interactions of CCDC22/TRIM72 and of CCDC22/CCDC93 were confirmed by Myc-IP and Flag-IP, respectively (Figure 5E). We then expressed CCDC22-Myc, CCDC93-Flag, and EGFP-TRIM72 together in 293FT cells and the interaction of CCDC22/CCDC93 was not impaired or reduced by expression of TRIM72, evidenced by both Myc-IP and Flag-IP, suggesting that the interaction of CCDC22/CCDC93 is not bothered by TRIM72 expression.

As a top hit of TRIM72 interacting partners, CCDC22 contains an N-terminal calponin homology-like (NN-CH) domain and C-terminal coiled-coil (CC) domain (Figure S19A. To examine how CCDC22 interacts with CCDC93, we expressed CCDC22-Myc variants together with Flag-tagged CCDC93 in 293FT cells and our Flag-IP results suggested that coiled-coil domain (321-667a.a.) of CCDC22 is sufficient for the interaction and 447-667a.a. of the coiled-coil does not mediate the interaction between CCDC22 and CCDC93 (Figure S19B. However, the 447-667a.a. of CCDC22 is crucial for interaction with TRIM72 (Figure S19C, which is consistent with our findings that expression of TRIM72 does not bother the interaction between CCDC22 and CCDC93.

### Structural prediction of the TRIM72-Commander complex

To illustrate the detailed interfaces between TRIM72 and Commander complex, we employed AlphaFold3 to form a potential protein complex of TRIM72 (dimer), CCDC22 (390-627a.a.), CCDC93 (380-631a.a.), VPS29, VPS26C, and VPS35L. We then adapted the resulting complex to the previously completed Commander complex by the overlap between the two (Healy et al., 2023) (Figure 5F). Consistent with our results shown in Figure 5D and S19C, AlphaFold3 predicted an interaction between TRIM72 (131-166a.a., 241-270a.a.) and CCDC22 (480-627a.a.), primarily through their coiled-coil domains (Figure 5F). The part of CCDC22 (480-627a.a.) forms an alpha-helix structure that interacts with dimerized TRIM72, which does not bother the interaction between CCDC22 and CCDC93 via their Coiled-Coil domain. These data suggest that the structural predictions by AlphaFold3 fit well with our experimental results.

To gain better understanding of interactions between TRIM72 and CCC complex, a possible interaction site of CCDC22 (K506Q507) was predicted with a close proximity of TRIM72 in our model (Figure 5F). These two residues (K506Q507) are evolutionarily conserved across the species (Figure 5G), suggesting a functional significance of these two residues. In addition, we ended up with another two pairs of residues (L532R534 and D540E541) in the same CCDC22 Coiled-Coil domain, which are evolutionarily conserved (Figure 5G, 5H, and 5I). To demonstrate that these residues play a critical role in mediating the interaction between TRIM72 and CCDC22, we generated three mutant CCDC22 with Myc tags, including K506D/Q507L-Myc, L532Q/R534E-Myc, and D540K/E541R-Myc, and expressed these mutant CCDC22 together with Flag-tagged TRIM72 in cells (Figure 5J). Although the expressions of mutant CCDC22 had a reasonable expression level in the cells, the interaction between TRIM72 and CCDC22 was largely decreased by expressions of K506D/Q507L-Myc and L532Q/R534E-Myc, but not D540K/E541R-Myc (Figure 5J). Indeed, when we examined the site of these three pairs of residues, we noticed that Q507 and L532 are the first (a) or fourth (d) positions of the predicted heptad repeats (a seven-residue periodicity) of CCDC22 Coiled-Coil domain (Figure 5G and 5I). In contrast, D540E541 residues are the second (b) and third (c) positions of a potential heptad repeat, position of which are less critical than ‘a’ and ‘d’ site in mediating the protein/protein interaction (Burkhard et al., 2001). Taken together, our experimental data supports the structural predictions of TRIM72 and CCC complex by AlphaFold3.

TRIM72 interacts with both CCDC22 and CCDC93 to form a large composite complex with the Retriever complex that is anchored to early endosomal membrane through interaction between VPS26C and a sorting nexin SNX17 with a phosphoinositide-binding module (McNally et al., 2017). Concurrently, the PRY/SPRY domain of TRIM72 is positioned on the same side of the membrane as VPS26C in the composite complex, suggesting that PRY/SPRY domain of TRIM72 facilitates membrane anchoring (Figure 5F). Indeed, the PRY/SPRY domain of TRIM72 was reported to recognize negatively charged phospholipid membrane via a clustering of positive charged residues on the surface of PRY/SPRY (Ma et al., 2023; Park et al., 2023). Altogether, these results further suggest the crucial role of TRIM72 in large composite complex formation that may mediate membrane-associated biological functions.

### CCC and Retriever complexes participate TRIM72-mediated antioxidation and membrane repair

We next asked whether CCC and Retriever are involved in the TRIM72-mediated antioxidation. To this end, we employed previously reported dual gRNA approach to generate *CCDC22* KO cells (Figure S20) (Piao et al., 2022). Expression of TRIM72 protected control but not *CCDC22* KO cells from AS-induced oxidative stress (Figure S21). In addition, the expression was not able to protect *SNX17* or *VPS35L* KO cells from the oxidative stress; however, *CCDC22*, *SNX17*, or *VPS35L* KO alone did not severe the AS-induced oxidative stress in these KO cells without TRIM72 expression (Figure S20 and S21). These data suggest that both CCC and Retriever complexes are required for the TRIM72-mediated antioxidation.

In order to examine whether CCC and Retriever complexes are also required for previously reported TRIM72-mediated membrane repair (Cai et al., 2009), we employed laser light to induce membrane damage in U2OS cells (Figure 5K), the severity of which can be quantified by PI fluorescent intensity at the damage site as previously reported (Huang et al., 2022). In order to avoid the variation from individual cell to cell during photodamage, we selected single cell clones from *CCDC22* or *SNX17* KO cells and expressed EGFP-tagged TRIM72 (EGFP-TRIM72) or EGFP to localize the individual cell (Figure 5K and S22). Indeed, compared to EGFP control cells, EGFP-TRIM72 expression significantly protected the cells from laser light-induced membrane damage (Figure 5K and 5L). However, *CCDC22* or *SNX17* KO significantly decreased membrane repair effect of TRIM72 but the gene KO did not further severe membrane damages induced by laser light in these KO cells without TRIM72 expression (Figure 5L).

To further investigate the role of TRIM72 in mediating cell membrane repair, we utilized Correlative Light and Electron Microscopy (CLEM) to examine the damage site 10 minutes after photodamage. Membrane damages were evidenced in the U2OS cells no matter expressing EGFP or EGFP-TRIM72 (Figure 5M). Compared to the cells expressing EGFP, we observed that vesicles with varying sizes (> 200 nm, < 1 µm) were recruited to damage site 10 minutes after photodamage in U2OS cells expressing EGFP-TRIM72, suggesting that expression of TRIM72 promotes vesicle recruitment to damage site for membrane repair after membrane damage. We noticed that most of the recruited vesicles at damage site were empty with low electron density inside, suggesting that these vesicles do not belong to multivesicular bodies (MVB).

We next asked whether neuronal TRIM72 expression is involved in membrane repair. To this end, we cultured C/C and C/C;−/− cortical neurons and damaged soma with laser light (Figure 5N and Movie S2). C/C;−/− neurons displayed severe neurite injury/breaking as early as 271.0 ± 133.3s after soma photodamage, which was much quicker than that shown in C/C cultured neurons (551.0 ±50.43s) (Figure 5N and 5O). However, the average of neurite breaking time was restored by neuronal expression of EGFP-tagged full-length TRIM72 (FL) by lentivirus in these neurons (523.8 ±132.3s) (Figure 5O). Expression of ΔCoiled-coil of TRIM72 did not restored the neurite breaking time, suggesting that interaction between CCDC22 and TRIM72 is required for the neuronal protection effect of TRIM72. We noticed that expression of PRY/SPAY alone failed to restore the neurite damages in these C/C;−/− cultured neurons (Figure 5O).

In order to examine whether CCDC22 is required for TRIM72-mediated neurite protection, we employed lentivirus-based shRNA to knockdown *Ccdc22* in the cultured neurons (Figure S23A and S23B). Among three *Ccdc22* shRNAs, one shRNA (shRNA2) showed the strongest knockdown efficiency. Indeed, knockdown of *Ccdc22* by shRNA2 significantly shortened the neurite breaking time in C/C cultured neurons (Figure S23C and Movie S3), in which the TRIM72 expression is high (Figure 3E). Taken together, our results suggest that CCC and Retriever complexes are involved in TRIM72-mediated antioxidation, membrane repair, and neurite protection, likely through interaction between TRIM72 and Commander complex.

### TRIM72 presents in exosome

In CSF of FUS-R521C knockin rodent models, we observed significantly higher level (715.07 ± 73.07pg/μL CSF for mouse) of TRIM72 in comparison with that of wildtype controls, suggesting that TRIM72 is secreted by neurons highly expressing TRIM72 (Figure 6A). Indeed, TRIM72 was released to the culture medium in 293FT cells stably expressing Flag-tagged FL TRIM72 (Figure 6B and 6C). Deletion of Ring or B-box domain (ΔRing and ΔB-box) did not significantly affected TRIM72 release; however, ΔCoiled-coil and ΔPRY/SPAY abolished the release, suggesting that Coiled-coil and PRY/SPAY domains of TRIM72 are required for the process (Figure 6B and 6C).

**Figure 6.**
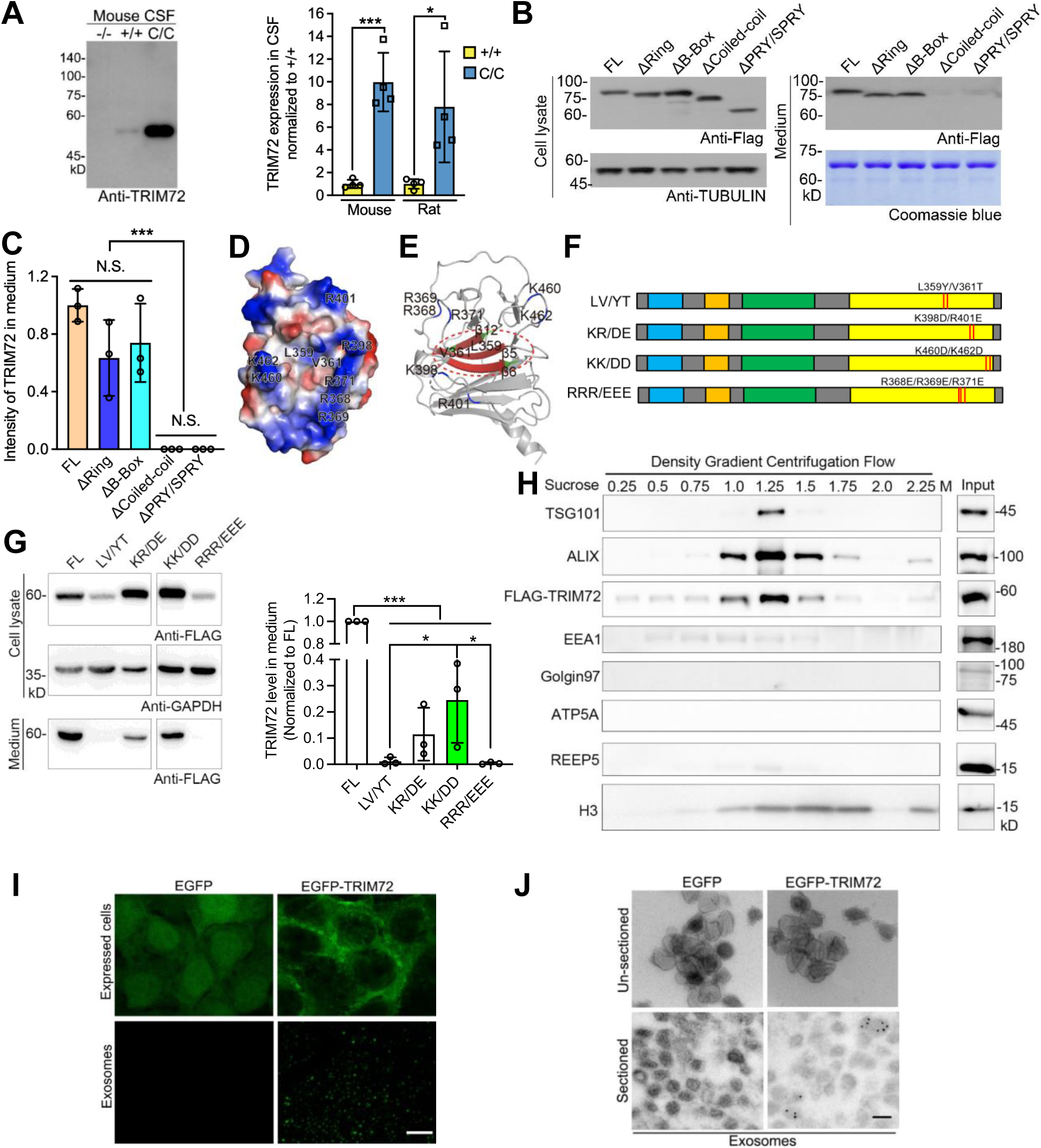
TRIM72 presents in exosome. (**A**) TRIM72 levels in the CSF in mice with the indicated genotypes. The same amount of CSF (5 μl) was loaded. Data was summarized in right. (**B** and **C**) 293FT cells were stably expressed TRIM72 FL, ΔRing, ΔB-box, ΔCoiled-coil, and ΔPRY/SPRY, respectively. Cell lysate and culture medium were applied for western blot to examine the release (**B**). Total protein amount was visualized with Coomassie blue. Data summary was shown in (**C**). (**D** and **E**) The key positive charged residues form a horse-shoe shape in PRY/SPRY domain and two key residues (L359 and V361) in a beta-sheet of TRIM72. (F) Potential key residue mutations of TRIM72 PRY/SPRY. (G) TRIM72 FL and PRY/SPRY mutations were stably expressed in 293FT cells, respectively. Cell lysate and culture medium were applied for western blot to examine the release (left). Data summary was shown in (right). (H) Culture media of 293FT cells expressing FLAG-TRIM72 was subjected to sucrose density gradient centrifugation. ALIX and TSG101, exosome markers; EEA1, Golgin92, ATP5A, and REEP5, other organelle markers. (**I** and **J**) 293FT cells were expressed EGFP and EGFP-TRIM72, respectively. The precipitation of exosomal flow from the cells was imagined (**I**). The precipitation was applied for GFP immunogold staining before and after sectioning (**J**). Scale bar, upper, 10 µm; lower, 100 nm. In **A**, **C**, and **G**, values were presented as mean ± SD. \**p* <0.05, \*\**p* <0.01, \*\*\**p* <0.001, T-Test or ANOVA SPSS.

Previously, TRIM72 recognizes negatively charged phospholipid membrane through its PRY/SPAY domain, especially, several positively charged residues that form a horseshoe shape in the surface of PRY/SPRY, including K460K462 and R368R369R371 (Park et al., 2023) (Figure 6D and 6E). To examine whether these residues contribute to the secretion, we generated charge-reversal mutants of these residues, including K460D/K462D (KK/DD) and R368E/R369E/R371E (RRR/EEE) (Figure 6F). The mutants KK/DD and RRR/EEE significantly impaired the TRIM72 release, especially RRR/EEE almost abolished the release (Figure 6G). From the TRIM72 structure (Figure 6D and 6E) (Park et al., 2023), the other two positively charged residues (K398R401) drew our attention, which have a close proximity to R368R369R371. Indeed, the charge-reversal mutants of these two residues (K398D/R401E) significantly impaired the release (Figure 6G). These results suggested that TRIM72 secretion depends on the interaction between the positive charges on the PRY/SPRY and the lipid membrane.

In addition to the positively charged residues in the horseshoe shape, we also examined the two conserved residues (L359 and V361) in a β-sheet (β5) of PRY/SPRY on TRIM72 release (Figure 6E, 6F and Figure S24). The β5, β6 and β12 topologically are close to each other and are major components to form a prominent pocket, which are predominantly composed of nonpolar amino acids (Figure 6E) (Park et al., 2010). To study whether the nonpolar property is essential for the release, we generated L359Y/V361T mutant that change nonpolar to polar amino acids without affecting hydrophobicity of these two residues. L359Y/V361T almost abolished TRIM72 secretion into the culture media (Figure 6G), suggesting that the nonpolar property of the pocket is crucial for the secretion.

In order to examine whether CCC and Retriever complexes are required for the release, we expressed Flag-TRIM72 in *CCDC22*, *SNX17*, and *VPS35L* KO cells. The release was significantly reduced in these KO cells with or without treatment of AS (Figure S25), although oxidative stress in general increased the release as previously reported (Cao et al., 2010). These data suggested that TRIM72 release depends on the CCC and Retriever complexes.

To further examine how TRIM72 is released, we applied the culture medium from 293FT cells stably expressing TRIM72 for sucrose density gradient ultracentrifugation, a powerful technique for fractionating cellular organelles and macromolecules (Momen-Heravi, 2017) (Figure S26). TRIM72 was co-fractionated with exosome, represented by exosome markers ALIX and TSG101, and exosome associated protein histone H3 (Tutanov et al., 2022), as well as early endosome, evidenced by EEA1, but not other cellular organelles, including Golgi apparatus (Golgin92), mitochondrion (ATP5A), and ER (REEP5) (Figure 6H). We then expressed EGFP and EGFP-TRIM72 in 293FT cells and purified the exosomes (Figure 6I). The exosomes from EGFP-TRIM72-expressed but not EGFP-expressed cells displayed EGFP fluorescence, suggesting that TRIM72 presents in exosome. Our EGFP immunogold electron microscopy (EM) data confirmed that TRIM72 presents in the exosome, more specifically inside of exosome, evidenced by EGFP-positive immunogold signals found in sectioned but not un-sectioned exosome (Figure 6J). As a negative control, the EGFP-positive immunogold signals were found in neither sectioned nor non-sectioned exosomes derived from EGFP-expressed cells.

### Exosomal TRIM72 is protective

Compared to FL, ΔPRY/SPAY and PRY/SPAY were present much less in exosomal fractions by our sucrose density gradient ultracentrifugation (Figure 7A), suggesting that PRY/SPAY is required but not sufficient for TRIM72 presence in exosome. In addition, expressions of FL and PRY/SPAY but not ΔPRY/SPAY significantly improved presence of exosomal proteins, including ALIX and TSG101, in exosomal fractions (Figure 7A), suggesting that PRY/SPAY domain of TRIM72 is essential for the improvement of TRIM72-mediated exosome formation.

**Figure 7.**
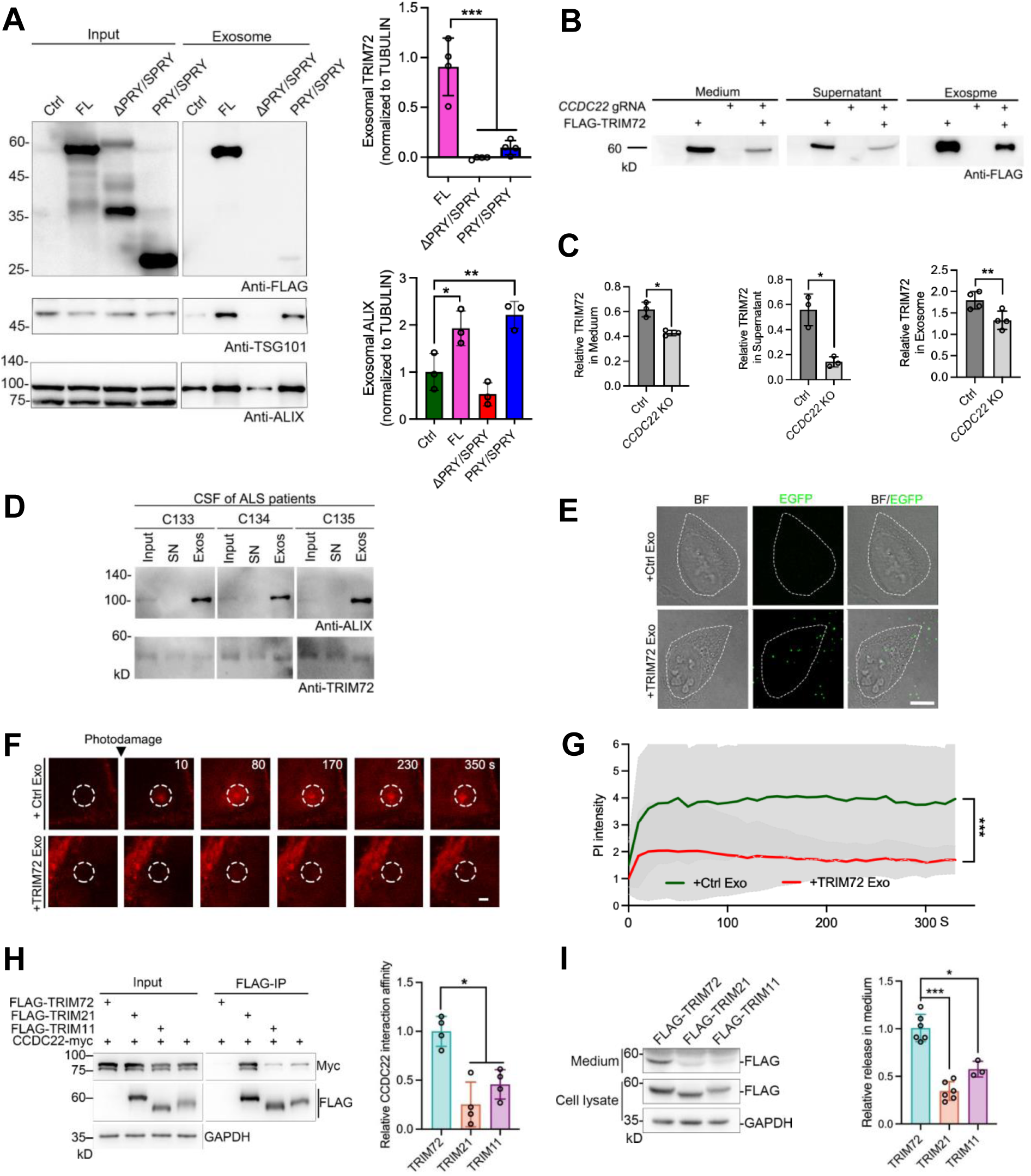
Exosomal TRIM72 is protective and CCDC22 promotes release of TRIMs. (**A**) 293FT cells were stably expressed FLAG-tagged TRIM72 (FL), ΔPRY/SPRY and PRY/SPRY, respectively. The culture media was subjected to sucrose density gradient centrifugation. Normalized expressions of exosomal TRIM72 (upper right panel) and ALIX (lower right panel) were summarized. (**B** and **C**) Culture media from naïve or *CCDC22* KO cells with or without FLAG-TRIM72 expression were subjected to sucrose density gradient centrifugation (**B**). The amounts of FLAG-TRIM72 in the medium, supernatants and exosomes were summarized (**C**). (D) TRIM72 levels in the CSF (input) and corresponding supernatants (SN) and exosomes (Exos) from ALS patients. (E) Exosomes isolated from the culture medium of 293FT cells expressing EGFP (Ctrl Exo) or EGFP-TRIM72 (TRIM72 Exo) and extracellularly applied to U2OS cells (also see **Figure S28** and the **Movie S4**). Cell contours were highlighted with dotted lines. (**F** and **G**) U2OS cells were photodamaged and monitored by time-lapse imaging after extracellular applications of indicated exosomes (**F**). Circles indicate the damage sites. Normalized PI intensities were summarized (**G**). (**H** and **I**) TRIM21 or TRIM11 interacted with CCDC22 (**H**) and was released in the culture medium (**I**). In **A**, **C**, **G**, **H** and **I**, values were presented as mean ± SD. \**p* <0.05, \*\**p* <0.01, \*\*\**p* <0.001, T-Test or ANOVA SPSS. In **E**, scale bar, 10 µm; in **F**, 2 µm.

To examine whether CCDC22 is present in exosome, we expressed CCDC22-Myc in 293FT cells and detected CCDC22-Myc in the exosomal fraction in the cells stably expressing or not expressing FLAG-TRIM72, indicating that association of CCDC22 with exosome does not depend on the expression of TRIM72 (Figure S27A and S27B). To examine the roles of CCDC22 in exosomal presence of TRIM72, we purified the exosomes from control or *CCDC22* KO cells expressing FLAG-TRIM72 (Figure 7B and 7C). Consistent with our data that *CCDC22* KO significantly impairs TRIM72 release (Figure S25), *CCDC22* KO also significantly reduced the TRIM72 levels in exosomal fraction and supernatant (Figure 7B and 7C). Our data suggest that TRIM72 presents in exosome and promotes exosome formation, which most likely depends on interaction between TRIM72 and CCDC22. Indeed, the exosomal interaction between TRIM72 and CCDC22 was documented in exosomal fractions of cells expressing both CCDC22-Myc and FLAG-TRIM72 (Figure S27C.

To examine whether CSF TRIM72 exists in exosome, we performed ultracentrifugation by using the patient CSF (Figure 7D). As exosome preparation evidenced by presence of ALIX, TRIM72 appeared in the exosome fractions as well as inputs and supernatants. The relative supernatant TRIM72 level was 46.7% ±0.6 (mean ±SD) to that of input, suggesting that fairly amount of TRIM72 in human CSF are in the exosomes. To examine whether application of exosomes containing TRIM72 is functional, we purified the exosomes from 293FT cells expressing EGFP (control exosome) or EGFP-TRIM72 (TRIM72 exosome) (Figure S28A. Then we extracellularly applied control and TRIM72 exosomes in U2OS cells and observed intracellular EGFP fluorescence in the cells that we applied for TRIM72 but not control exosomes (Figure 7E and Movie S4). Extracellular application of TRIM72 but not control exosomes protected U2OS cells from membrane damages caused by laser lights (Figure 7F and 7G). In addition, we observed intracellular EGFP fluorescent signals when we extracellularly applied for TRIM72 but not control exosomes in cultured neurons and the application protected the neurons from neurite damages induced by laser lights (Figure S28B. Therefore, we concluded that extracellular application of exosome containing TRIM72 protect cell from membrane damage.

TRIM11, TRIM21 and TRIM72 belong to TRIM C-IV subfamily, members of which contain RING-finger, B-box, Coiled-Coil, PRY/SPRY domains (Hatakeyama, 2017; Ozato et al., 2008). To examine whether TRIM11 and TRIM21 are functional similar to TRIM72, we expressed FLAG-tagged TRIM11, TRIM21, or TRIM72, together with CCDC22-Myc in the 293FT cells (Figure 7H). Interaction between other TRIMs and CCDC22 was documented in these cells, but the interaction was much weaker than that of TRIM72. Correspondingly, releases of TRIM11 and TRIM21 were much less than that of TRIM72 in these cells (Figure 7I). Therefore, our findings raise a possibility that TRIM C-IV subfamily members may protect cell through a similar mechanism as TRIM72 we disclosed.

### CSF TRIM72 level associates ALS disease progression and AAV-mediated TRIM72 expression in an ALS patient

To measure TRIM72 levels in the CSF of ALS patients, we collected CSF from 168 sporadic ALS patients (100 men and 68 women) and 34 age-matched non-ALS controls with no history of neurological diseases from local hospitals (Table 2). Compared to non-ALS controls, the level of NfL (neurofilament light chain), a biomarker of axonal damage, in patient CSF was significantly higher (Table 2). Subsequently, we measured TRIM72 levels by western blot and detected a band with a similar molecular weight as TRIM72 from rodent CSF (Figure S29A. To confirm the protein identity of the band, we conducted mass spectrometry (MS) and the results showed a strong correlation (r = 0.989, n = 8) with that of our western blot results (Figure S29B.

**Table 2.** Baseline of clinical features in the cohort.

| | Numbers | Male:<br>Female | Age<br>($p = 0.29$) | TRIM72<br>(pg/μl)<br>($p = 0.017$) | NfL in CSF<br>($p < 0.05$) |
| --- | --- | --- | --- | --- | --- |
| ALS | 168 | 100:68 | 53.88±11.07 | 34.4±25.2 | 4483.81±3315.92 |
| Non-ALS | 34 | 16:18 | 56.06±9.88 | 23.6±13.2 | 1049.67±1733.78 |

CSF TRIM72 levels in ALS patients (34.4 ± 25.2 pg/µl) were significantly higher than that of non-ALS controls (23.6 ±13.2 pg/µl) (Figure 8A and Table 2). The TRIM72 level was negatively correlated with ALS disease stages as illustrated by the King’s clinical staging system (KCSS) (Figure 8B). Kaplan‒Meier survival analysis ascertained higher CSF TRIM72 levels associated with better disease prognoses, starting from the time of diagnosis (Figure 8C). In detail, patients with CSF TRIM72 level over 44.52 pg/µl (> 75% percentile) had significantly better prognosis (mean estimated survival, 110.62 months) than that (73.96 months) of patients with low TRIM72 level (< 44.52 pg/µl). The diagnostic performance of CSF TRIM72 was further assessed by ROC (receiver operating characteristic) curve analysis. With an optical cut-off value (24.64 pg/μl), the Area Under the ROC curve (AUC) was 62.11% (*p* = 0.026) (Figure 8D), which is lower than the AUC score (> 0.8) acceptable for diagnostic performance (Muller et al., 2005; Nahm, 2022). We also checked the correlation between CSF TRIM72 level and various factors, including sex, disease onset position, diagnostic delay from symptoms, ALSFRSR, and patient ages, but did not observe any statistical significance (Figure S30 and Table 3). In order to examine the relationship between target tissue expression of TRIM72 and motor function, we examined spinal cord expression of TRIM72 in 8-month C/C male mice (Figure S31). The expression was correlated well with travel distance detected by Open Field test (R^2^ = 0.7074, *p* < 0.0012), suggesting that higher target tissue expression of TRIM72 is associated with better motor performance.

**Figure 8.**
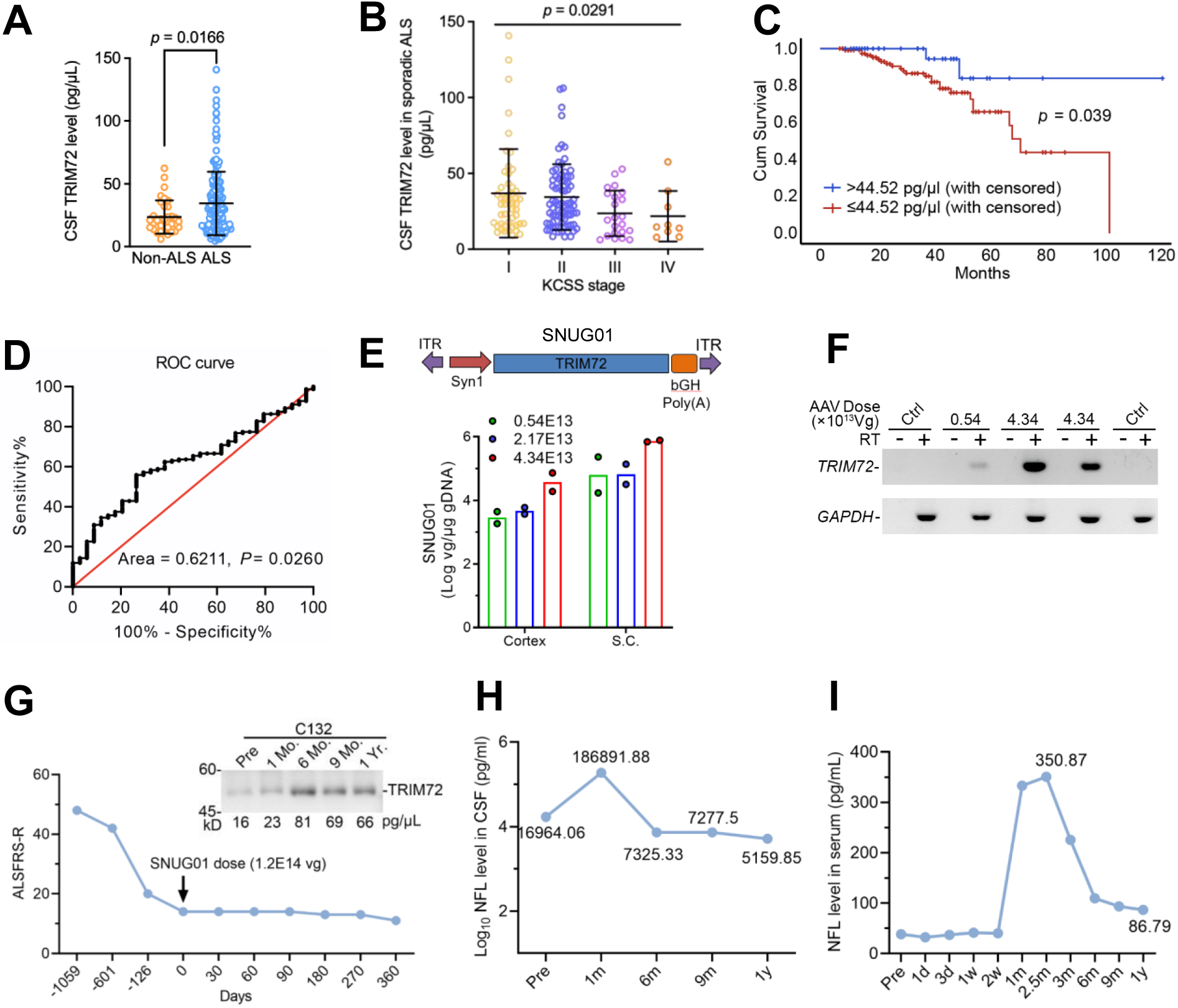
CSF TRIM72 level was negatively correlated with ALS disease progression. (A) Significant increase of CSF TRIM72 level (34.4±25.2 pg/μl) in ALS patients, compared to that (23.6±13.2 pg/μl) of Non-ALS group (also see **Table 2**). (B) With the disease progression evidenced by KCSS, TRIM72 level was significantly down-regulated in patients (*p* = 0.0291). KCSS, King’s clinical staging system. (C) KM (Kaplan Meier) survival analysis showed that the survival time of Q1 patients with TRIM72 in CSF was significantly longer than other patients in the population (*p* = 0.039, 110.61 vs 73.96 month). (D) ROC (Receiver Operating Characteristic) curve analysis was used to evaluate the accuracy of employing CSF TRIM72 level as a diagnostic tool (62.11%, *p* = 0.0260). (E) AAV9-based expression cassette for *TRIM72* (SNUG01, upper). Syn1, a neuronal promoter; bGH Poly(A), bovine growth hormone Poly(A) sequence. Biodistributions of SNUG01 DNA in the cortex and spinal cord (S.C.) of macaques treated with indicated SNUG01 doses (lower). Samples were collected 3 months after treatment. (F) Spinal cord *TRIM72* mRNA expression in macaque intrathecally injected with indicated SNUG01 dose. Ctrl, spinal cords of macaques without SNUG01 injection. RT, reverse transcriptase. *GAPDH* served as loading control. (G) ALS Functional Rating Scale in its revised version (ALSFRS-R) of a patient before and after SNUG01 treatment. SNUG01 (1.2E14vg) was administered by intrathecal injection. Expressions of TRIM72 in the patient CSF before (Pre), 1-month (1 Mo.), 6-month (6 Mo.), 9-month (9 Mo.), and 1-year (1 Yr.) after treatment were semi-quantified by western blot. (**H** and **I**) NFL levels in the CSF (**H**) and blood (**I**) of the patient before and after SNUG01 treatment. In **A**, and **B**, values were presented as mean ± SD. In **L**, \*\**p* <0.01, T-Test SPSS. In **H**, and **I**, d, day; w, week; m, month; y, year.

**Table 3.** Baseline for clinical features of ALS patients.

|  | Onset age | Onset position<br>(Bulbar: Limbs) | ALSFRS-R | ECAS |
| --- | --- | --- | --- | --- |
| ALS | 51.77±11.36 | 25:143 | 41.41±4.80 | 87.70±26.38 |

To explore the therapeutic potential of TRIM72 in patient, we employed Syn1 promoter and AAV9, latter of which has been used for SMA (spinal muscular atrophy) gene therapy (Mendell et al., 2017), to express our GOI (gene of interest) *TRIM72*. We named the resulting AAV9 particle SNUG01 (Figure 8E). Before applying to patient, we intrathecally (IT) injected SNUG01 in cynomolgus macaques with 3 dosages (0.54, 2.17, and 4.34 E13 vg/animal, 2 animals a dose), which are relevant to 1.2, 4.8, and 9.6 E14/person in human, according to body weight difference between human and cynomolgus monkey, in order to examine the safety of SNUG01 (Figure 8E). The peaks of virus shedding in blood, feces, and urine were detected 2 days after injection, which went back to normal one month after injection in feces and urine but maintained a relatively high level in peripheral blood (Figure S32A-C). The anti-drug antibody (ADA) in peripheral blood reached peak a week after injection (Figure S32D. SNUG01 DNA levels were detected 3 months after injection in the spinal cord and cortex in a dose-dependent manner (Figure 8E), suggesting target engagement. GOI mRNA was detected in plus but not minus RT groups 3 months after injection in the spinal cords (Figure 8F). Compared to level of CSF TRIM72 detected by western blot before SNUG01 injection, the level was increased or kept no change 3 month after the injection in these macaques (Figure S32E.

A panel of physiological and biochemical tests, including body weight, body temperature, blood pressure, and electrocardiogram (ECG), were examined and monitored during our study, and no obvious abnormality was detected. The values of alanine aminotransferase (ALT) and aspartate transaminase (AST) were transiently increased 2 days after SNUG01 injection, which went back to normal in a week (Figure S32F and S32G). Taken together, our data suggested that our administration of SNUG01 is safe in macaque.

We then applied SNUG01 intrathecally to an ALS patient with FUS-R521C mutation (KCSS stage IV) (Registration number: ChiCTR2400085764). The patient received 1.2E14 vg SNUG01 and the virus shedding in urine, feces, saliva, and blood were transiently increased in days and went back to baselines in weeks or months (Figure S33A-S33D). In contrast, AAV9-neutralizing antibody in peripheral blood or CSF reached plateau in 3 or 6 months after treatment and kept in a relatively high level at the last checking point we examined (Figure S33E and S33F), suggesting target engagement. Till we submitted this manuscript, the patient has been survived for 1.5 year without serious adverse events (SAEs).

CSF TRIM72 expressions in this patient were monitored before and after SNUG01 administration (Figure 8G). Compared to that of pre-treatment (16 pg/μl), the level was slightly increased 1-month after treatment (23 pg/μl), which was increased about 4 times (81 pg/μl) 6 months after the treatment and kept in a relative high level 9- and 12-month after treatment, suggesting target engagement. Compared to the sharp drop of ALSFRS-R score before treatment, the drop became flat after the treatment (Δ2/ year) (Figure 8G). The disease progression of the patient was slowed down and the overall situation was stable after treatment. A previous study reported a transient upregulations of NfL in macaque CSF and serum a month after AAV9 treatment (Johnson et al., 2023). Consistently, the levels in CSF and serum were transiently upregulated in this patient 1 month after SNUG01 treatment (Figure 8H and 8I). However, compared to the level before our treatment (16964.1 pg/ml), the level in CSF were dropped to 7325.3 pg/ml 6 months after SNUG01 treatment, suggesting potential drug efficacy of SNUG01. The level kept declining 9 month (7277.5 pg/ml) and 1 year (5159.9 pg/ml) after treatment. The NfL in peripheral blood showed a similar pattern as we examined in CSF.

## Discussion

Here, we reported that TRIM72 is highly expressed in neuronal but not non-neuronal tissues in the both FUS-R521C knock-in mouse and rat ALS models (Figure 1). Loss-of-function of *Trim72* worsens ALS progression in the knockin mouse and induces mislocalization of mutant FUS and oxidative stress in the mutant spinal cord and brain (Figure 2 and 3). Neuronal expression of TRIM72 by AAV extends lifespan in both SOD1-G93A and TDP-43-A315T ALS models (Figure 4). TRIM72 joins in Commander complex, which we would like to name it Commander-TRIM, and facilitates membrane repair probably through direct interaction with CCDC22, a protein component of CCC complex (Figure 5). The interaction between TRIM72 and CCDC22 depends on a Coiled-Coil-dependent oligomerization (Figure 5F-5J and Movie 5). Exosomal expression of TRIM72 depends on TRIM72/CCDC22 interaction and TRIM72 PRY/SPRY domain, which tethers membrane by a cluster of positively charged residues in a horseshoe shape and two nonpolar residues in a groove (Figure 6). Exosomal TRIM72 protects cell from membrane damage and is detected in ALS patient CSF (Figure 7). CSF TRIM72 level associates with ALS disease progression and neuronal expression of TRIM72 by AAV9 slows down disease progression in an ALS patient (Figure 8).

Like other neurodegenerative diseases, ALS display high heterogeneity in pathologies and clinical manifestations. Several common pathological hallmarks have been identified, including abnormal RBP aggregates and protein misfolding. Notably, vesicle trafficking dysregulation, especially dysfunction in the endosomal/lysosomal pathway or axonal transport/exosome pathway, recently has emerged as a critical mechanism in ALS pathogenesis (Todd et al., 2023). However, the exact mechanisms according to how endomembrane system contributing to the disease remain to be elucidated. Many molecular/cellular mechanisms have been proposed to be involved in membrane repair, including but not limited to SNAREs (Soluble N-ethylmaleimide-sensitive factor attachment protein receptors)/exocytosis, ESCRT (endosomal sorting complexes required for transport)/endocytosis, ESCRT/scission (Ammendolia et al., 2021). TRIM72 has been demonstrated to protect cell from membrane damage and associated with vesicles; however, the detailed mechanisms of TRIM72-mediated membrane repair are still largely unknown. Here, we propose that TRIM72 facilitates vesicle trafficking path, more specifically a Commander/Retriever path, to participate in membrane repair and protect neuron from oxidative insults and neurodegeneration. Given that structural similarity between Retriever and Retromer complexes and involvement of dysfunctions of Retromer in other neurodegenerative diseases (Gershlick and Lucas, 2017; Hu et al., 2015; Schreij et al., 2016), the membrane damage prevented by TRIM family proteins could be a common protective mechanism for neurodegenerative diseases.

Retriever complex, a recently identified novel protein complex in mediating Retromer-independent endosomal trafficking of various proteins (McNally et al., 2017). Recently, a structural biology study has shown how Retriever interacts with the CCC complex to form the Commander complex (Healy et al., 2023). Our data suggests that TRIM72 and CCDC22 probably directly interact each other through their Coiled-Coil domains by a Coiled-Coil-dependent oligomerization, in which the key residues are located in the first or fourth positions of the heptad repeats (Figure 5I). High molecular weight of TRIM72 is seen in mutant FUS mouse CNS (Figure 1), which we speculate is a dimer of TRIM72 (Figure S17). TRIM72 dimer fits well into Commander complex by AlphaFold3 (Figure 5F), which provides two additional anchoring sites (two PRY/SPRY of TRIM72, Figure S34 and Movie 5) for membrane attachment/recognition, besides VPS26C/SNX17 anchoring site. The additional two anchoring sites provided by TRIM72 are supposed to stabilize the membrane attachment of the whole complex (Commander-TRIM), which may explain how TRIM72 facilitates membrane repair. Recently, E3 ligase activity-dependent neuronal protections of TRIM21 and TRIM11 have been disclosed and they belong to the same TRIM subfamily of TRIM72 (Hatakeyama, 2017; Ozato et al., 2008). Both TRIM21 and TRIM11 interact with CCDC22 and are released, although the interaction and release are not as strong as that of TRIM72 (Figure 7H and 7I). Therefore, an E3-independent neuronal protective mechanism could be common and shared by the TRIM family members.

There are two major insults from membrane damage: elevations of intracellular [Ca^2+^] ([Ca^2+^]_in_) and oxidation, due to high concentrations of extracellular [Ca^2+^] ([Ca^2+^]_ex_) and oxidative species (Ammendolia et al., 2021) (Figure S34D. However, both of them trigger TRIM72-mediated membrane repair, oligomerization, and release (Cai et al., 2009; Cao et al., 2010). Long-lasting high concentrations of intracellular Ca^2+^ and ROS are detrimental to neuron survival eventually. Loss-of-function of *Trim72* itself seems not sufficient to drive membrane damage in neurons without the presence of mutant FUS (Figure 3G and Figure S9). Expression of mutant FUS has been linked to oxidative stress and mitochondrion damages (Deng et al., 2015; Kodavati et al., 2024; Wang et al., 2018a). Indeed, in absence of TRIM72, we observed oxidative stress in the FUS mutant neurons (Figure 3G) and mislocalized FUS in the mutant spinal cord (Figure 3H and 3I), latter of which could be secondary to ROS (Zhang et al., 2020). As one of the major sources for membrane damage (Ammendolia et al., 2021), the intracellular ROS generated by mutant FUS damages motor neuron membrane; however, TRIM72 expression facilitates Commander/Retriever path to protect the mutant neuron from membrane damage and the damage-induced oxidative stress and neuron loss (Figure S34). We proposed in absence of TRIM72 the mutant FUS mislocalized to axon to form FUS-positive condensates that impair mitochondrion dynamics, a mechanism for axonal ROS generation mediated by mutant FUS (Figure S8 and Figure S34D. Expression of TRIM72 reduces axonal FUS-positive condensate formation and protects axon (Figure 5N and 5O), which may explain the NfL drops in the ALS patient after administration of AAV-TRIM72 (Figure 8H).

## Materials and methods

### Generation of *Trim72* knock-out mouse, animal care, and behavior tests

The CRISPR-Cas9 system was used to generate *Trim72* knock-out mouse lines. The gRNA (GTCAGGCACCTACACGGCCG) and *Cas9* mRNA were injected into C57BL/6J and FUS-R521C knock-in mouse embryos (Zhang et al., 2020). Successful *Trim72* knock-out was confirmed by PCR and sequencing. To minimize off-target effects, F0 heterozygous animals were backcrossed to C57BL/6J and FUS-R521C knock-in mice over 5 generations.

All mice were housed in isolated ventilated cages (maximum of six mice per cage) in the barrier facility at Tsinghua University. Mice were maintained on a 12-/12- hr light/dark cycle at 22-26°C with sterile pellet food and water ad labium. The laboratory animal facility has been accredited by AAALAC (Association for Assessment and Accreditation of Laboratory Animal Care International), and the IACUC (Institutional Animal Care and Use Committee) of Tsinghua University approved all animal protocols used in this study.

The open-field behavior test was performed as previously described (Zhang et al., 2020). Briefly, each mouse was placed in the center of an open field (50 ×50 cm) and tracked with the TopScan behavioral analysis system (CleverSys) for 10 min per trial, and total distance travelled was quantified. Rearing time was measured by the ratio of total standing time on hind paws to total test time (10 min) at each trial. For the rotarod behavior test, each mouse was placed on the spindle with an acceleration from 4 to 40 rpm. Three trials of 5 min each were performed per animal for 4 days. Each trial was followed by a 20-min break. The number of falls was recorded by the rotarod system (Med Associates Inc.). The average time on the spindle per trial was computed for statistical analysis.

Cynomolgus macaque’s husbandry and study were conducted in Shanghai Institute of Materia Medica, an Association for Assessment and Accreditation of Laboratory Animal Care International (AAALAC) accredited Good Laboratory Practice (GLP) facility. Study protocol were reviewed and approved by Test facility Animal Care and Use Committees with IACUC No. 2021-08-RJ-245.

### Electromyography (EMG) analysis

One-year-old mice of each genotype were subjected to EMG using the Keypoint Electromyography System (Medoc Ltd., Israel) (Han et al., 2006; Sun et al., 2016). Mice were anesthetized and the limbs were fixed for the recording sessions. Ground wire was placed on the abdomen and a sterilized concentric needle electrode was inserted into the quadriceps muscle. Abnormal spontaneous potentials were noted. Mice were stimulated at the groin to simulate a light contraction-equivalent motor unit potential (MUP). If the lower limb muscles contracted lightly, single MUPs were captured. 10-15 MUPs were collected per mouse, and the average MUP duration and amplitude were analyzed.

### Cerebrospinal fluid (CSF) collection, arsenite administration, AAV retro-orbital injection in rodents

Mouse and rat CSF extractions were performed as previously described (Liu and Duff, 2008). CSF solutions were collected, mixed with SDS loading buffer and incubated for 10 min at 95°C. Arsenite (1mg/ml in ddH_2_O) was injected intragastrically (6mg/kg body weight) as previously described (Zhang et al., 2020). For AAV retro-orbital injection, mice were anesthetized by isoflurane and AAV particles expressing our target genes (100μl) were injected into the retro-orbital sinus (Prabhakar et al., 2021). One month after injection, the mice were used for analysis.

### Human subjects

For human samples, the subjects provided written informed consent for the study. The collection of samples (including blood and CSF) and clinical information was approved by the ethics committees at Peking University Third Hospital (IRB00006761-M2020461), and the use of samples was approved by the ethics committees at Tsinghua University (THU01-20230185).

### Cell culture, lentivirus packaging, lentiviral infection, and KO cell lines

HEK293FT, Hela, and U2OS cells were maintained in DMEM (Coring) with 10% FBS at 37°C with 5% CO_2_. For lentivirus packaging, HEK293FT cells were seeded in culture medium in 10-cm culture dishes. When a ∼90% confluency was reached, cells were co-transfected with VSVG (10μg), pxPAX2 (15μg), and pLJM1-EGFP (Addgene) or pLentiCRISPRv2 (Addgene) or pLenticas9-Blast (Addgene) (20μg) using PEI (Sigma). The transfection medium was replaced with fresh growth medium 5-6 hours after transfection. The medium was harvested 72 hours after transfection and centrifugated at 20,000rpm for 2 hours at 4°C to obtain enriched lentivirus pellets. The pellets were resuspended with 100μl DPBS (Dulbecco’s phosphate-buffered saline) and stored at - 80°C. HEK293FT, Hela, or U2OS cells were infected with the appropriate lentivirus and selected with 2μg/ml puromycin or 10μg/ml blasticidin starting from 3 days post-infection, for at least one week. Puromycin- or blasticidin-resistant cells were selected for further analysis. To generate *CCDC22*, *SNX17*, or *VPS35* knock-out single clone cell lines, stable cell lines were seeded in 48-well plates by dilution and cultured for 12-18 days. The top 3 clones were selected for further analysis based on the monoclonality and knock-out efficiency by PCR validation and western blot confirmation.

### Cortical and motor neuron culture

Cortical neurons were cultured as previously described (Guo et al., 2023). Cell culture dishes were coated with Poly-D-Lysine (Sigma) overnight at 37°C. Postnatal day 0 mouse cortices were dissected in HBSS (Invitrogen) and digested with trypsin in HBSS (Invitrogen) for 15 mins at 37°C. DNAse I (Sigma,10μg/ml) was added in the last 10 min of the digestion phase. Subsequently, Neurobasal medium (Invitrogen) with 10% FBS was added to terminate the trypsinization process. The resulting cortical tissues were dissociated by gentle trituration to obtain a homogenous cell suspension and passed through a 70μm cell strainer. Cells were pelleted by centrifugation at 1000g for 10 mins and resuspended in Neurobasal medium supplemented with 10% FBS and GlutaMAX^TM^ (Invitrogen). Cells were plated in a 24-well plate (1 × 10^5^ cells/ml, 500 μl/well). After 5-6 hours of incubation (5% CO_2_, 37°C), the medium was replaced with Neurobasal medium supplemented with B27 (Invitrogen) and GlutaMAX^TM^. Motor neuron culture was performed as previously described (Zhang et al., 2020).

### Protein expression and purification

Recombinant human TRIM72 was expressed in E. coli BL21 strains at 37°C. Once the optical density (OD) reached 0.4, cells were induced with 0.5 mM IPTG and harvested 4 hours post-induction. The cells were collected, washed with ice-cold PBS, and lysed in BC150 buffer (20 mM Tris, 150 mM NaCl, 10 mM imidazole, 1 mM DTT, 5 mM PMSF). The protein was purified by binding to Ni-NTA affinity beads in BC150 buffer at 4°C for 2 hours, followed by elution with elution buffer (20 mM Tris, 150 mM NaCl, 250 mM imidazole, 1 mM DTT, 5 mM PMSF). Further purification was performed using a Superdex 200 Increase column (GE Healthcare). The purified protein was flash-frozen in liquid nitrogen and stored at -80°C for subsequent analysis.

### RNA extraction, real-time PCR, and RNA sequencing

Total RNA was extracted from cells and tissues using TRIzol reagent (Thermo Fisher Scientific) according to the manufacturer’s instructions. For real-time PCR, 1 μg of total RNA was reverse transcribed into cDNA using M-MLV reverse transcriptase (Promega) and oligo (dT) primers.

For RNA sequencing, total RNA was extracted from mouse spinal cords wth TRIzol. RNA quality was analyzed by an Agilent 2100 Bioanalyzer. mRNA was isolated with mRNA Magnetic Isolation Module (NEB) and then converted into cDNA with First Strand Synthesis Module (NEB). The cDNA was fragmented using a Covaris S2 system and purified with AMPure XP beads (Beckman Coulter). Library preparation was completed following the manual provided by the NEB Next Ultra II Directional RNA Library Prep Kit. Libraries were prepared from three biological replicates of animals. The constructed libraries underwent quality control (QC) analysis on a Bioanalyzer and were sequenced on an Illumina HiSeq instrument.

### Immunoprecipitation and western blot

For the immunoprecipitation assay, cultured cells were lysed with IP buffer (50mM Tris-HCl, pH 7.5, 150mM NaCl, 1% Triton X-100, 1mM EDTA, pH 8.0) supplemented with a protease inhibitor cocktail for 30 minutes on ice. The supernatant was incubated with anti-FLAG M2 magnetic beads (Sigma) overnight at 4°C. The beads were washed five times with the IP buffer, and the immunoprecipitated proteins were eluted with 2 ×SDS loading buffer for 10 minutes at 95°C.

For western blots, total protein in tissues or cells was extracted using RIPA lysis buffer (50mM Tris-HCl, pH 8.0, 150mM NaCl, 1mM EDTA, 0.1% SDS, 1% TritonX-100, 0.5% Sodium deoxycholate) supplemented with PMSF and a proteinase inhibitor cocktail. The lysate was separated by SDS-PAGE and transferred onto a PVDF membrane (Cytiva). Membranes were incubated with diluted primary antibodies overnight at 4°C, followed by incubation with HRP-conjugated secondary antibodies (GE Healthcare) 1 hour at room temperature.

The following primary antibodies were used for western blot: anti-FUS (Zhang et al., 2020), anti-TRIM72 (a kind gift from Dr. Jianjie Ma), anti-CCDC22 (Sigma), anti-ALIX, anti-ATP5A, anti-GFP), anti-Histone3, anti-TSG101 (Abcam), anti-Flag, anti-Myc (Abmart), anti-EEA1, anti-GAPDH, anti-Golgin97, anti-REEP5, anti-SNX17, and anti-TUBULIN (Proteintech). The secondary antibodies (GE Healthcare) were applied and images were analyzed by using the Fiji ImageJ to calculate the integrated intensities.

### LC-MS/MS

Stable Hela/293FT cell lines expressing Flag-TRIM72 were generated by lentiviral infection. Electrophoresis was performed to allow the eluted samples to enter the upper portion of the SDS-PAGE gels. The gels were stained by Coomassie blue and excised. LC-MS/MS was performed as previously described (Meng et al., 2023). The gel slides were reduced, alkylated and digested with trypsin (Promega). Digested samples were quenched with 10% trifluoroacetic acid and peptides were extracted and dissolved in 0.1% trifluoroacetic acid. Purified peptides were separated with a C18 column (75 μm inner diameter, 150 mm length, 5 μm, 300 Å) and connected to an Orbitrap Fusion Lumos mass spectrometer (Thermo Fisher Scientific). Raw data were compared with the Uniport human database using the SEQUEST search engine with Proteome Discoverer software (version PD1.4, Thermo Scientific^TM^). Peptide spectrum match (PSM) was calculated by Percolator (Proteome Discoverer), and only peptides with FDR less than 0.01 were included for further analysis. Peptides that were only assigned to a given protein group were considered “unique”. FDR was set to 0.01 for protein identification.

For quantitative proteomics analysis (Jiang et al., 2019), CSF samples and the standard reference peptides (155-EHQLVEVEETVR-166, and 311-EVECSEQK-317) were reduced, alkylated, and digested with trypsin. Peptides were purified on SDB-RP StageTips. CSF samples and standard reference peptides were desalted and labeled by TMT10plex^TM^ (Thermo Fisher Scientific), separately. The reference peptides were labeled with TMT10 126. CSF samples were grouped into two groups and labeled with the other six isobaric compounds of TMT10. TMT-labeled peptides were mixed and desalted using Sep-Park C18 cartridges and analyzed on the Q-Exactive HF-X mass spectrometers (Thermo Fisher Scientific). Data analysis follows the steps described above.

### Immunostaining and RNA *in situ*

Brain and spinal cord were embedded in paraffin and cut into 5μm sections. Sections were deparaffinized thoroughly with xylene solvent and hydrated in a graded series of alcohol solutions. Deparaffinized sections were boiled in 10mM citrate buffer, pH 6.0, three-five times in a microwave box for antigen retrieval. After blocking with 1% BSA in PBS-T for one hour at room temperature, sections were incubated with primary antibodies, including anti-FUS (Zhang et al., 2020), anti-G3BP1(Abcam), anti-G3BP2 (Abcam), anti-eIF3η (Santa Cruz), and anti-ChAT (Millipore) overnight at 4°C. After PBS washes, Alexa Fluor-conjugated secondary antibodies (Invitrogen, 1:1000) were applied and images were captured by Nikon A1 confocal microscope. The maximum intensity projection images were assessed for fluorescence intensity and mean intensity analysis by using Fiji ImageJ. The expression level in different groups was reflected by RMI (RMI = mean intensity of given protein divided by that of background).

For RNA *in situ*, mice were perfused with PBS and 4% PFA. Brain and spinal cord tissues were dissected and embedded in OCT (Electron Microscopy Science, Japan) and stored at -80°C. Frozen sections (10μm) were applied for RNA FISH. In details, sections were incubated with *Trim72* probe (RNAscope), *Vglut1* probe (Mm-*Vglut1*-C2) or *Vgat* probe (Mm-*Vgat*-C2). Fluorescent probes were detected with Opal^TM^ 570 or Opal^TM^ 690 reagents (AKOYA biosciences). Subsequently, probed sections were blocked with 1% BSA in PBS-T (0.3% TritonX-100) at room temperature for 1 hour and incubated with primary antibodies, including anti-ChAT (Millipore), anti-GFAP (Cell Signaling Technology), and anti-IBA1 (Wako), overnight at 4°C. Secondary antibodies (Invitrogen) were incubated for 1 hour at room temperature. Images were acquired with the Nikon A1 confocal microscope and *Trim72* mRNA level was quantified by the number of *Trim72* positive puncta per cell.

### Immunogold Electron Microscopy

Samples were embedded using LR Gold resin and UV-polymerized at -20°C for 24 hours, followed by polymerization at room temperature for 72 hours. Ultrathin sections of 70 nm were cut and collected on 100 mesh grids coated with carbon-coated Formvar film. The grids were washed with phosphate buffer (PB) and then blocked with blocking buffer (containing 2% BSA and 0.2% cold fish gelatin in PB) at room temperature for 20 minutes. They were incubated with the primary anti-GFP antibody (Abcam, ab6556, 1:50 dilution) in blocking buffer at room temperature for 1 hour, followed by five washes with 0.2% cold fish gelatin in PB. Subsequently, the grids were incubated with a 10 nm gold-labeled secondary antibody (Sigma, G7402, 1:20 dilution) in blocking buffer at room temperature for another hour, followed by three washes with PB containing 0.2% cold fish gelatin, and two additional washes with PB. The grids were then fixed in 0.5% glutaraldehyde for 10 minutes and washed three times with ddH2O. The sections were stained with 4% uranyl acetate (UA) for 5 minutes. After air drying, they were examined using a transmission electron microscope (TEM; H-7800) at 80 kV.

### Correlative Light and Electron Microscopy (CLEM) procedure

Immediately after imaging, an equal volume of pre-warmed 4% PFA was added into the cultured cells. Brightfield images were captured using a 10× objective, while fluorescent images were captured using a 100× oil objective. The positions of the photodamaged cells were recorded. Cells were fixed with 2.5% glutaraldehyde for 2 hours at room temperature and then dehydrated in a gradual ethanol gradient ethanol (50-100%) for 2 minutes at each gradient. Subsequently, cells were infiltrated and embedded in resin (SPI-Pon 812 kit) and polymerized at 60°C for 48 hours. Ultrathin sections (70 nm) were cut using a diamond knife and collected on slot grids. Sections were stained with uranyl acetate (UA) and lead citrate, air-dried, and examined using a transmission electron microscope (TEM, H-7800) at an acceleration voltage of 80 kV.

### Exosome purification

For exosome purification experiments, FBS for cell culture was centrifuged at 120,000g for 12 hours at 4°C, and the supernatant was collected and passed through a 0.22μm sterile filter, to remove the background exosome in FBS. Cell culture media was collected 60 hours post-seeding and centrifuged at 1,000g for 10 min at 4°C. The supernatant was collected and passed through a 0.22μm sterile filter. The filtered supernatant was centrifuged at 120,000g for 70 min at 4°C, and the supernatant was removed and the precipitated pellet was collected. 20mM HEPES (pH7.4, stored at 4°C) was used to make a serial dilution of sucrose solutions from 0.25M to 2.25M (9 sucrose concentrations with 10ml each). The pellet was resuspended in the 0.25M sucrose solution and overlaid with the serially diluted sucrose stock at 430μl per concentration. This layered column was centrifuged at 210,000g for 4 hours at 4°C and collected into 1.5ml centrifuge tubes by layer. The fractions were collected without mixing the gradient with 430μl per fraction. This collected fraction can be analyzed immediately or stored at -80°C for short-term storage. To concentrate the collected fractions, 900μl DPBS was added, mixed thoroughly and centrifuged at 120,000g for 1 hour at 4°C, followed by collection of the pellet.

For exosome purification from patient CSF, the collected CSF was initially centrifuged at 1000g for 10 minutes at 4°C. The supernatant was carefully transferred into Eppendorf tubes (1.4 ml/tube) and subjected to a second centrifugation step (120,000g, 4°C). Subsequently, the supernatant was discarded, and the resultant pellet, containing the concentrated exosomes, was resuspended in 40 μl with DPBS.

### Reactive Oxygen Species (ROS) measurement and CCK-8 assay

ROS levels were measured using dihydroethidium (DHE, Invitrogen) and MitoSOX (Invitrogen). Cortical neurons were cultured in 24-well chamber slides for 12-14 days. Subsequently, neurons were incubated with DHE (20μM in medium) or MitoSOX (5μM in medium) for 20 mins at 37°C and fixed with 4% PFA for 10 min at room temperature. Images were captured by a Nikon A1 confocal microscope. DHE fluorescence was excited at 535nm, and the emission was collected at 610 nm. MitoSOX fluorescence was excited at 510nm, and the emission was collected at 580 nm. The mean intensity of DHE or MitoSOX fluorescence was assessed by Fiji ImageJ software. The ROS level was expressed as the RMI of DHE or MitoSOX fluorescence in different groups. The RMI = mean intensity of DHE or MitoSOX fluorescence divided by the mean intensity of the background. For lentiviral infection, cortical neurons were seeded in 24-well chamber slides and cultured for 3-5 days. 10μl purified lentivirus was added per well, and the neurons were subjected to DHE staining on 12-14DIVs.

For CCK-8 assay, 293FT cells were seeded in 96-well plates at a density of 1.5 × 10^3^ cells per well. Arsenite (Sigma) was added to each well at a concentration of 0.125mM, 0.25mM, or 0.5mM and washed after 2 hours of incubation at 37°C. 10μl CCK-8 solution (Yeasen) was added per well and incubated for 2 hours at 37°C. The optical density (OD) of each well was measured using a microplate reader with an excitation wavelength of 450nm. The CCK-8 value of 293FT cells was calculated by computing the mean value of 4-5 biological replicates.

### Photodamage assay

Cells were cultured at 37°C, 5% CO2 for 20hours and stained with PI dye (Beyotime). The Nikon A1HD25 confocal microscope with a 100×oil objective was used to select and position the selected cells in the center of the field. The focal plane was adjusted to focus on the bottom of the cells. For photodamage, 405nm and 488nm lasers at 100% power were applied to irradiate the selected region in a 4s loop and repeated for 4 loops. The selected region should be in the cytoplasm but not in the nucleus or axon. Images were acquired every 10s for 10 mins and processed and analyzed using the Fiji ImageJ. **Mitochondrion labeling in cultured cortical neuron in microfluidic devices** Mouse embryonic day 18 (E18) cortical neurons were cultured in microfluidic devices as previously described (Wang et al., 2016). Briefly, microfluidic devices were coated overnight with 0.5 mg/ml Poly-L-Lysine in Borate buffer (1M, pH 8.5) across all chambers. Cortical neurons were dissected and seeded into the soma chambers of the microfluidic devices. On DIV8, the culture medium was removed from the axon chambers and replaced with MitoTracker Deep Red FM (Cell Signaling) solution diluted in culture medium for 1 hour at 37°C, 5% CO2 to label mitochondria. Prior to imaging, the culture medium was replaced with Neurobasal minus phenol red imaging medium (Thermo Fisher Scientific).

For immunofluorescence microscopy, microfluidic devices were carefully removed and primary cortical neurons were fixed on 8DIVs and immunostained as previously described (Wang et al., 2016). In brief, neurons were fixed with 4% PFA, rinsed three times with PBS, and incubated with a blocking buffer (0.1% saponin, 1% BSA, 0.2% gelatin in PBS) for 1 hour at room temperature. Cells were incubated with rabbit anti-FUS antibody (self-made, 1:500) at 4°C overnight and Alexa Fluor 568 goat anti-rabbit secondary antibody (Thermo Fisher Scientific) for 1 hour at room temperature. Dishes were mounted with mounting medium (Sigma-Aldrich) and imaged with a Nikon TI2-E inverted microscope equipped with a Yokogawa spinning confocal disk head (CSU-W1 dual camera).

### Live imaging of mitochondrion movement

Live imaging of mitochondrion movement was conducted using a Nikon TI2-E inverted microscope equipped with a Yokogawa spinning confocal disk head (CSU-W1 camera) and a 63 × 1.4 NA/219.15 µm WD/0.1826 µm/pixel (1,200 × 1,200) objective. Neurobasal minus phenol red imaging medium (Thermo Fisher Scientific) was used during imaging sessions. Time-lapse images were acquired continuously for a minimum of 15 min to capture axonal trafficking of mitochondria. Subsequently, time-lapse images were exported to ImageJ2 (version 1.54f, NIH) for analysis. Mitochondrial movement analysis was performed using the Trackmate Plugin of ImageJ2, as previously described (Wang and Meunier, 2022). Raw live stacks of the mitochondrial channel were stabilized and converted to binary images using the Threshold function. Mitochondrial movement was traced using the ‘Trackmate’ plugin, and the mean speed of each track was recorded and exported. For further analysis, tracks spanning over 2 frames were selected, and mitochondria with a track mean speed exceeding 0.005 μm/s were classified as the mobile fraction. All data were analyzed and illustrated using GraphPad Prism.

### Automatic analysis of FUS aggregates

Z-stack images of axons were exported to ImageJ2 and analyzed using the ‘Analyze Particles’ function. FUS aggregates were extracted based on the following criteria: circularity > 0.8 and area > 0.5 μm². The number of selected FUS puncta was normalized to the total axonal area of each region of interest (ROI). Additionally, the average size of extracted FUS puncta was calculated within each ROI. All data were analyzed and illustrated using GraphPad Prism.

### Anti-drug antibody (ADA) measurement in macaque

Macaque serum samples were collected, and the levels of total IgG against AAV9 were determined by ELISA. Briefly, 96-well plates were coated with AAV9 capsids (1E10/mL) in carbonate coating buffer (antigen-positive [ag+]) and carbonate coating buffer only (antigen-negative [ag-]) for each sample and incubated at 4°C overnight. Samples were washed with PBS-T (PBS containing 0.05% Tween-20) and blocked with blocking buffer (PBS containing 5% skim milk and 2% BSA) at room temperature for 1 hour. Two-fold serial dilutions (starting at 1:50) of serum samples in blocking buffer were added to the plates and incubated at 37°C for 1 hour. Plates were then incubated with HRP-conjugated anti-human IgG at 37°C for 1 hour. After washing, plates were developed with TMB at room temperature for 5 minutes. The reaction was stopped, and OD was measured by a spectrophotometer at 450 nm. Serum total anti-AAV IgG levels were determined as the ADA titer, which was calculated as the minimum dilution factor from the formula > 2: [OD450 (sample) - OD450(Blank)] / OD450 (Blank).

### AAV9 shedding

The viral genome of each sample was extracted using the TIANamp Virus DNA/RNA Kit (TIANGEN). 100μl RNAse-free ddH2O was used for elution of plasma, saliva, and fecal samples, and 50μl for urine samples. PCR was performed using the Taq Pro HS Universal U+ Probe Master Mix (Vazyme). Each reaction consists of 2 ×Taq Pro HS Universal U+ Probe Master Mix (10ul), Primer F (10uM, 0.4ul), Primer R (10μM, 0.4μl), Probe (10μM, 0.2μl), nuclease-free ddH2O (4μl) and template DNA (5μl). The AAV9 copy number in each sample was calculated using a standard curve generated from AAV9 standards with standardized concentrations.

### AAV9 neutralizing antibody measurement in patient

Serum or CSF samples were treated for 30 minutes at 56°C, then centrifuged at 2,000g for 10 minutes at 4°C. The supernatant was diluted with DMEM, and AAV9 viral particles expressing luciferase were added to each dilution gradient. After incubation at 37°C, the mixture was added to HEK293FT cells and luciferase activity was measured using the Stable-Lite^TM^ (Vazyme). The half-maximal inhibitory concentration (IC50) of the samples was calculated using regression analysis.

### TRIM72-Commander complex structure prediction

The protein complex, including CCDC22 (386-627), CCDC93 (377-631), VPS29, VPS26C, VPS35L, and TRIM72 dimers, were predicted by AlphaFold3. The structure of COMMD1-10/CCDC22/CCDC93 protein complex was reported previously (Healy et al., 2023). The two complex structures were merged by overlaps between CCDC22 and CCDC93 and optimized by PyRosetta to generate the final complex structure.

## Acknowledgements

We wish to express our gratitude to all the staff of the Laboratory Animal Research Center at Tsinghua University for their handling and care of mice. We greatly appreciate the assistance provided by the Imaging Core Facility and the Technology Center for Protein Sciences (Tsinghua University) with Nikon A1 confocal imaging, as well as the imaging support from the SLSTU-Nikon Biological Imaging Center. We would like to acknowledge Dr. Ying Li from the Cryo-EM Facility of Tsinghua University, a branch of the National Protein Science Facility, for her help with exosome negative staining, immunogold electron microscopy, and CLEM. This work was supported by the Tsinghua-Peking Joint Center for Life Sciences, the lDG/McGovern Institute for Brain Research (Brain+X project), National Natural Science Foundation of China (82471447, 81974197), Beijing Municipal Natural Science Foundation (L242033;7222215), Beijing Physician Scientist Training Program (BJPSTP-2024-03), the China Postdoctoral Science Foundation (2020TQ0179), the National Science and Foundation for Young Scholars of China (82101495), and the National Key Research and Development Program of China (2022YFA1303003). We would also like to extend our gratitude to Dr. Jianjie Ma at the University of Virginia Medical School for sharing the TRIM72 protein and antibodies. Our thanks also go to Dr. Jing Xu from the Institute of Neuroscience, Chinese Academy of Sciences, for sharing the FUS KI rats, as well as to patient C132 for participating in the SNUG001 clinical trial. We thank the National Key Laboratory of Difficult, Severe and Rare Diseases and the Center for Regeneration, Aging and Chronic Diseases (School of Basic Medical Sciences, Tsinghua University). We thank Shan Shan Shi for her contributions to manuscript writing.

## Author contributions

D.F. and Y.J. designed and supervised the experiments. X.Z. performed the biochemical experiments, immunofluorescence staining, and data analysis, and characterized the animals. N.S. analyzed the TRIM72 expression and performed the RNA *in situ* experiments. Z.C., H.C., and X.H. conducted EMG measurements in mice. X.Z. and N.S. performed ROS measurement. X.Z., N.S., C.Y., P.Z., and X.J. performed the co-immunoprecipitation (co-IP) experiment. X.Z. and Y.H. carried out exosome purification and the photobleaching assay. X.P. and L.L. generated the *Trim72* KO mouse lines. X.Z. and D.M. analyzed RNA-seq data. W.W., X.Z., and H.G. performed Commander-TRIM72 complex prediction with AlphaFold3. T.W., Q.L., and Y.H. analyzed mitochondrial movement. W.G. coordinated the non-human primate experiments, which were performed by P.Y., X.Q., and J.R. J.H., X.L., and D.F. collected ALS samples and performed patient diagnosis, conceived of and designed the clinical study. L.P. coordinated the clinical research. X.Z., P.Z., C.Y., and N.S. conducted clinical sample analysis. X.Z., J.H., and Y.J. wrote the manuscript.

## Declaration of Interests

Y.J. is a scientific advisor at Beijing SineuGene Therapeutics Co., Ltd. W.G. and L.P. are full-time employees of Beijing SineuGene Therapeutics Co., Ltd. X.Z., and Y.J. have patent applications related to this work.

---

**Figure S1.**
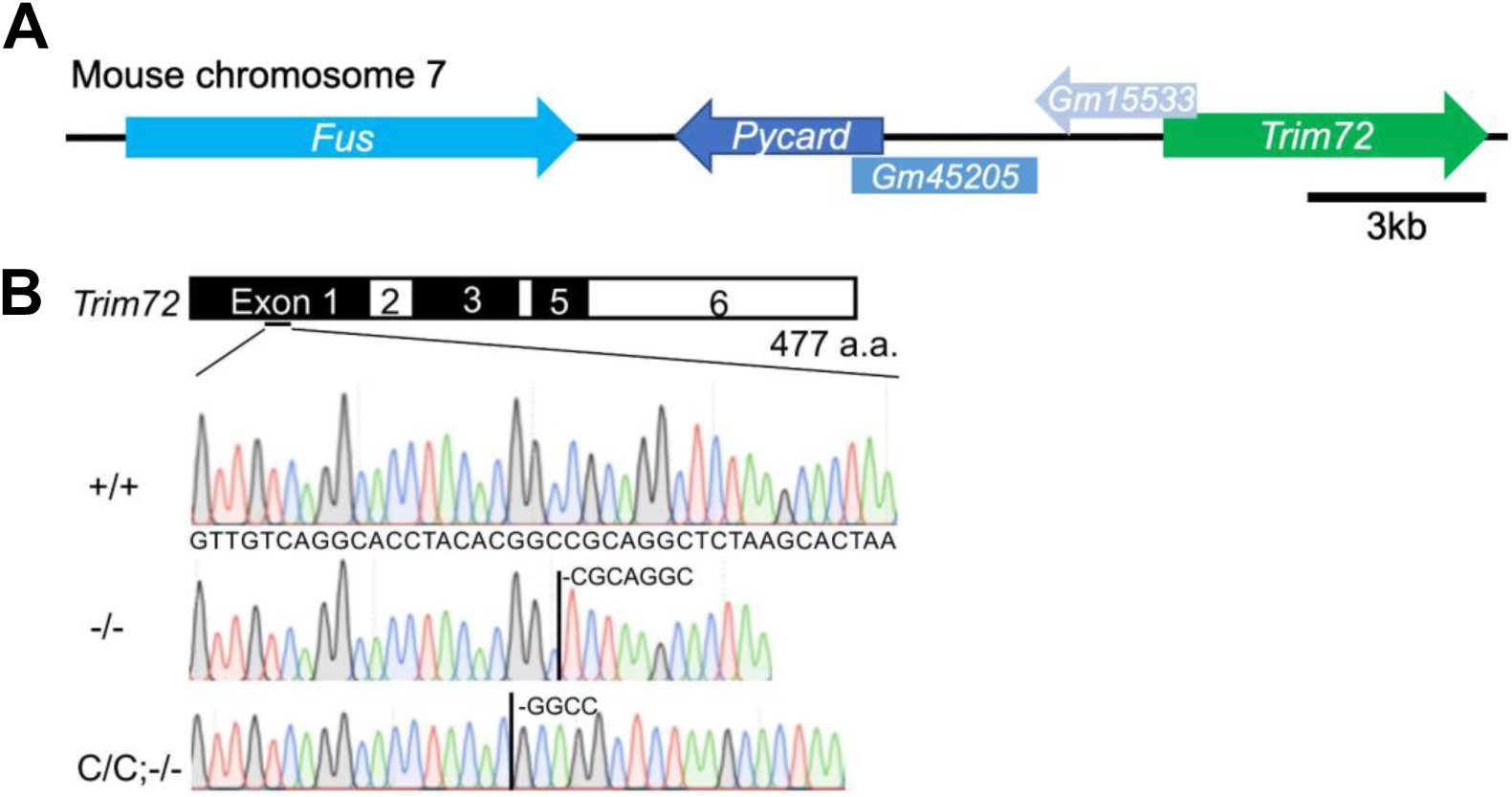
Generation of *Trim72* KO mouse lines by Crispr/Cas9. (A) Chromosomal structure of mouse *FUS* locus. *Trim72* was several kilo-base-pairs away from the *FUS* locus. (B) DNA chromatograms confirmed the genome editing in exon1 of *Trim72*. Due to the close chromosome distance between *Trim72* and *FUS*, we generated *Trim72*_−/−_ in both wildtype (+/+) and FUS-R521C (C/C) backgrounds.

**Figure S2.**
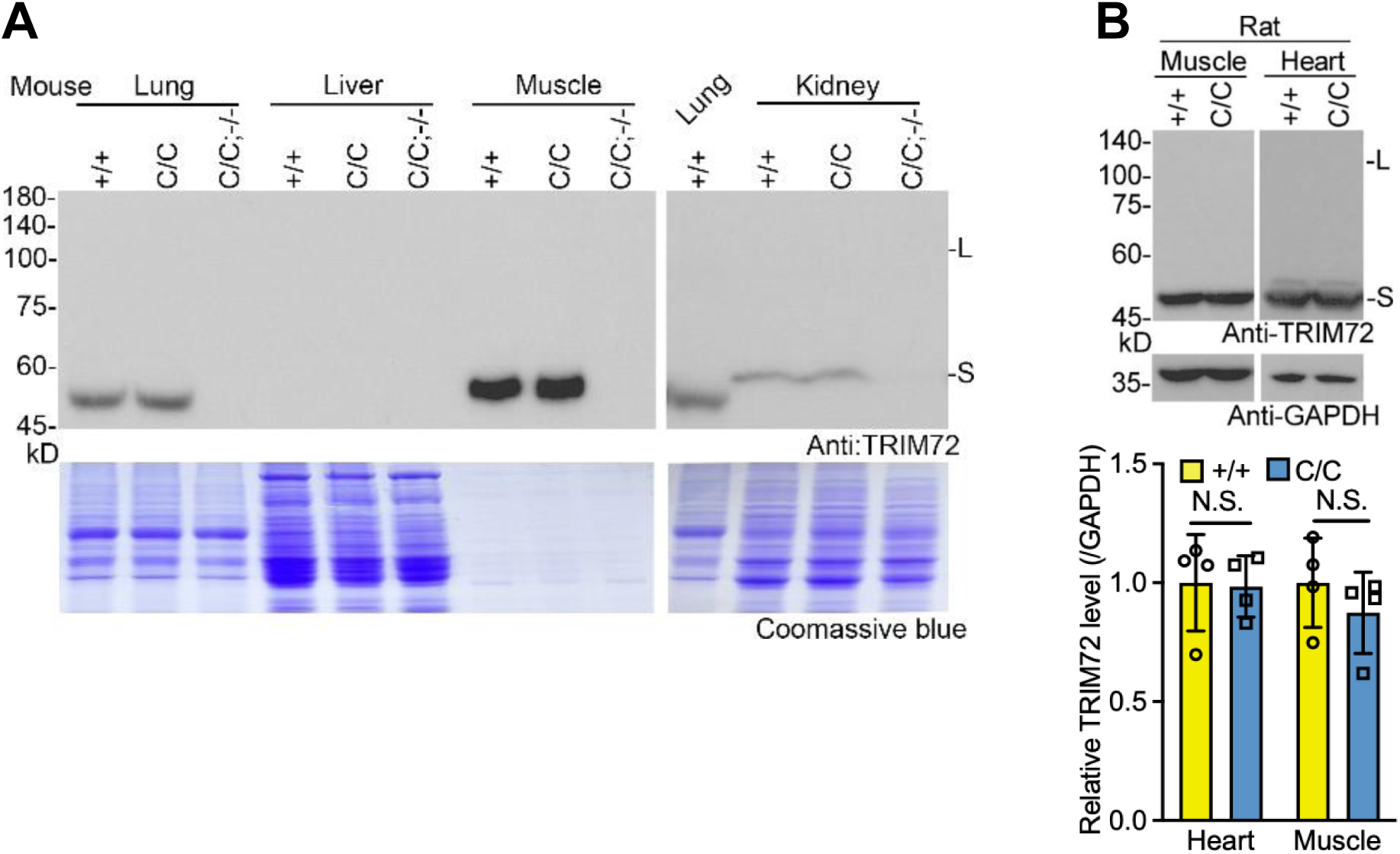
TRIM72 expression in FUS-R521C rodent non-neural tissues. TRIM72 expression in non-neural tissues in mouse (**A**) and rat (**B**) with the indicated genotypes. Total protein amount was visualized by Coomassie blue. Data summary for (**A**) was shown in Figure 1E. In A, *Trim72* knockout mouse with FUS-R521C mutation (C/C;−/−) served as a negative control. In **B**, values were presented as mean ± SD. N.S., no statistical significance (T-Test, SPSS).

**Figure S3.**
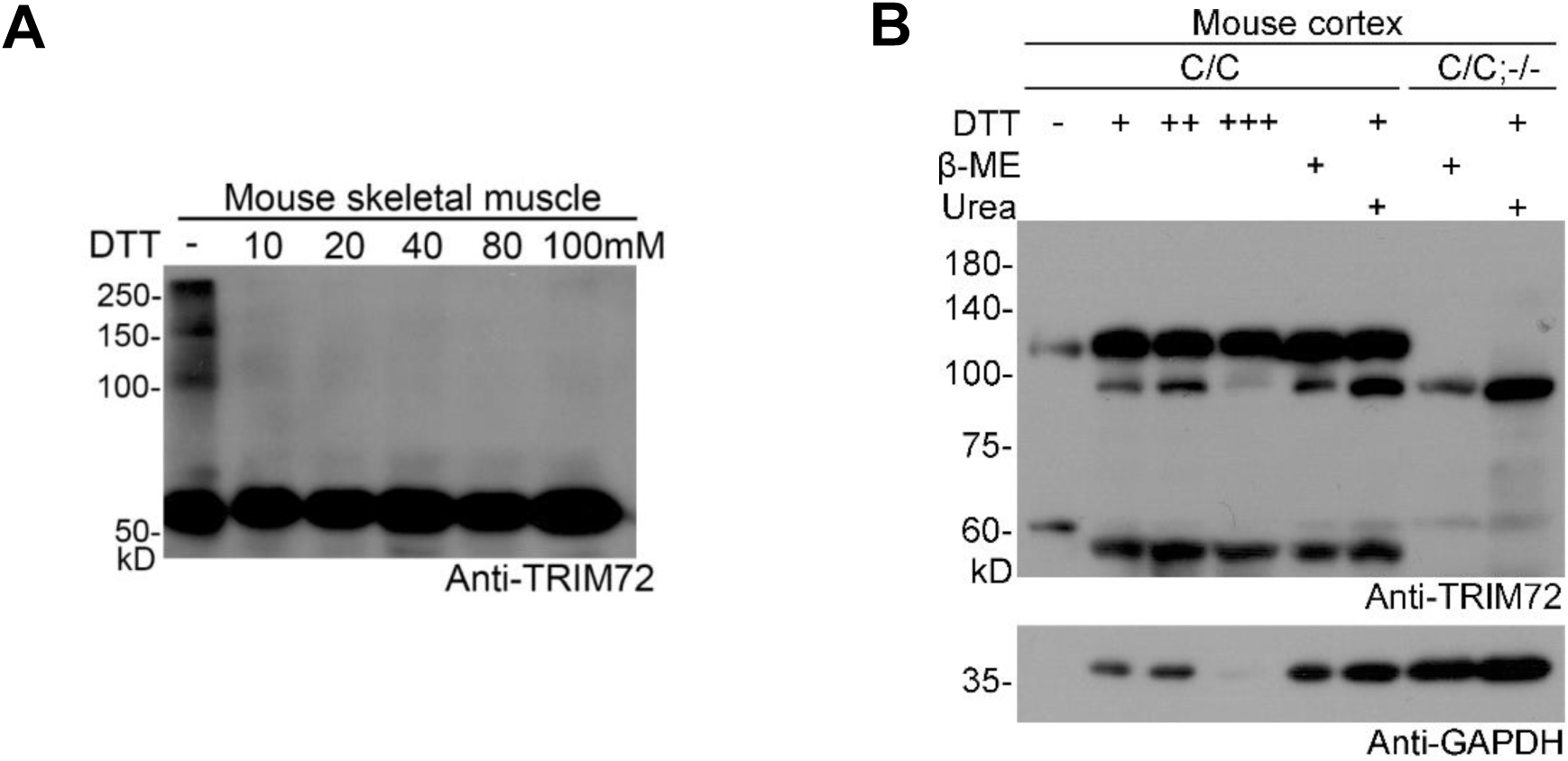
The high molecular weight TRIM72 in FUS-R521C mutant neural tissues is insensitive to dithiothreitol (DTT), urea, 2-mercaptoethanol (β-ME). (A) Consistent with a previous report (PMID: 19043407), the oligomerization of TRIM72 in skeletal muscle is sensitive to DTT at a very low concentration of 10 mM. (B) The high molecular weight TRIM72 (∼110 kD) in the knockin (C/C) mouse cortex is insensitive to DTT, urea, and β-ME. DTT: +, 10; ++, 150; +++, 200 mM; urea: 8 M; β-ME: 200 mM. Mice age, 2 months. *Trim72* knockout mice with FUS-R521C mutation (C/C;−/−) served as negative control. *, non-specific bands.

**Figure S4.**
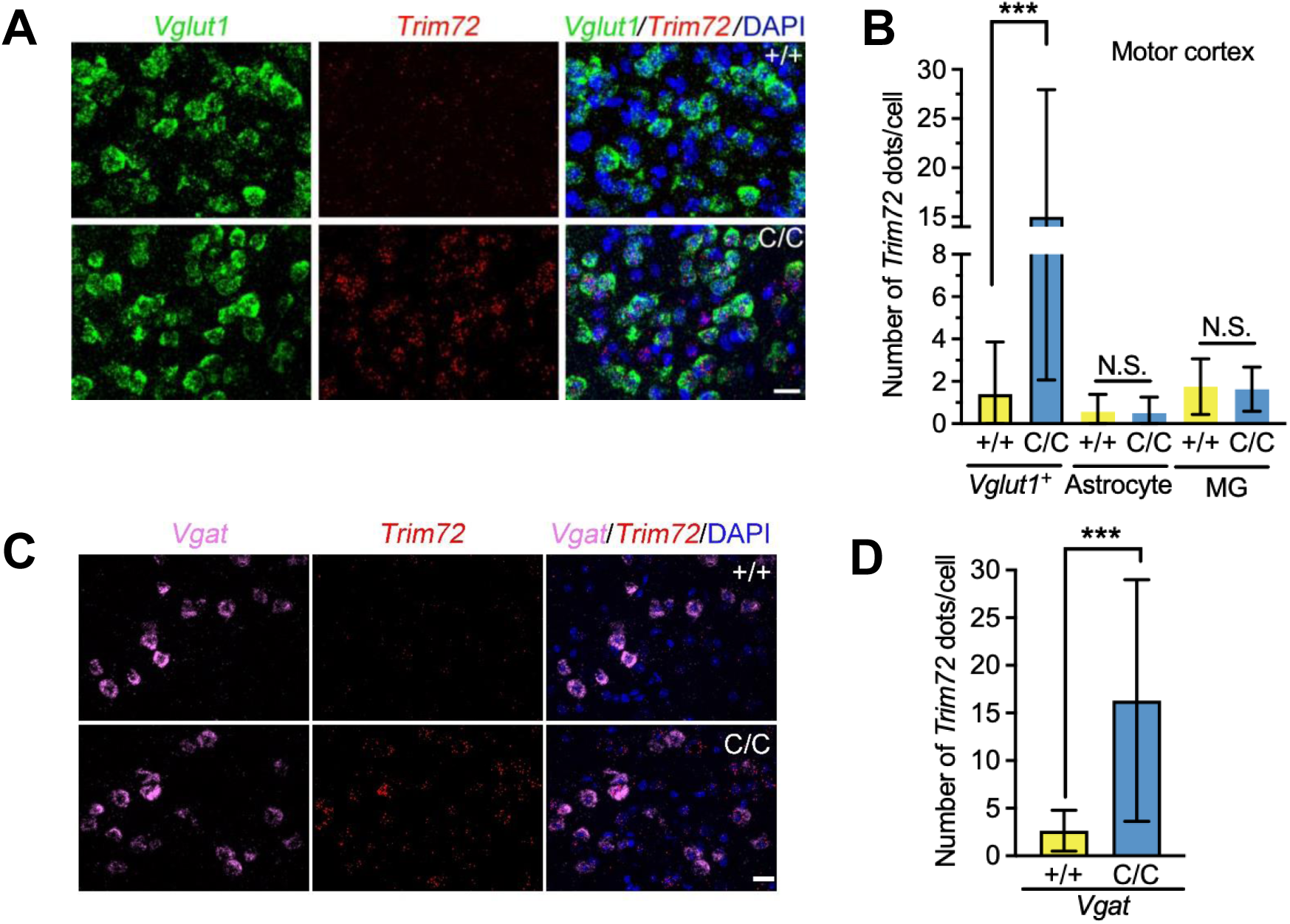
Neuronal upregulation of *Trim72* in FUS-R521C KI motor cortex. (**A**-**D**) *Trim72* RNA *in situ* hybridization was performed in motor cortex with the indicated genotypes. In the motor cortex, excitatory neurons, inhibitory neurons, astrocytes, and microglial (MG) were marked by *Vglut1*, *VGAT*, GFAP, and Iba1, respectively. In **A** and **C**, scale bar, 20 μm. Mice, 3 months of age (n = 3). Data summaries were shown in **B** and **D** and values were presented as mean ± SD. \*\*\**p* <0.001; N.S., no statistical significance (T-Test, SPSS).

**Figure S5.**
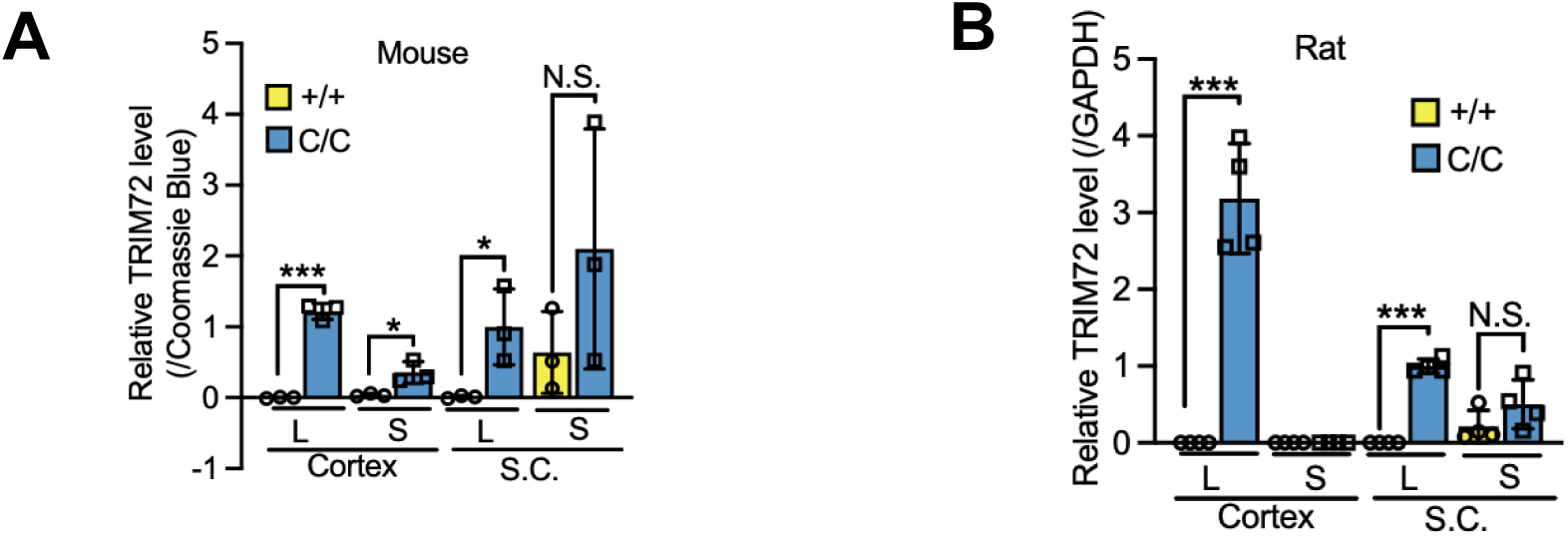
Large and small molecular weight TRIM72 are upregulated in the cortex and spinal cord of rodent ALS models. TRIM72 expression level in the cortex and spinal cord in mice (**A**) and rats (**B**) with the indicated genotypes. Large (L, ∼110 kD) and small (S, ∼55 kD) molecular weight TRIM72 were upregulated in neural tissues derived from mutant animals.

**Figure S6.**
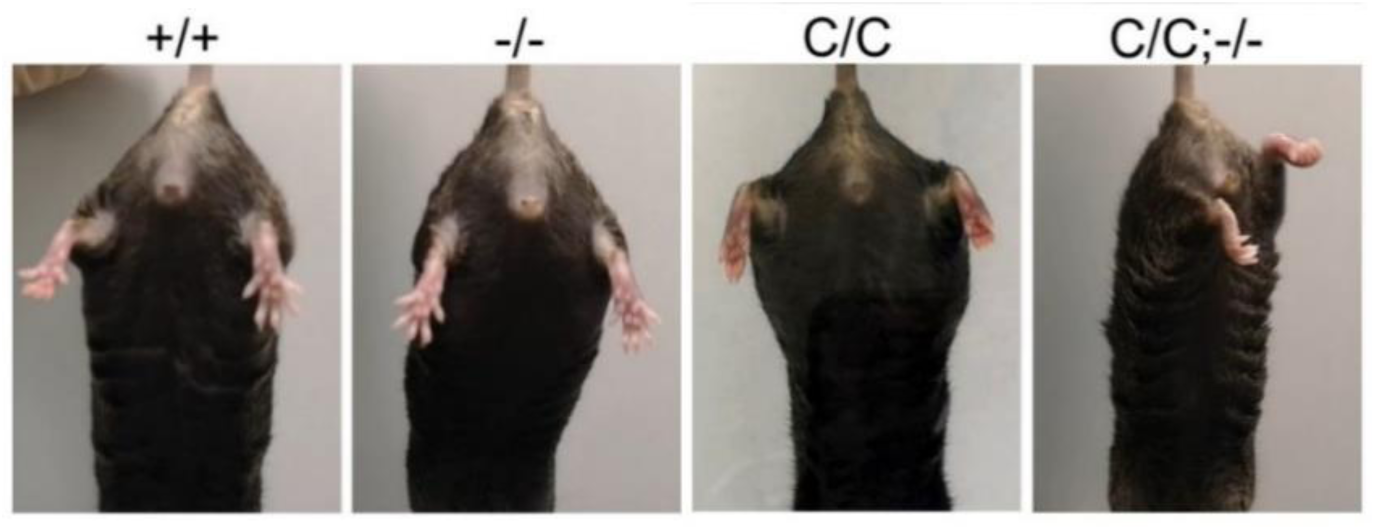
*Trim72 l*oss-of-function accelerates disease progression in FUS-R521C ALS mouse model. Hindlimb extension test of the indicated genotypes at 1.5 year of age. Normal spreading of the hindlimbs was observed in wildtype and *Trim72* knockout (−/−) mice. Hindlimb pullback appeared in C/C;−/− mice.

**Figure S7.**
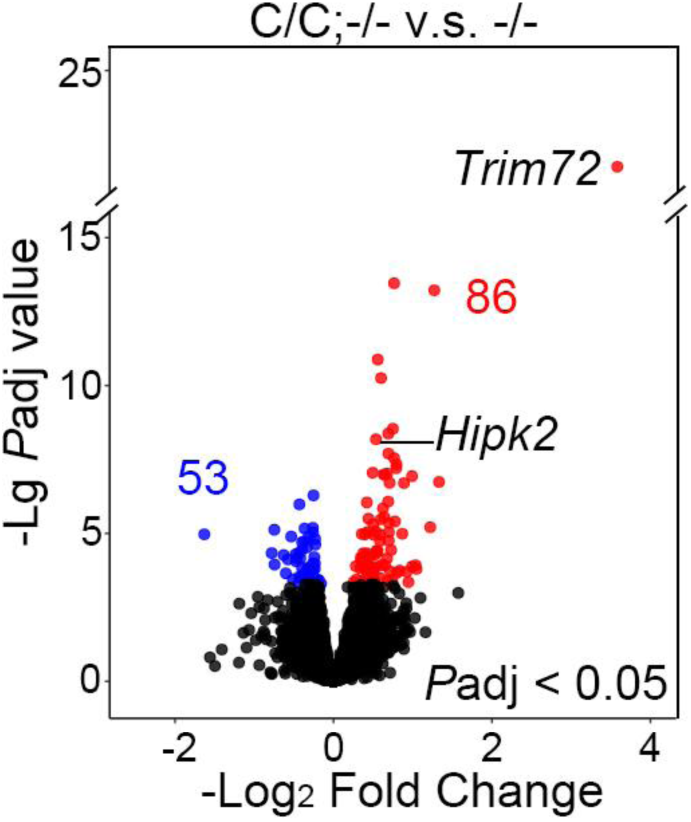
Differentially expressed genes between *Trim72* KO (−/−) and C/C;−/− identified by RNA-seq. Volcano plot of differentially expressed genes between −/− and C/C;−/−. Significantly upregulated genes (86 genes, red) and downregulated genes (53 genes, blue)*, P*adj < 0.05. *Trim72* and *Hipk2* were highlighted.

**Figure S8.**
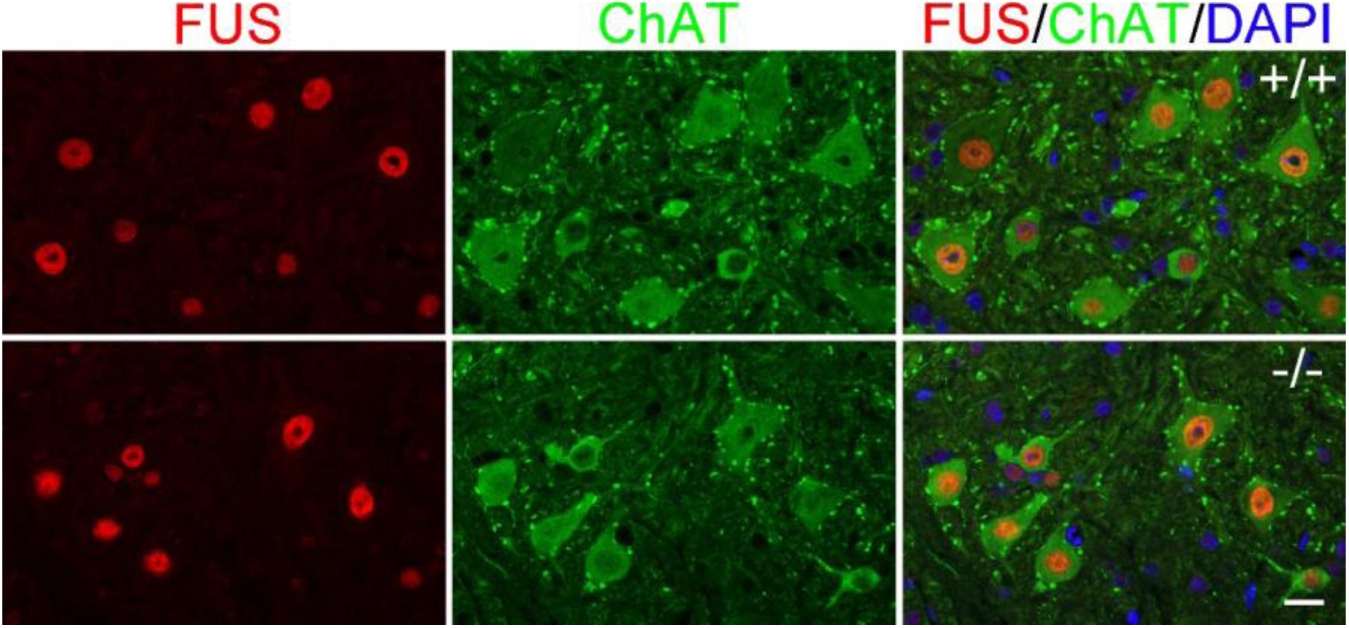
*Trim72* KO does not affect FUS nuclear localization in spinal cord motor neurons. Sections of lumbar 4-5 spinal cords of wildtype (+/+) and *Trim72* KO (−/−) mice were stained with FUS and ChAT antibodies. 1.5 months of age. Scale bar, 20 μm.

**Figure S9.**
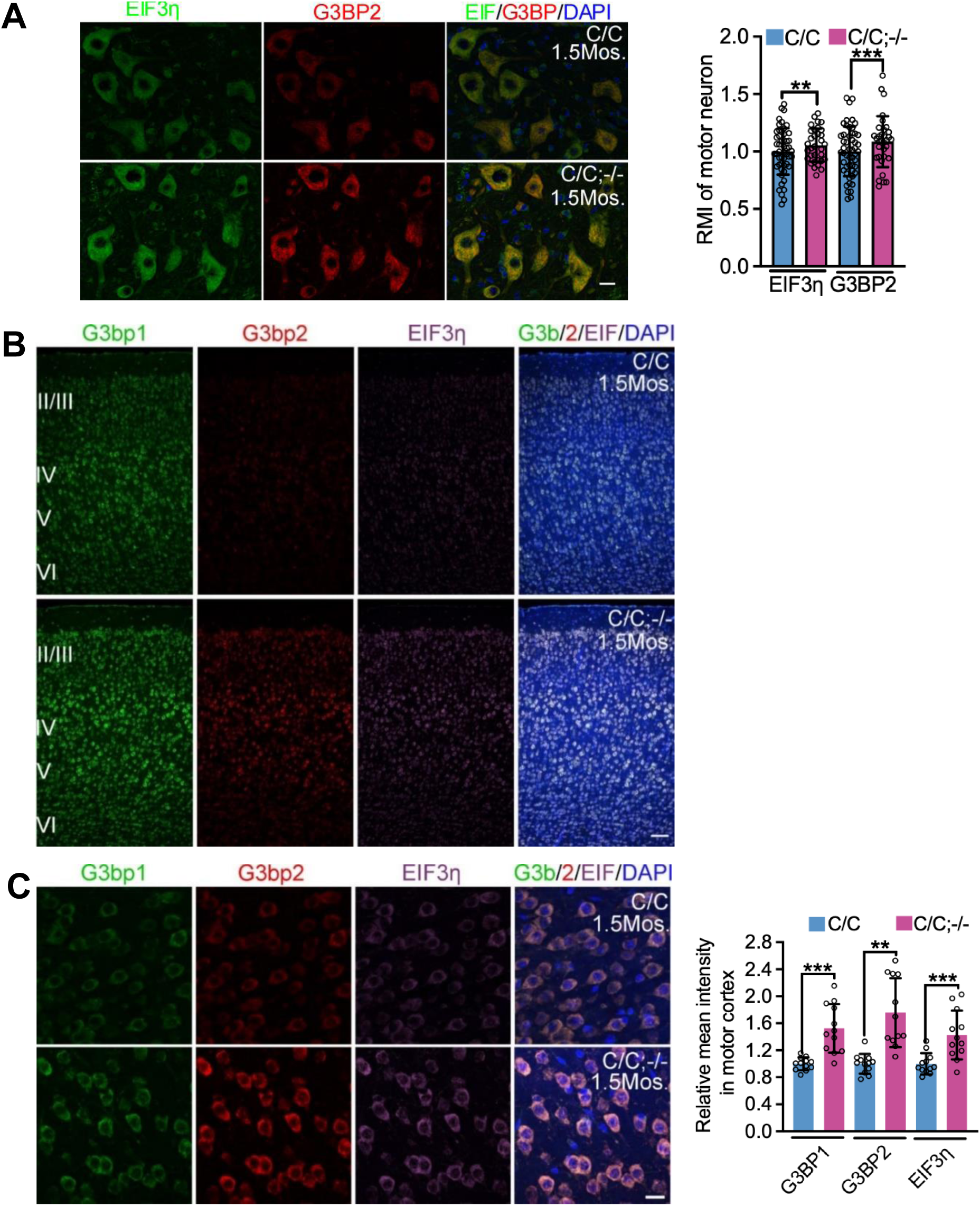
Stress granule proteins were upregulated in C/C;−/− mutant spinal cord and motor cortex. Immunostaining of stress granule proteins, G3BP1, G3BP2, and EIF3η, in C/C and C/C;−/− spinal cord (**A**) and motor cortex (**B** and **C**). In **A**, scale bar, 20 μm; in **B**, 100 μm; in **C**, 20 μm. Mouse, 1.5 months of age (n= 3). Values in **A**, and **C** were presented as mean ± SD. \*\**p* < 0.01; \*\*\**p* < 0.001; T-test, SPSS.

**Figure S10.**
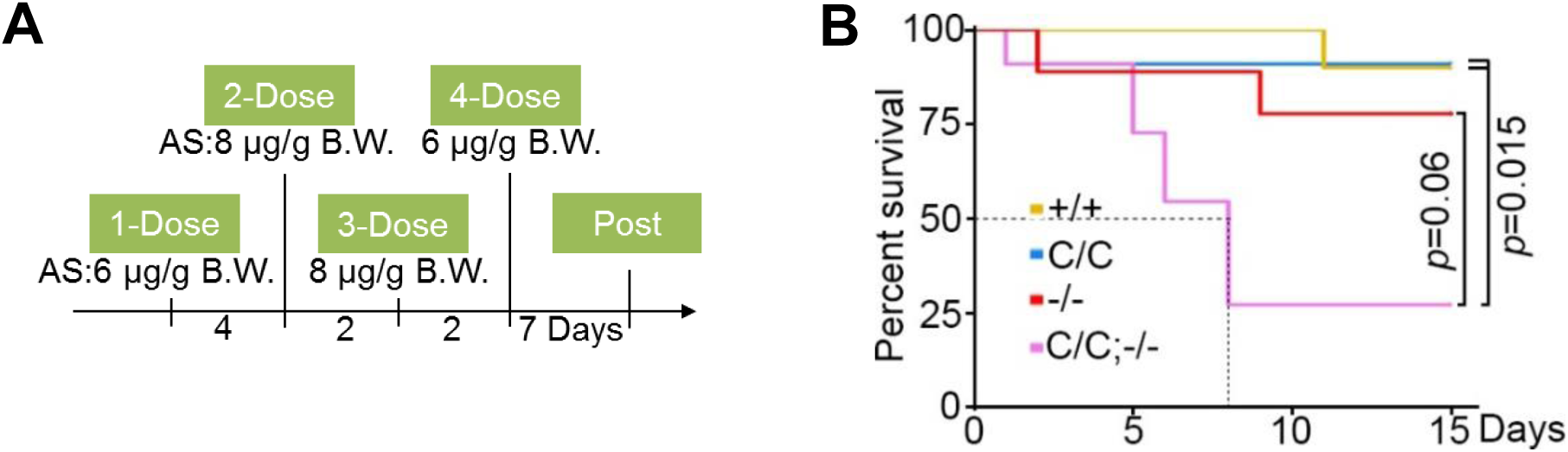
The C/C;−/− mutant mouse is sensitive to AS-induced toxicity. (**A** and **B**) Two-week experimental procedure for AS challenge (**A**). Mice with the indicated genotypes were treated with AS by intragastric administration and the survival curve was plotted (**B**). Mouse, male; age, 2 months; +/+ (n = 10); C/C (n = 11); C/C;−/−, n = 11; −/−, n = 9. B.W., body weight.

**Figure S11.**
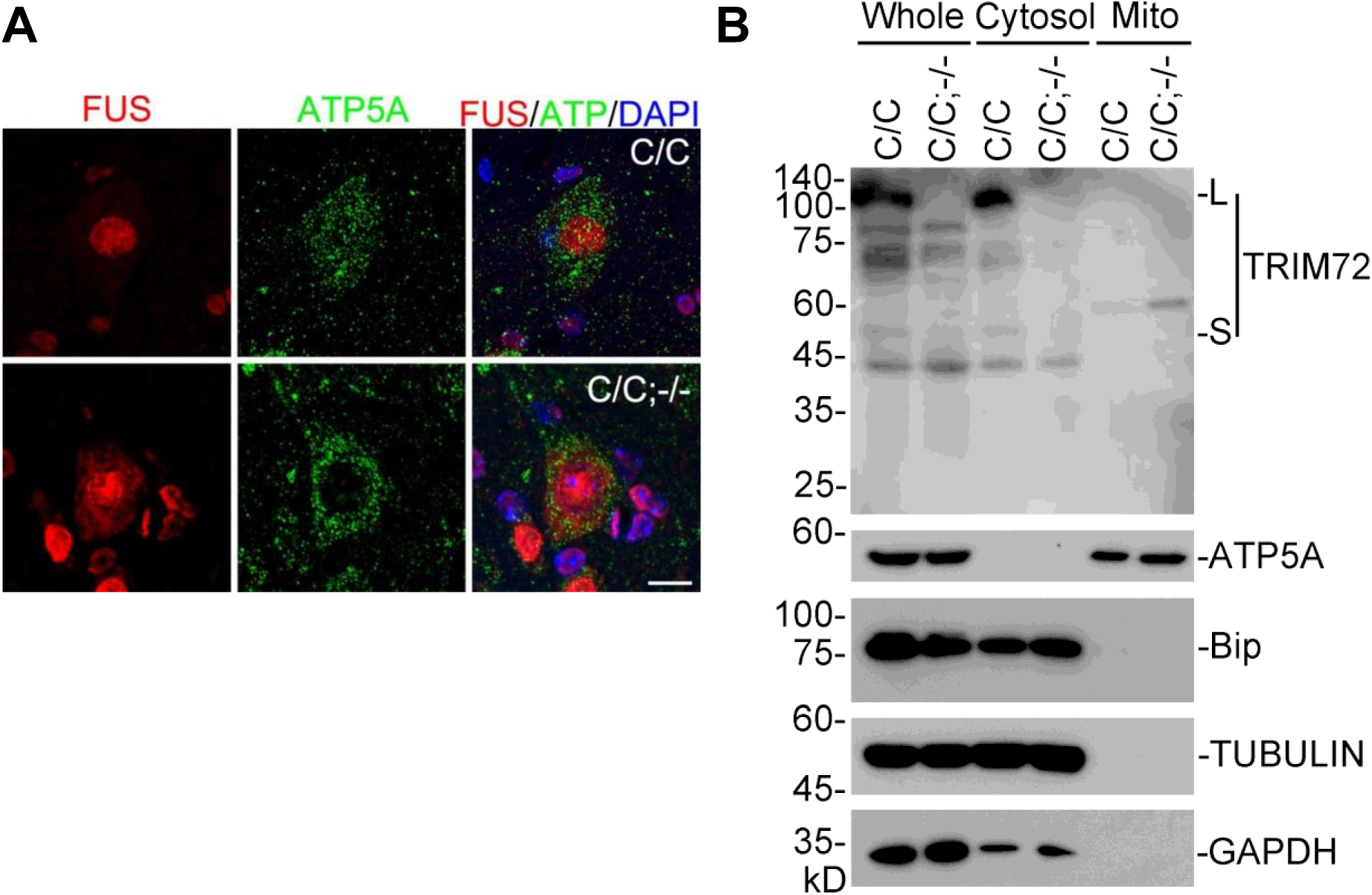
Mutant FUS and TRIM72 are not associated with mitochondria. (A) Sections of lumbar 4-5 spinal cords in C/C and C/C;−/− mice were stained with FUS and ATP5A antibodies. Age, 1.5 month of age. Scale bar, 10 μm. (B) Mitochondria were separated from mice cortices by gradient centrifugation. Bip served as cytosol marker, ATP5A served as mitochondria marker.

**Figure S12.**
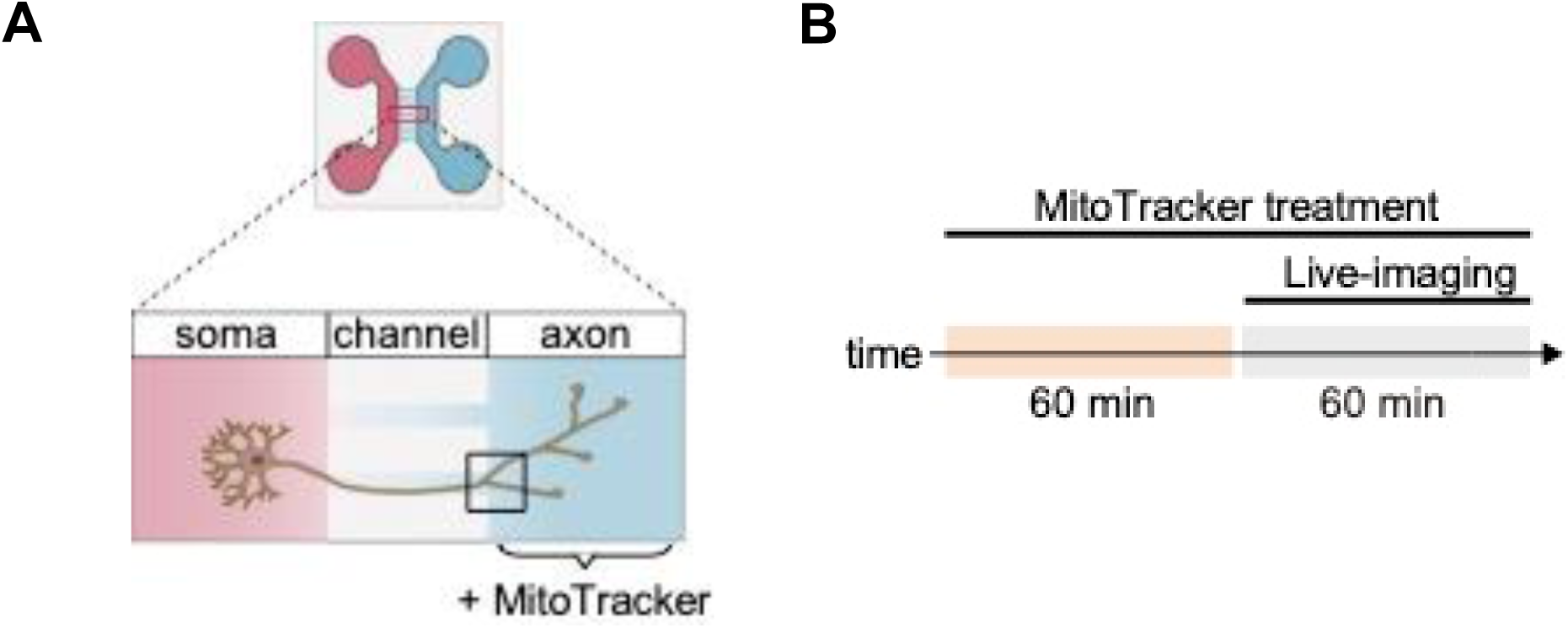
Experimental design for imaging mitochondria. (A) Schematic representation of the microfluidic device designed to separate axons from cell bodies. Device comprised of two chambers adhered to a glass-bottom dish, featuring a soma (red) and an axon chamber (blue). The black square box depicts the live-imaging field. (B) Timeline for mitochondrial labeling and live imaging. On 8DIVs, MitoTracker was added to the axon chamber. After 1 hour of labeling, live-imaging of the axon chamber was conducted to capture axonal mitochondrial dynamics over a 15-minute period.

**Figure S13.**
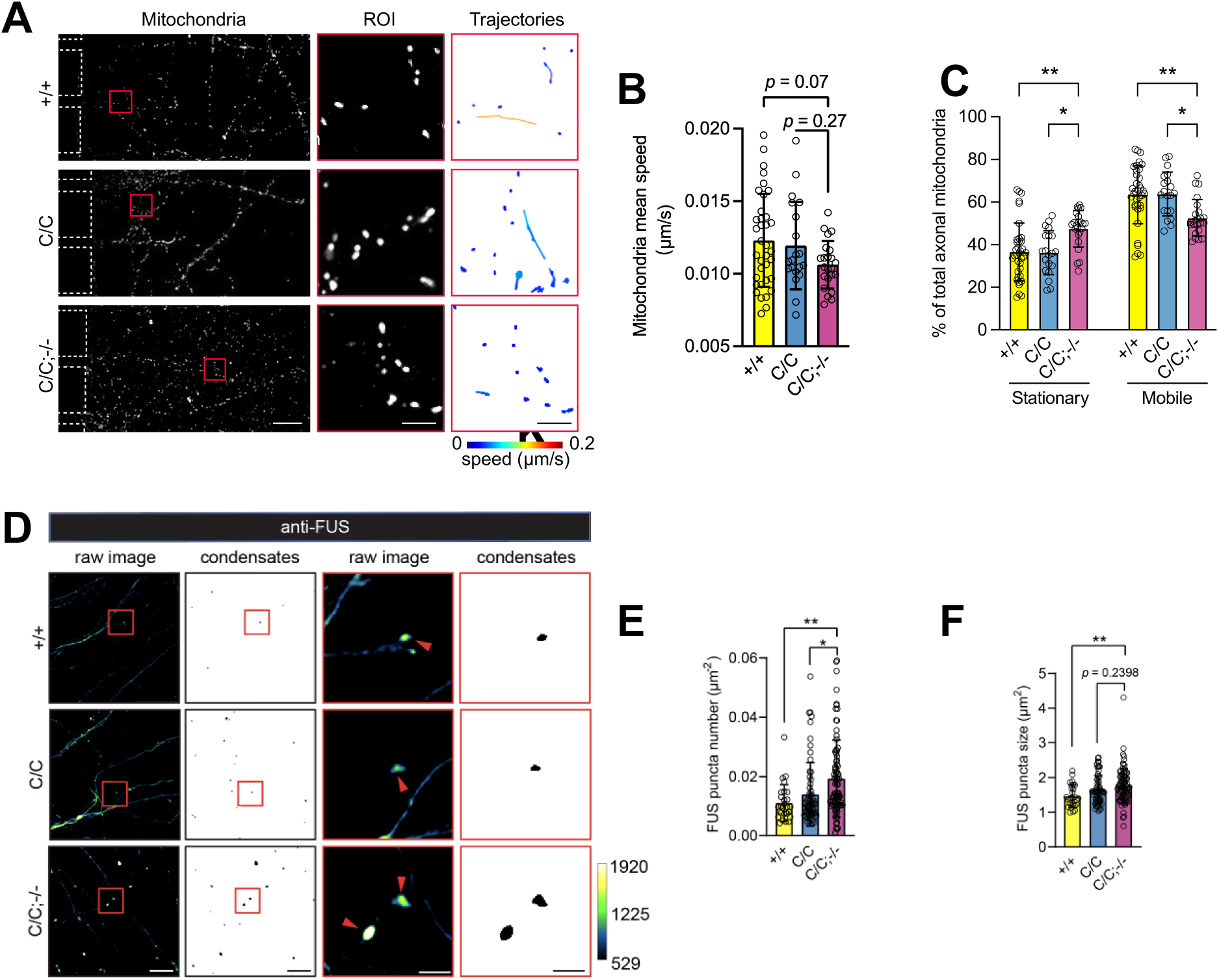
Loss-of-function of *Trim72* worsens axonal mutant FUS condensates, which in turn impairs axonal mitochondria dynamics. (**A**-**C**) Representative live-imaging graphs of axonal mitochondria with the indicated genotypes (**A**). Boxed regions are amplified in the middle panels, with automatically detected mitochondrial trajectories shown in the right panels. Scale bar, 20 (left); 5 μm (middle and right). Quantification of the mean speed of mitochondrial trafficking (**B**) and the ratio of stationary (mean speed < 0.005 μm/s) and mobile (mean speed ≥ 0.005 μm/s) mitochondria with indicated genotypes (**C**). (**D**-**F**) Representative confocal images of endogenous FUS levels in the axons (8DIVs) in microfluidic devices (**D**). Black-white masks depict automatically extracted FUS aggregates (circularity > 0.8 and area > 0.5 μm_2_). FUS aggregates indicated by arrowheads. Scale bar, 20 (left); 5 μm (right). Quantification of the numbers (**E**) and average sizes (**F**) of axonal FUS aggregates. In **B**, **C**, **E**, and **F**, values were presented as mean ± SD of three independent preparations; in **B** and **C**, +/+, n = 36; C/C, n = 20; C/C;−/−, n = 23; in **E** and **F**, +/+, n = 28; C/C, n = 63; C/C;−/−, n = 85. N.S., no statistical significance; \**p* < 0.05; \*\**p* < 0.01; \*\*\**p* < 0.001; ANOVA, SPSS.

**Figure S14.**
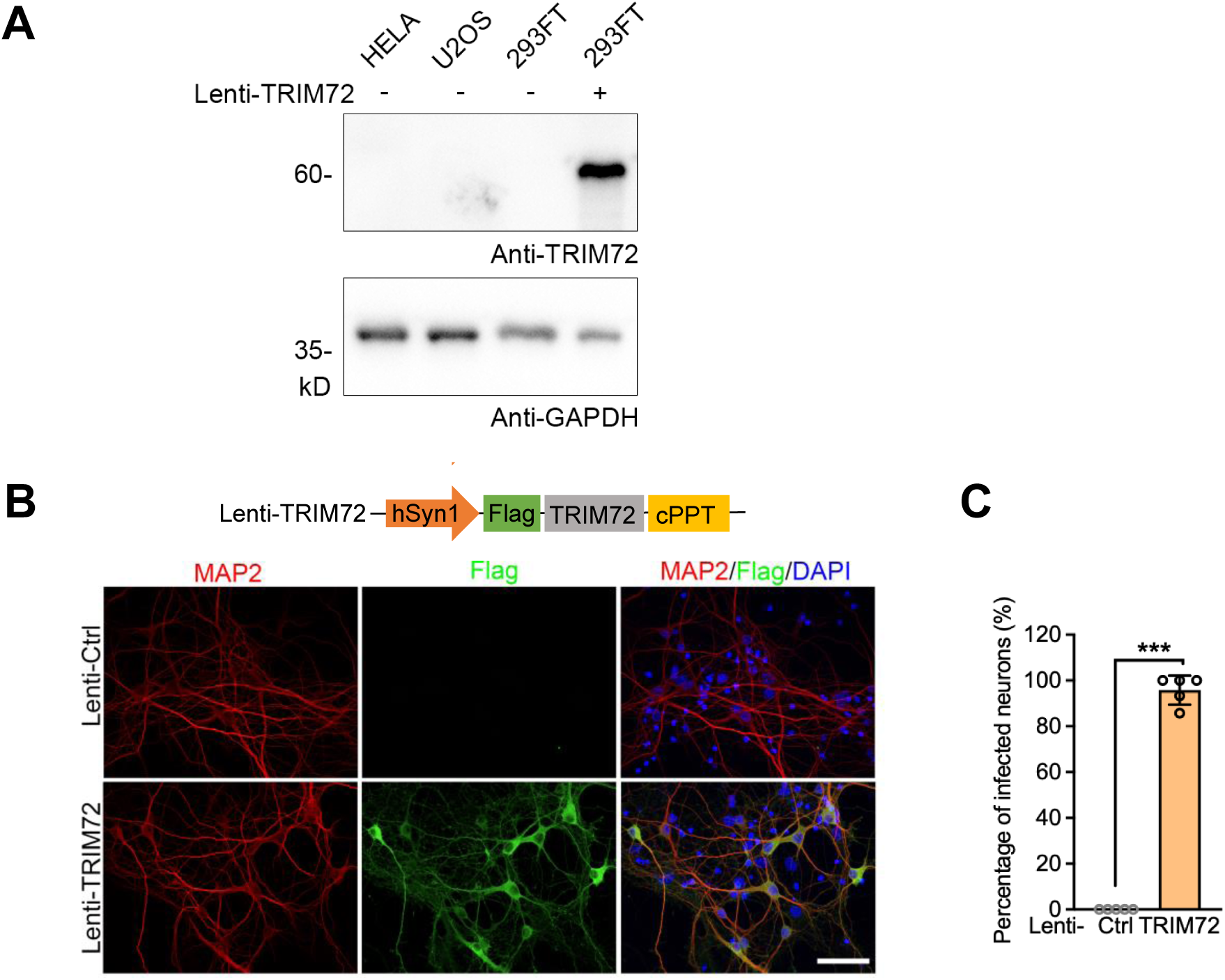
TRIM72 expression in cultured cell lines and in neurons by lentiviral particles. (**A**) Undetectable TRIM72 expression in HELA, U2OS, and 293FT cell lines. Stable expression of TRIM72 by lentivirus was employed as positive control. GAPDH served as loading control. (**B** and **C**) MAP2 was used as a neuronal marker. Flag-tagged TRIM72 was stained by Flag antibody. Scale bar, 50μm. Lenti-Ctrl, empty lentiviral particles. Data summary of infection rate of Lenti-TRIM72 was shown in (**C**). Values were presented as mean ± SD (n =5), T-test.

**Figure S15.**
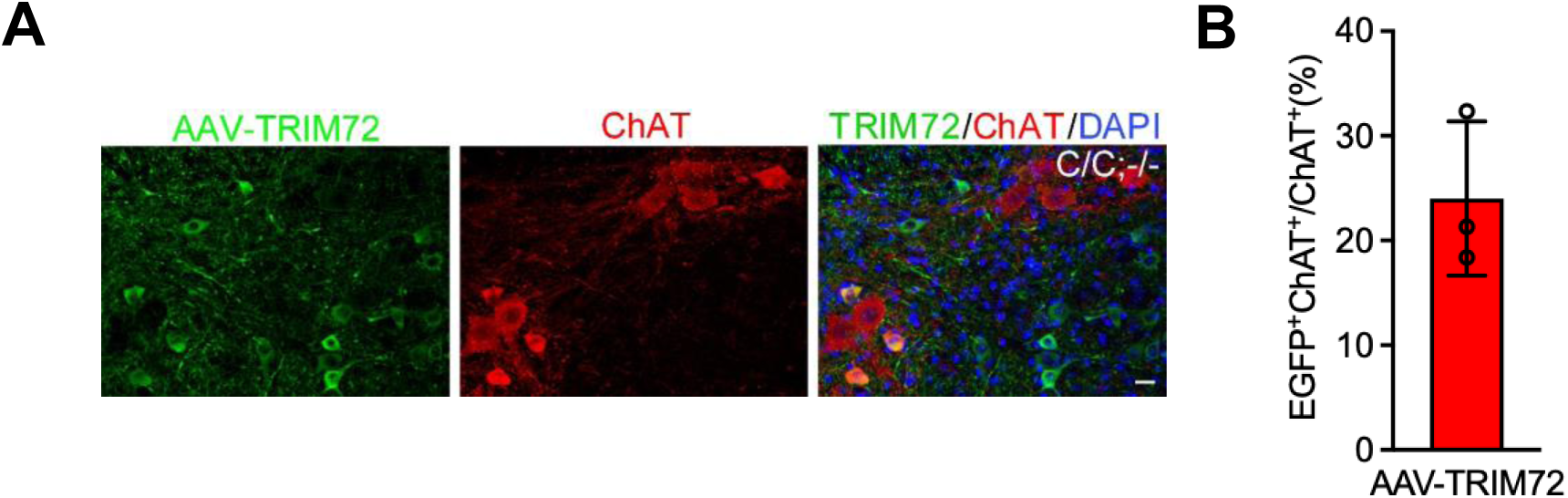
TRIM72 expression in spinal cord motor neurons by AAV-TRIM72. (**A**) ChAT was used as a motor neuron marker. AAV-TRIM72 (AAV PHP.eB-TRIM72) was stained by EGFP antibody. Scale bar, 50μm. TRIM72 expression in motor neurons was quantified by the ratio of EGFP-positive/ChAT-positive neurons to ChAT-positive neurons. Data summary was shown in (**B**). Value was presented as mean ± SD (n = 3).

**Figure S16.**
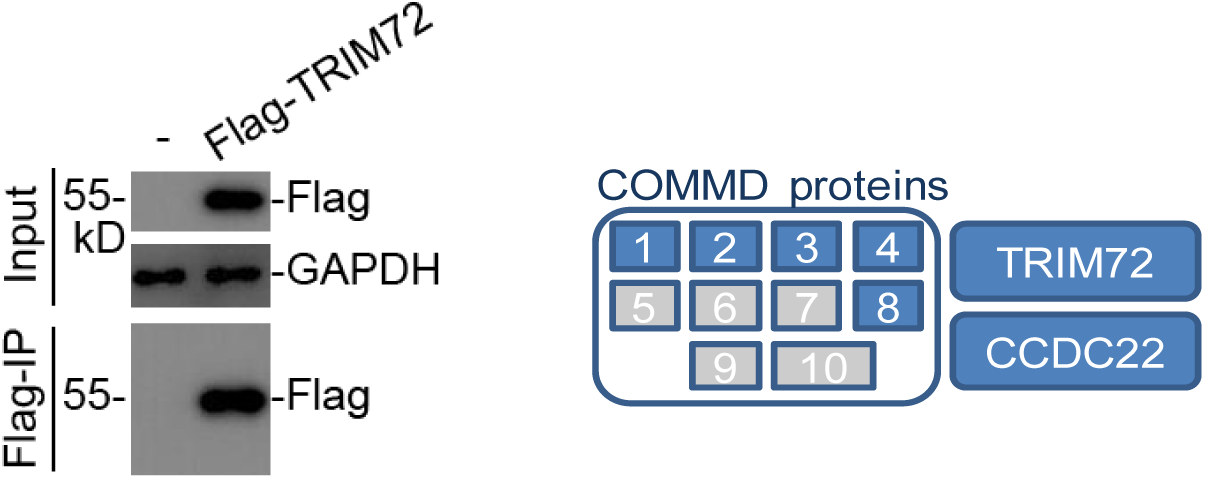
Identification of TRIM72-interacting partners by Flag-IP-MS. Hela cells were stably expressed with Flag-TRIM72 by lentivirus and applied for immunoprecipitation by Anti-Flag M2-beads. CCDC22 and COMMD proteins were identified as interacting partners of TRIM72 by Flag-IP-MS.

**Figure S17.**
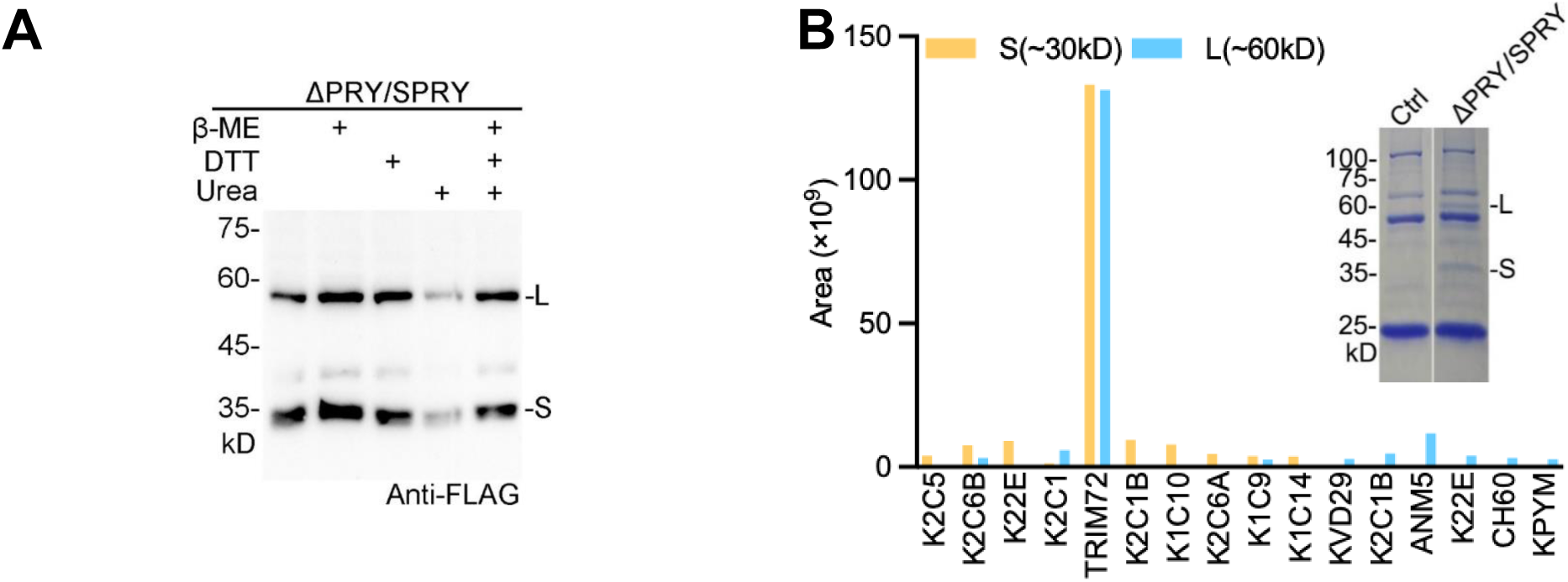
Small and large molecular weight of ΔPRY/SPRY. (**A**) Large molecular weight (L) of ΔPRY/SPRY was insensitive to DTT, urea, and β-ME. DTT: 10 mM; urea: 8 M; β-ME: 200 mM. (**B**) 293FT cells stably expressing ΔPRY/SPRY were precipitated by M2-beads. Bar plot presented protein identities at small (S) and large molecular weight (L) by Flag-IP-MS. Inserted, Coomassie blue-stained bands after Flag-IP. Note that the S and L bands appeared only in ΔPRY/SPRY but not in Ctrl (empty vector).

**Figure S18.**
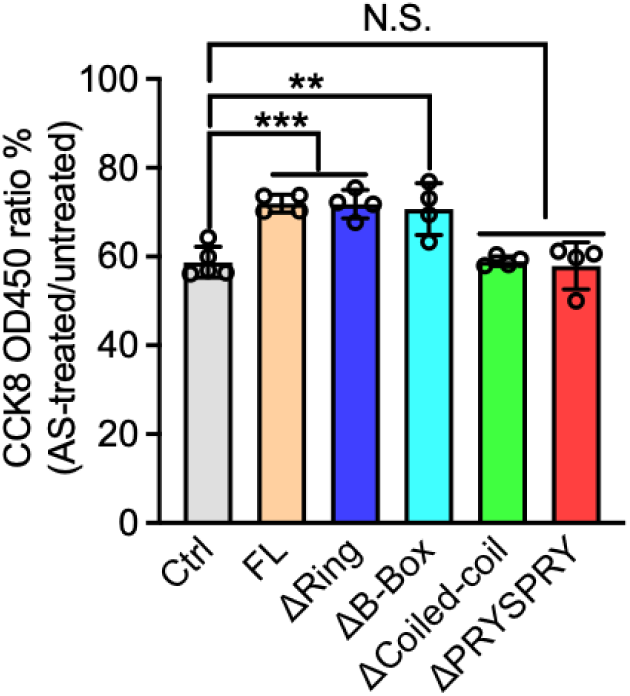
Coiled-coil and PRY/SPRY, but not Ring and B-box domains, are required for TRIM72-mediated antioxidation. Hela cells were stably expressed with and without TRIM72 full-length (FL) and truncated forms by lentivirus and subsequently treated with AS for 2 hours. After AS treatment, cell viability/dehydrogenase activity was measured by CCK-8 assay within 2 hours. Ctrl, empty vector. Experimental procedure was illustrated I (Figure 4A). n = 4-5.

**Figure S19.**
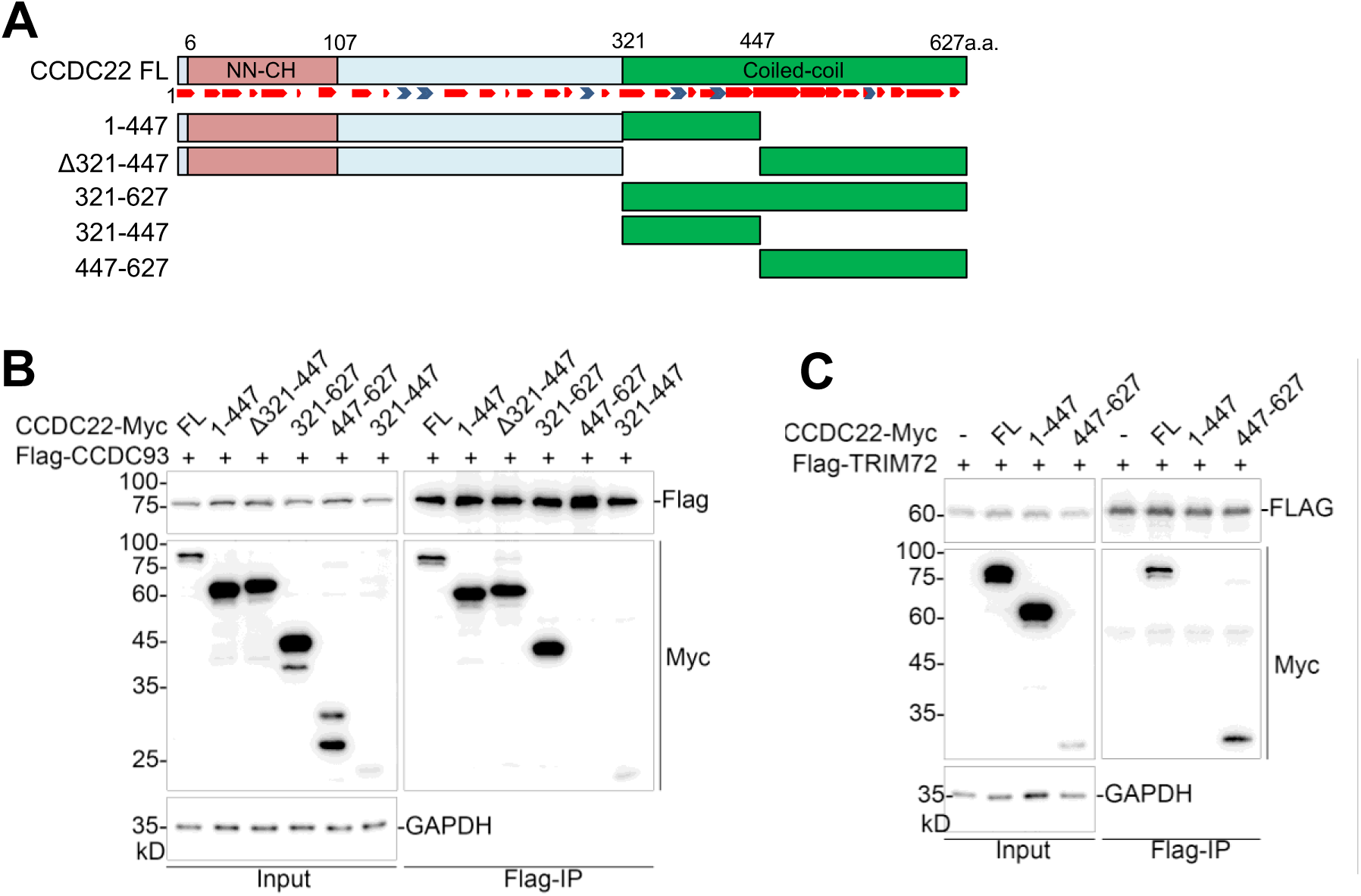
CCDC22 interact with CCDC93 and TRIM72 with its different domains. (**A**) Protein domain and secondary structure features of CCDC22, predicted by AlphaFold-2. (**B** and **C**) Expressions of truncated CCDC22 with Flag-tagged CCDC93 or Flag-tagged TRIM72, respectively, in cells applied for immunoprecipitation by anti-Flag M2-beads. CCDC22 interact with CCDC93 and TRIM72 with different domains.

**Figure S20.**
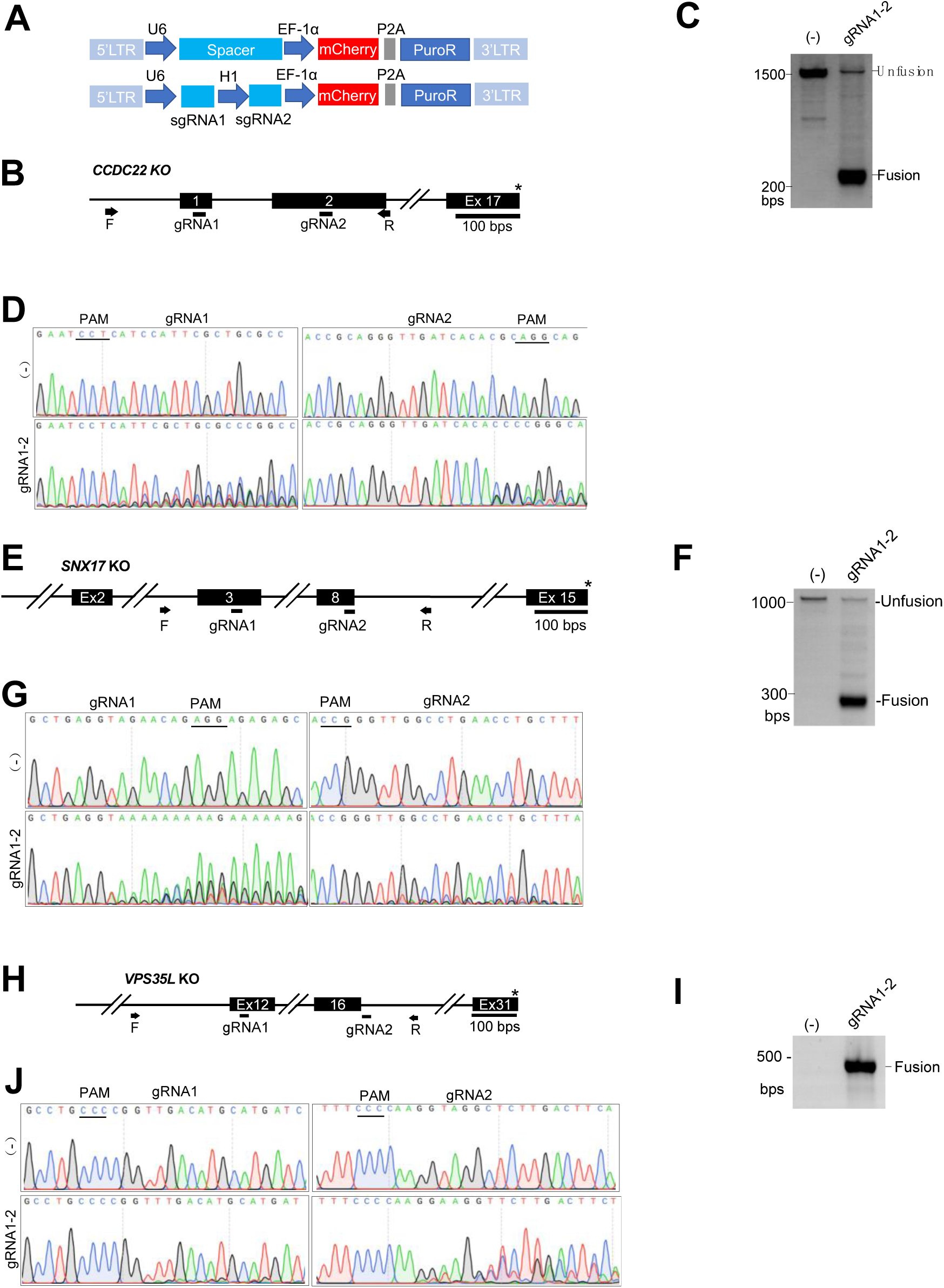
Generations of *CCDC22*, *SNX17*, and *VPS35L* KO single cell clones. (**A**) Lentiviral dual-gRNA systems for desired gene KO (PMID: 35383205). Dual gRNAs were driven by U6 and H1 promoters, respectively. Spacer sequences served as control. (**B**-**E**) Dual-gRNA design for *CCDC22* KO (**B**). The removal of DNA fragments between the dual gRNA sites in *CCDC22* KO single cell clone was detected by gDNA PCR (**C**), which was confirmed by Sanger sequencing (**D**) and western blot (**E**). (**F**-**I**) Dual-gRNA design for *SNX17* KO (**F**). The removal of DNA fragments between the dual gRNA sites sites in *SNX17* KO single cell clone was detected by gDNA PCR (**G**), which was confirmed by Sanger sequencing (**H**) and western blot (**I**). (**J**-**L**) Dual-gRNA design for *VPS35L* KO (**J**). The removal of DNA fragments between the dual gRNA sites sites in *VPS35L* KO single cell clone was detected by gDNA PCR (**K**), which was confirmed by Sanger sequencing (**L**). In **B**, **F**, and **J**, primers flanking the gRNA editing sites were labeled with black arrows for genotyping. In **E** and **I**, numbers represent individual KO single cell clones.

**Figure S21.**
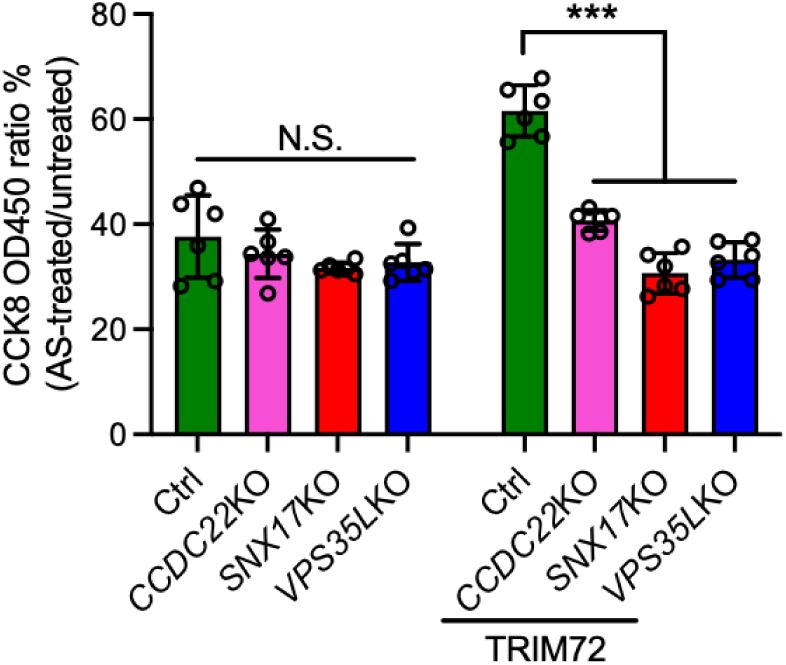
CCC and Retriever complex are involved in antioxidation effect mediated by TRIM72. *CCDC22*, *VPS35L*, and *SNX17* KO disrupted the antioxidation properties of TRIM72 in 293FT cells. Ctrl, cells expressing Cas9. n = 6

**Figure S22.**
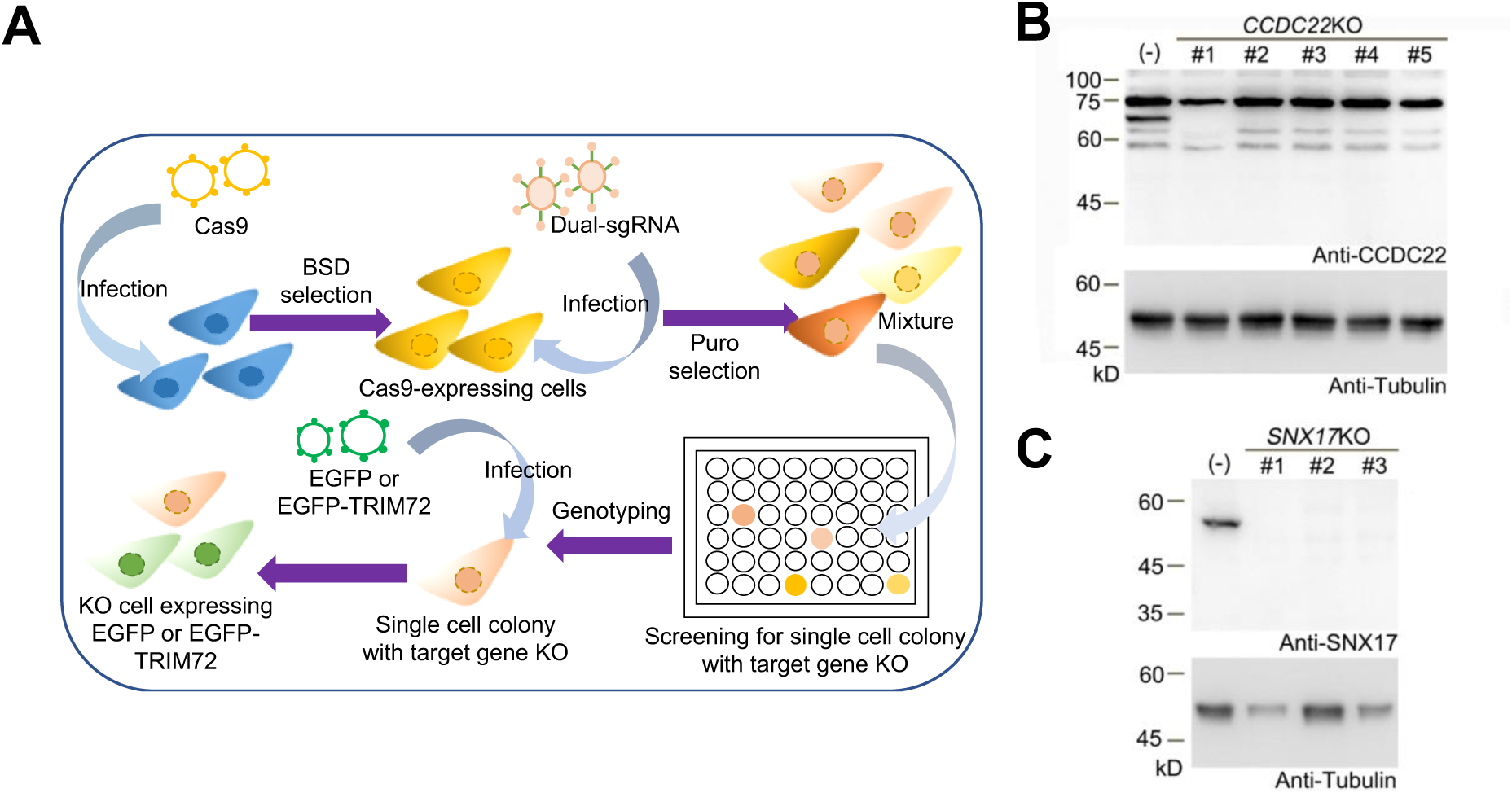
Workflow of construction of single cell clones with gene KO and expression of EGFP or EGFP-TRIM72. (**A**) 293FT or U2OS cells were infected with Cas9 lentivirus. After blasticidin (BSD) selection, Cas9 expressing cells were infected with dual-sgRNA lentivirus for target gene KO. After puromycin (Puro) selection, *CCDC22*, *SNX17*, and *VPS35L* KO cells were confirmed by gDNA PCR as shown in **Figure S20**. KO cells were cultured in 48-well plate for single clone selection. Resulting cells were infected by EGFP or EGFP-TRIM72 expressing lentivirus. EGFP-positive cells from these single cell clones were visualized under fluorescent microscope for membrane damage by laser light and exosome purification. (**B** and **C**) *CCDC22 and SNX17* KO single cell clones were confirmed by western blot. Numbers represent individual KO single cell clones.

**Figure S23.**
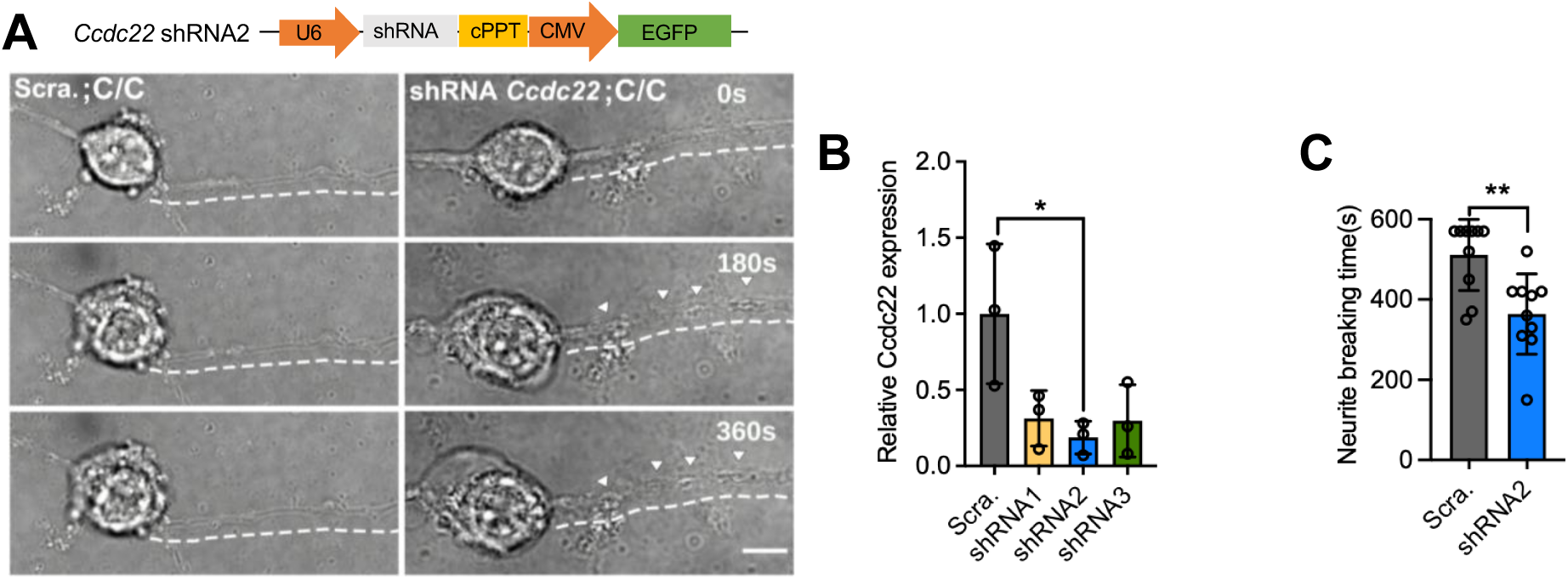
*Ccdc22* is required for TRIM72-mediated neurite protection. (**A**) Cultured cortical neurons (C/C genotype) infected with lentiviral scrambled (Scra.) shRNA (negative control) or shRNA2 against *Ccdc22*. GFP-positive infected neurons were photodamaged and monitored by time-lapse imaging. Lentiviral shRNA vector contains U6-driven shRNA and CMV-driven EGFP. Scale bar, 10 µm (**B** and **C**) *Ccdc22* shRNA knockdown efficiencies were summarized (**B**). Data summary of neurite breaking time in neurons (C/C genotype) infected with scrambled (Scra.) or shRNA2 (**C**). in **B**, n = 3, in **C**, n = 10.

**Figure S24.**
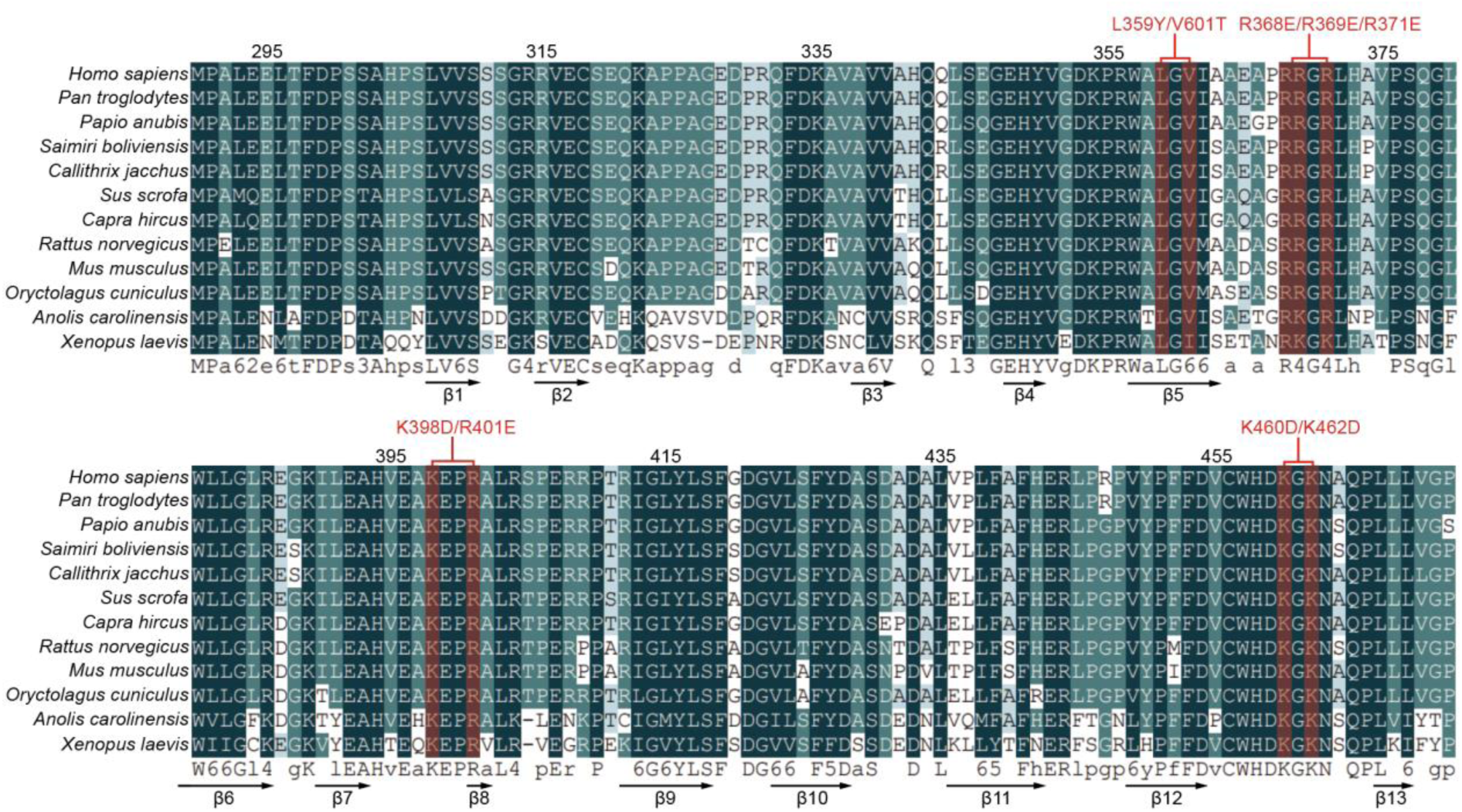
Conservation analysis of TRIM72 PRY/SPRY domain. Mutations of several potential key residues crucial for TRIM72 release were labeled in red. The β-sheets in the domain were labeled, as previously reported (PMID: 37770719).

**Figure S25.**
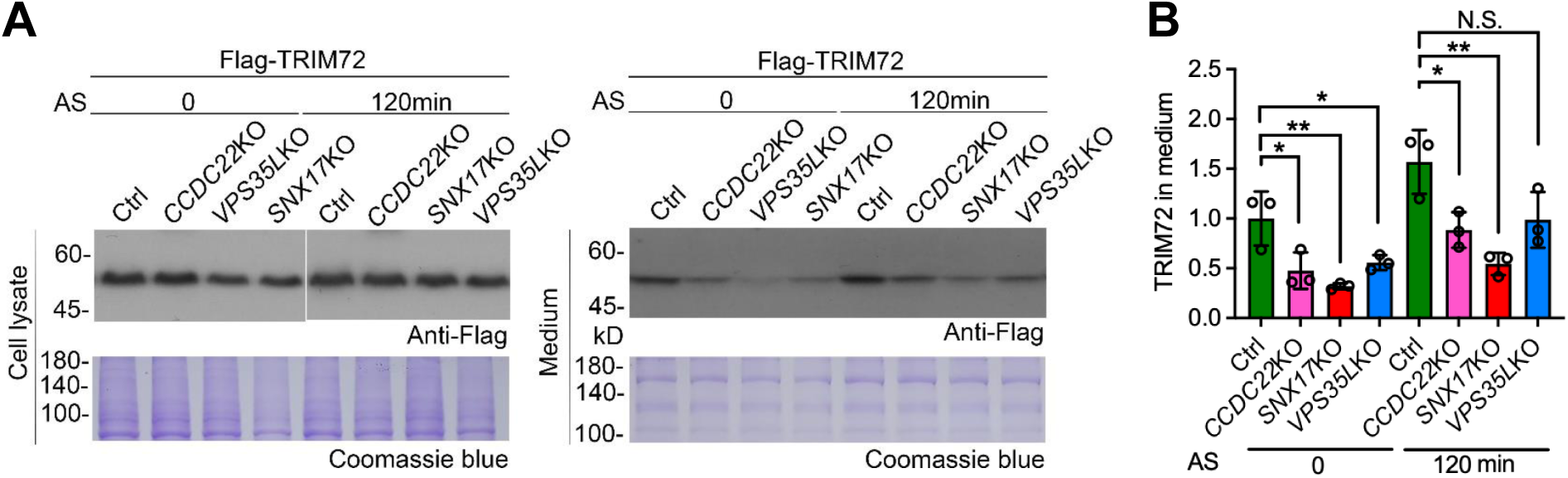
***CCDC22*, *VPS35L*, and *SNX17* are required for TRIM72 release.** (A) Release of FLAG-TRIM72 in culture medium was measured in *CCDC22*, *SNX17* or *VPS35L* KO cells with or without AS treatment (0.5 mM, 2 hours). (B) Relative amount of FLAG-TRIM72 in the media was summarized.

**Figure S26.**
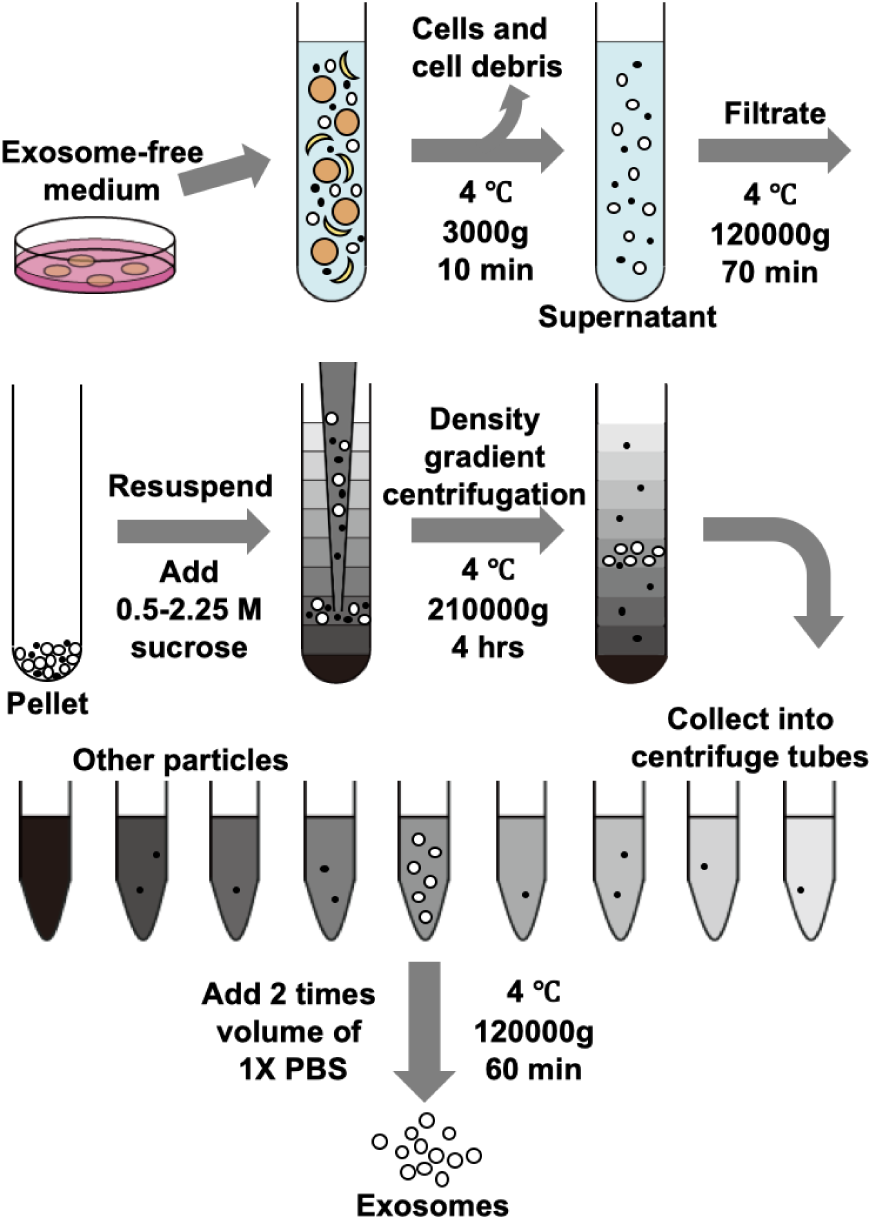
Workflow of exosome purification. 293FT cell culture media were collected for exosome purification by sucrose gradient centrifugation as previously reported (PMID: 8642258). Exosome-free FBS were used for cell culture and culture medium was collected and centrifuged at 3000g for 10 minutes to remove cells and cell debris. Supernatant was collected and passed through a 0.22μm sterile filter and the filtered supernatant was centrifugated at 120,000g for 70 minutes to collect exosomes. The pellet was resuspended and centrifuged by sucrose density gradient centrifugation at 210,000g for 4 hours to purify exosomes. Each fraction was collected in centrifugate tubes. Two-fold volume of PBS were added into exosomes fraction and centrifugate at 120,000g for 60 minutes to collect exosomes for further analysis.

**Figure S27.**
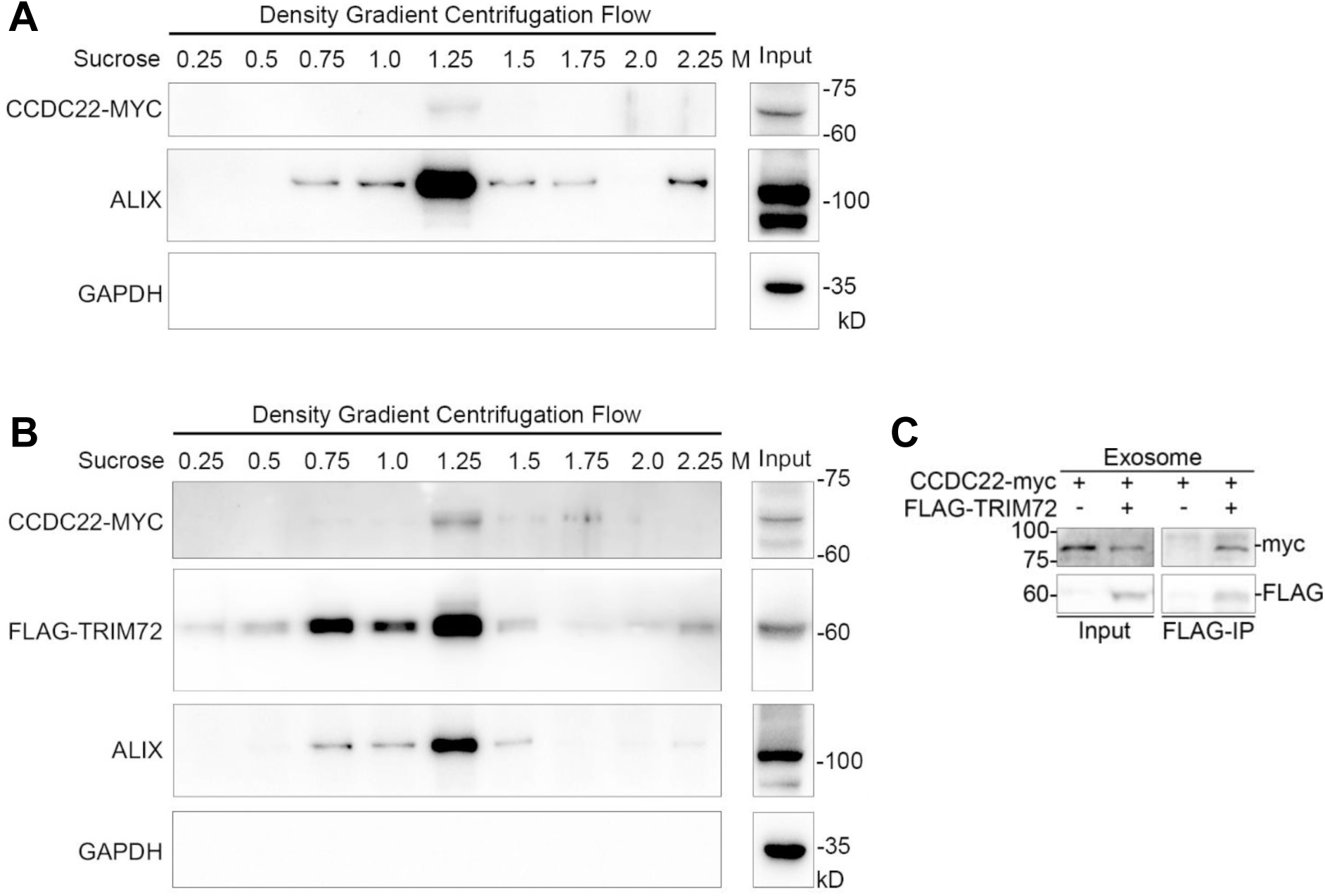
Presence of CCDC22 in exosome and exosomal interaction between CCDC22 and TRIM72. (A) The culture media of 293FT cells expressing CCDC22-MYC was applied for sucrose density gradient centrifugation. ALIX, an exosome marker. (**B** and **C**) CCDC22 was present in the exosomal fraction of cells stably expressing CCDC22-MYC and FLAG-TRIM72 (**B**). Exosomal interaction between CCDC22-Myc and FLAG-TRIM72 was demonstrated by FLAG-IP.

**Figure S28.**
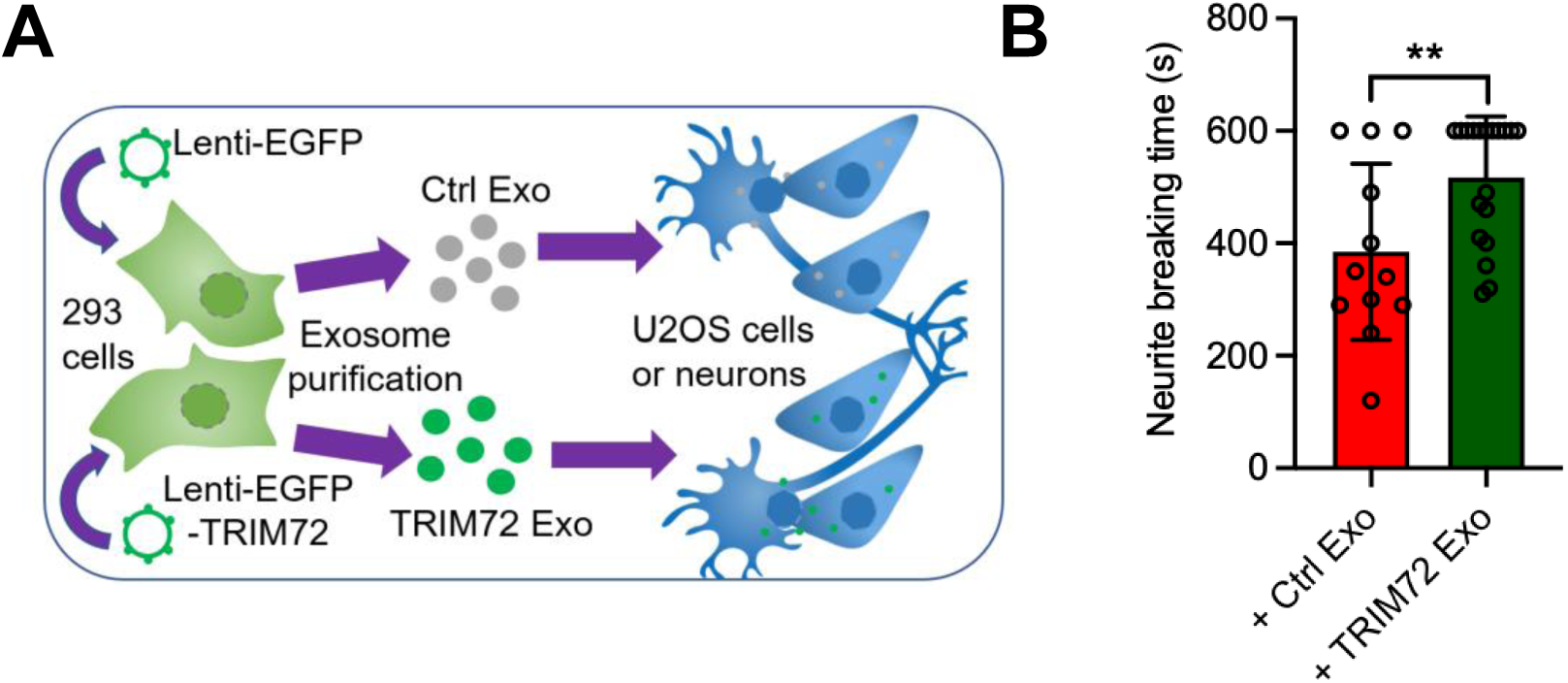
Experimental procedure for extracellular application of TRIM72-containing exosome. (A) Illustration of the experimental procedure for extracellular application of TRIM72-containing exosome. The exosomes were isolated from the culture medium of 293FT cells expressing EGFP (Ctrl Exo) or EGFP-TRIM72 (TRIM72 Exo), and then extracellularly applied to U2OS cells or cultured neurons. (B) Neurite breaking time of cultured cortical neurons (C/C;−/−) treated with Ctrl Exo or TRIM72 Exo was summarized. Neuron soma photodamage and time-lapse imaging conditions as shown in Figure 5.

**Figure S29.**
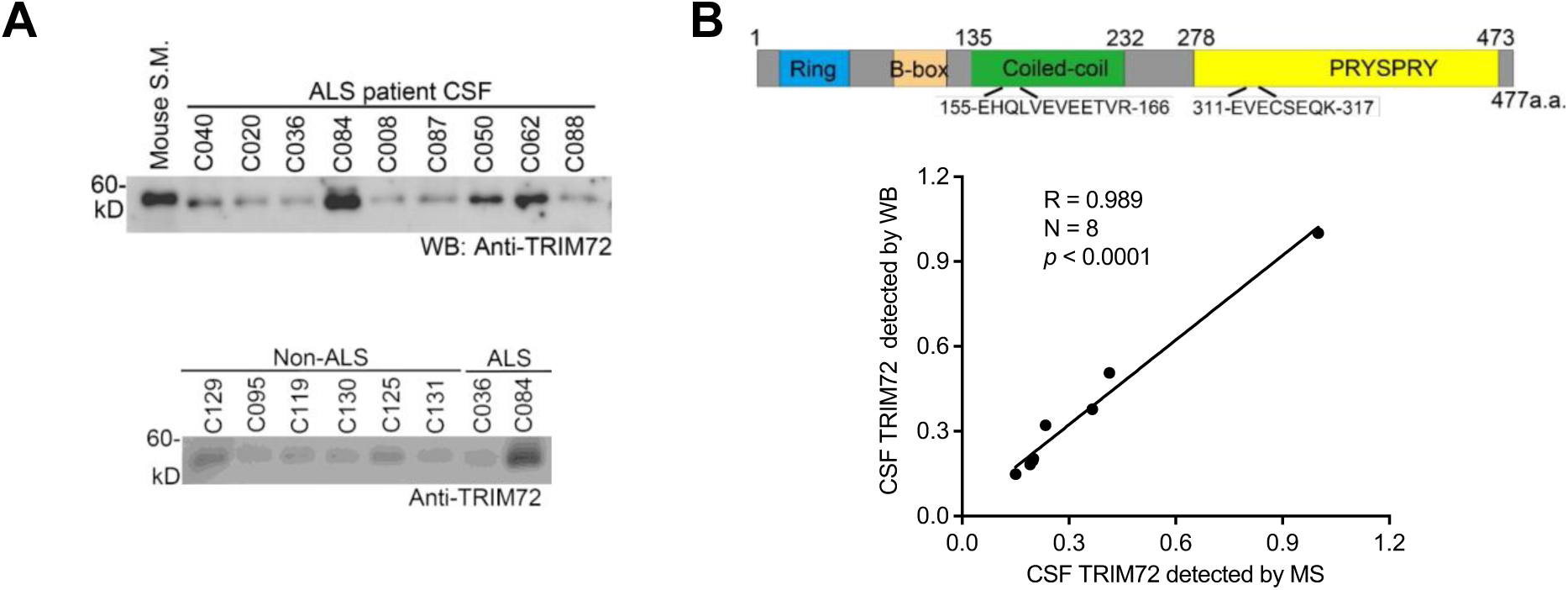
Identity of TRIM72 detected by western blot in CSF was confirmed by MS. (A) Representative images of CSF TRIM72 level in ALS and non-ALS patients. (B) Locations of 2 polypeptide sequences for TRIM72 MS detection. Correlation (r = 0.989, n = 8) between CSF TRIM72 levels detected by MS and by western blot.

**Figure S30.**
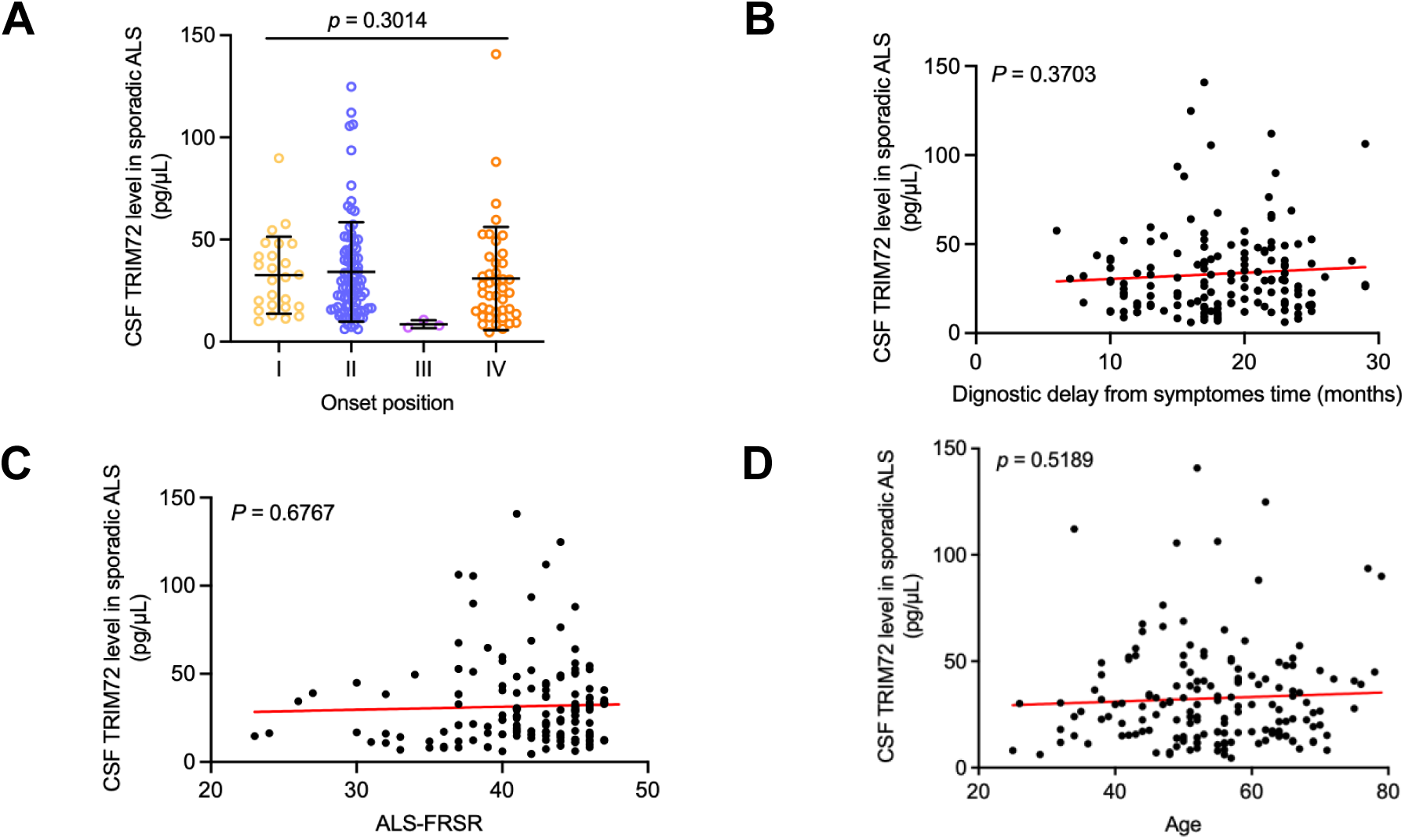
CSF TRIM72 level was not correlated with ALS onset position, diagnostic delay from symptoms, FRS score, and patient age. CSF TRIM72 level was not significantly correlated with ALS onset position (**A**, *p* = 0.3014), diagnostic delay from symptoms (**B**, r = 0.073, *p* = 0.3703), and ALS-FRSR (**C**, r = 0.11, *p* = 0.6767), and patient ages when CSF samples were collected (**D**, r = 0.14, *p* = 0.5189).

**Figure S31.**
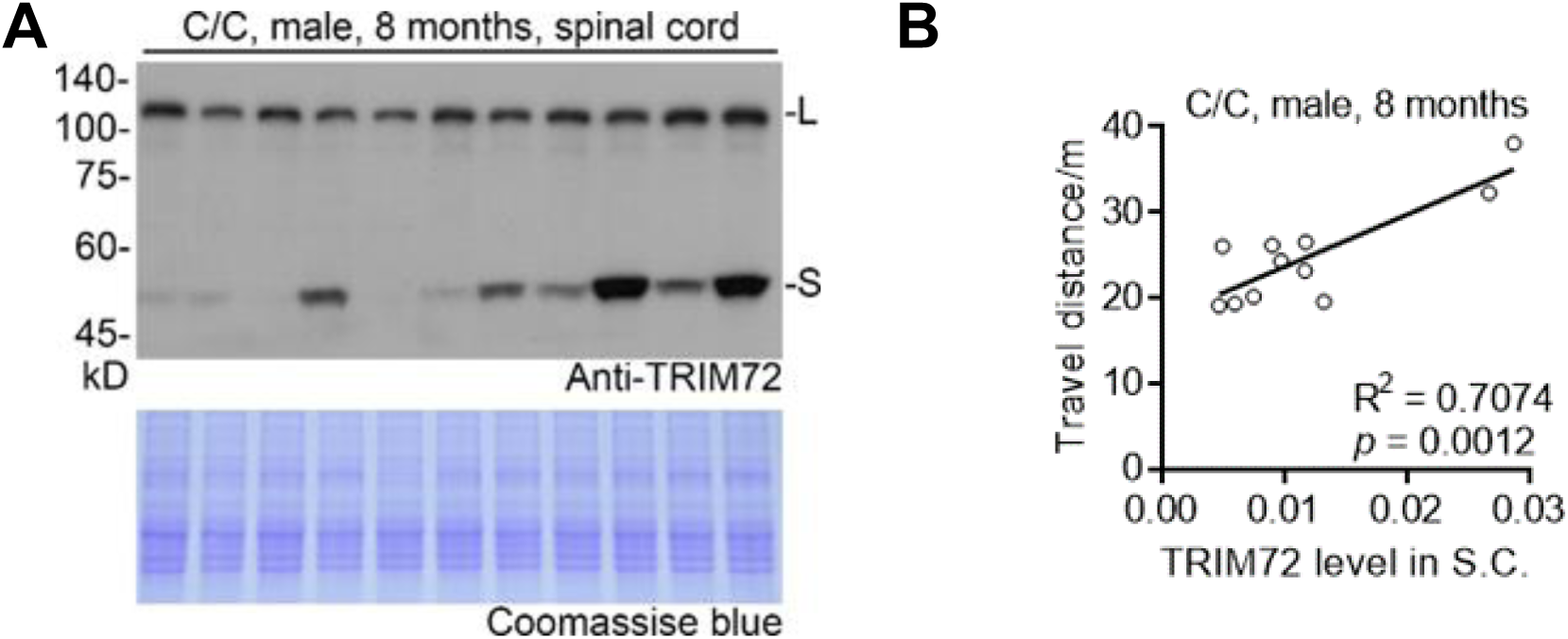
TRIM72 expression level in the FUS-R521C mutant spinal cord predicts motor function. (A) TRIM72 expression in the FUS-R521C (C/C) mutant spinal cord. Coomassie blue served as loading control. Age, 8 months (n = 11). Spinal cords were collected for western blot after these mice were applied for Open Field test. (B) TRIM72 level in the spinal cord (S.C.) was correlated with the mutant mouse travel distance in Open Field test (10 minutes, n = 11).

**Figure S32.**
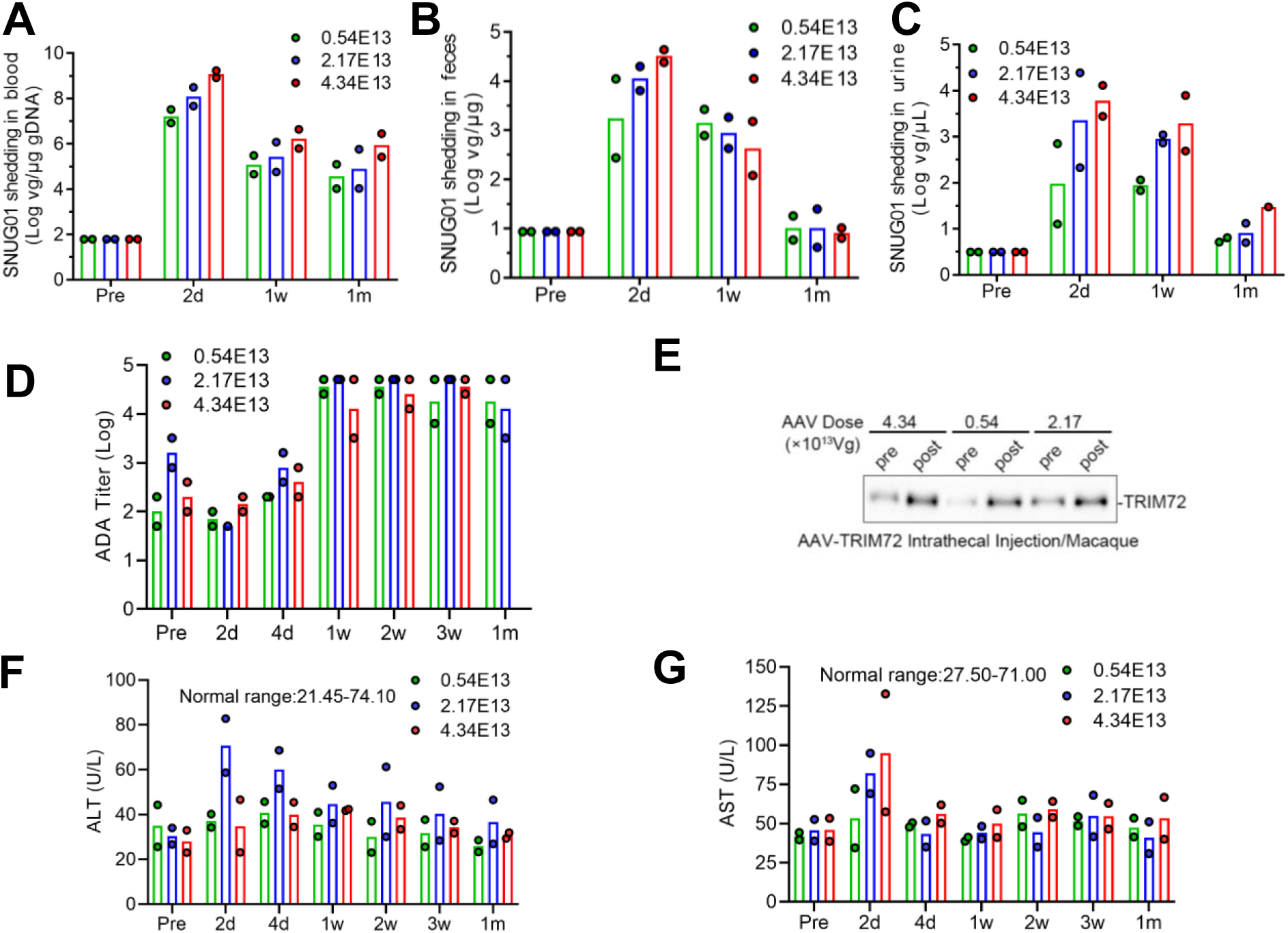
SNUG01 pharmacokinetics in macaques. (**A**-**C**) Viral shedding of SNUG01 DNA in blood (**A**), faces (**B**), and urine (**C**). Three doses of SNUG01 (0.54, 2.17, and 4.34E13 vg/macaque, intrathecal injection). Each dose group included 2 macaques (one male and one female). Samples were collected as indicated. (D) ADA (anti-drug antibody) was detected before and after SNUG01 injection. (E) CSF TRIM72 levels were measured by western blot before (pre) and after (post) SNUG01 administrations. (**F** and **G**) Normal ALT (alanine aminotransferase) and AST (aspartate transaminase) levels in peripheral blood in macaques intrathecally injected with SNUG01.

**Figure S33.**
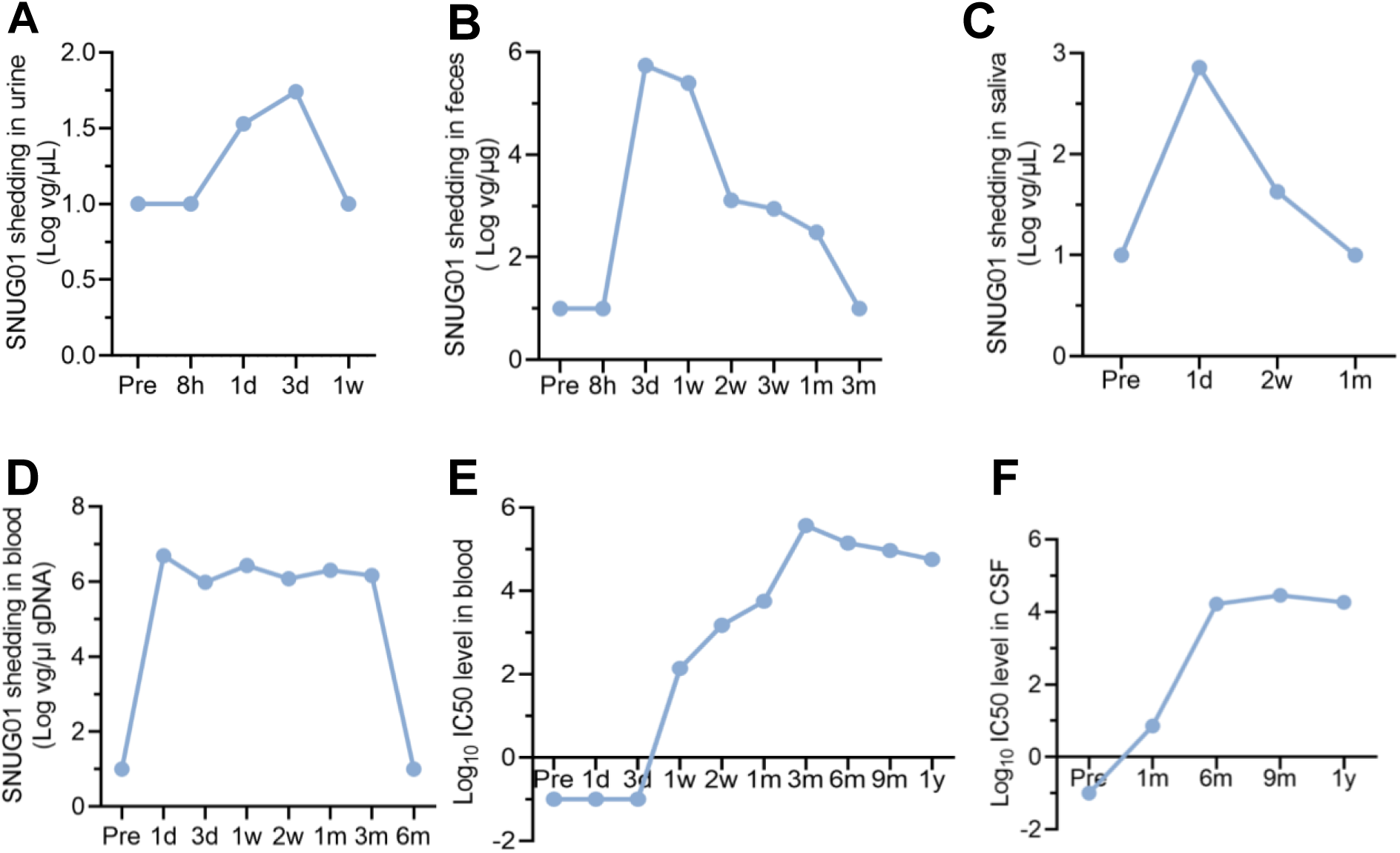
SNUG01 shedding and AAV9 neutralizing antibody in an ALS patient. (**A**-**D**) Viral shedding of SNUG01 in the urine (**A**), faces (**B**), blood (**C**), and saliva (**D**). SNUG01, 1.2E14vg, intrathecal injection. Samples were collected as indicated. (**E** and **F**) AAV9 neutralizing antibody level in the blood (**E**) and CSF (**F**). In **A**-**F**, d, day; w, week; m, month; y, year.

**Figure S34.**
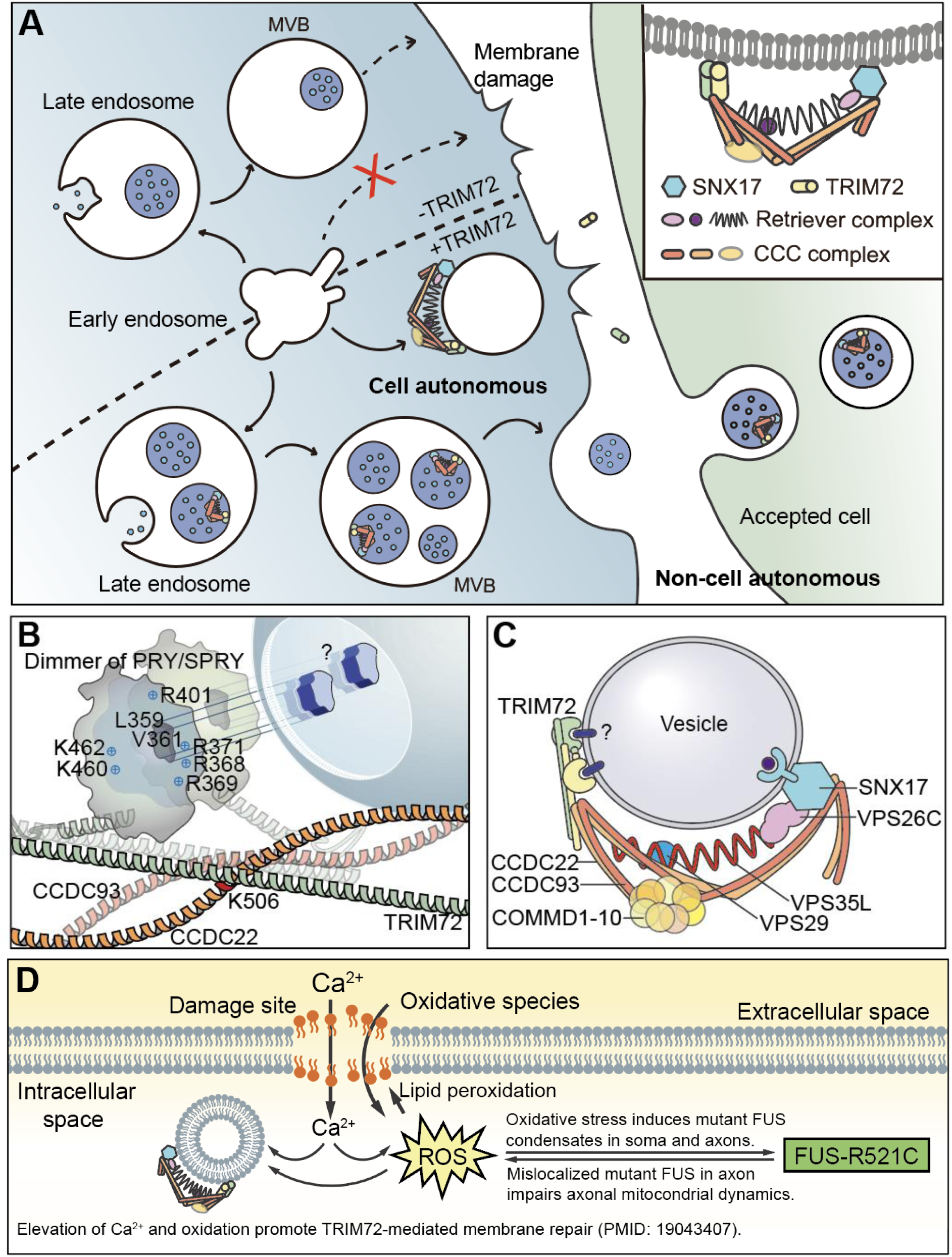
Working model for TRIM72-mediated protection. (A) A model illustrating how TRIM72 mediates protection during membrane repair through both cell-autonomous and non-cell-autonomous mechanisms. In response to cell membrane damage, TRIM72 works with the Commander complex to facilitate membrane repair. Exosomal TRIM72 may protect accepting cell from membrane damage. (B) Point mutations in many positively charged residues on the edge of the PRYSPRY domain were crucial for TRIM72 release, as well as L359 and V361. We hypothesize that L359 and V361 located in a binding pocket of TRIM72, which interact with unknown membrane protein (?) to facilitate the TRIM72 membrane recognition and tethering. Q507 and K506 (illustrated only) are key residues crucial for CCDC22 interaction. (C) Illustration of Commander/TRIM72 complex. Besides SNX17 anchoring site, dimer of TRIM72 provide additional two sites for vesicle recognition. (D) When cell faces membrane damage, the extracellular Ca_2+_ and oxidative species flush into neuron, because of high extracellular concentrations of these two. On one hand, elevation of Ca_2+_ and oxidation promotes TRIM72-mediated membrane repair (PMID: 19043407). On the other hand, the ROS induces mutant FUS condensate in neuron soma and axon. Especially, axonal mutant FUS condensates impair axonal mitochondrion dynamics, which may in turn generate more ROS. As one of the major sources for membrane damage, the intracellular ROS generated by mutant FUS will further damage membrane via lipid peroxidation.

**Movie S1.**
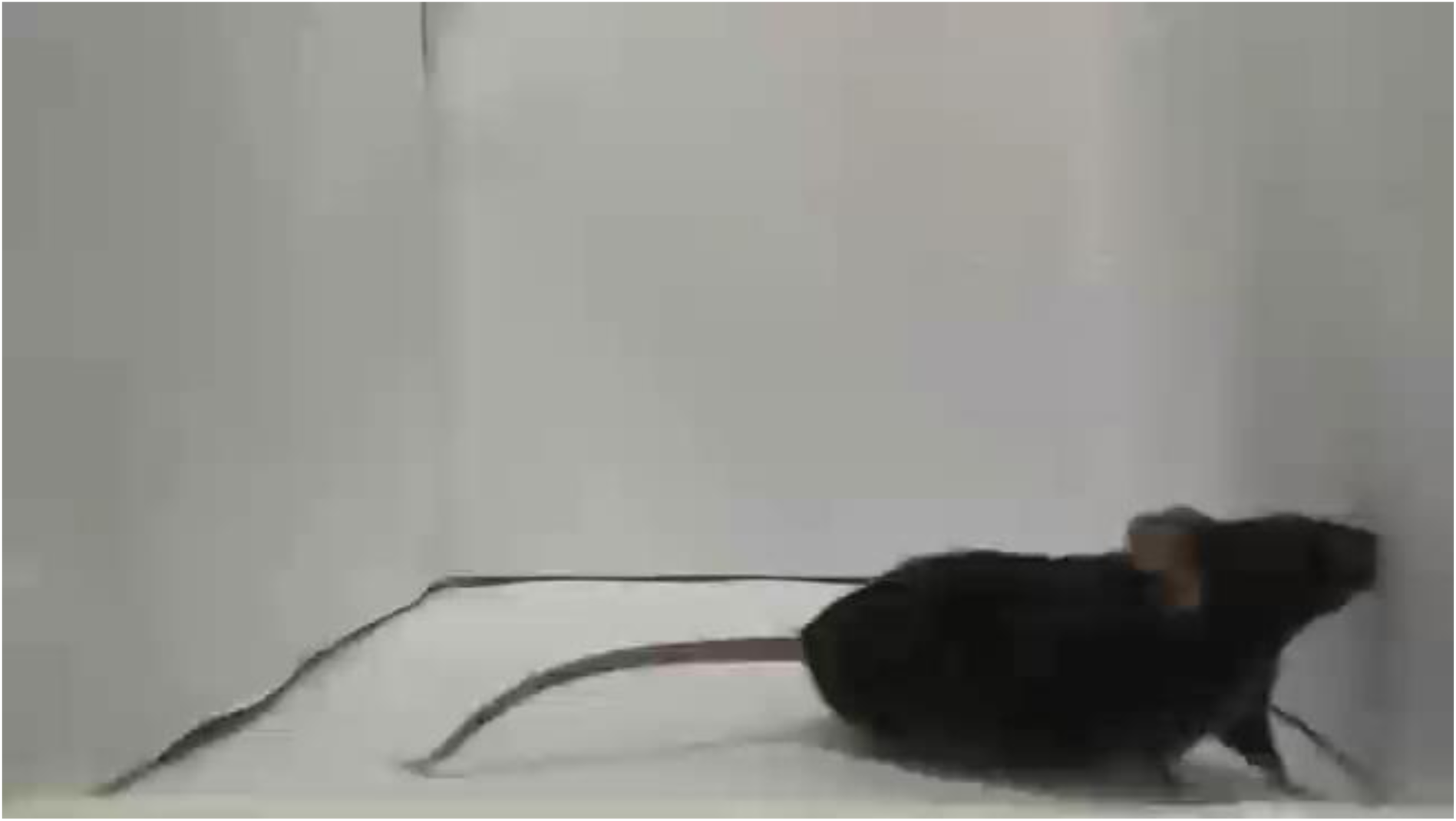
*Trim72* loss-of-function accelerates disease progression in FUS-R521C ALS mouse model. Locomotion of mice with the indicated genotypes at 1.5 year of age. Aged C/C;−/− animal displayed hindlimb weakness and lagging, trunk shaking, and impaired movements.

**Movie S2.**
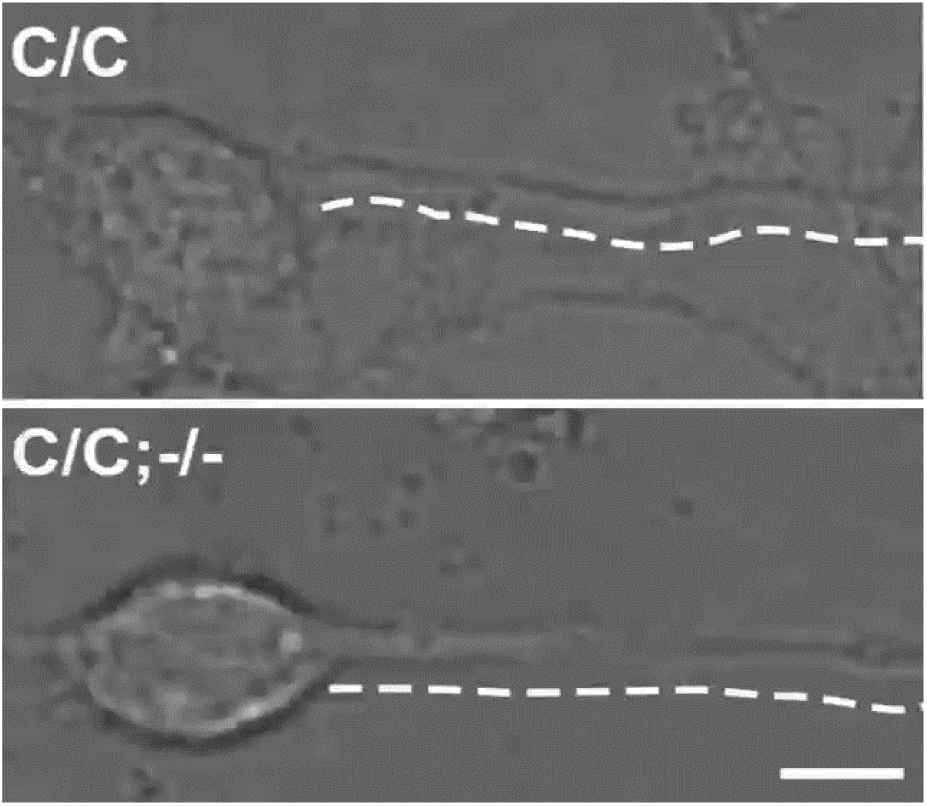
*Trim72* is required for integrity of neurite. Cultured cortical neurons with the indicated genotypes were photodamaged at the soma and imaged by time-lapse imaging. Note that neurites were labeled by dotted lines and broken neurites were labeled by arrowheads. Scale bar, 5 µm.

**Movie S3.**
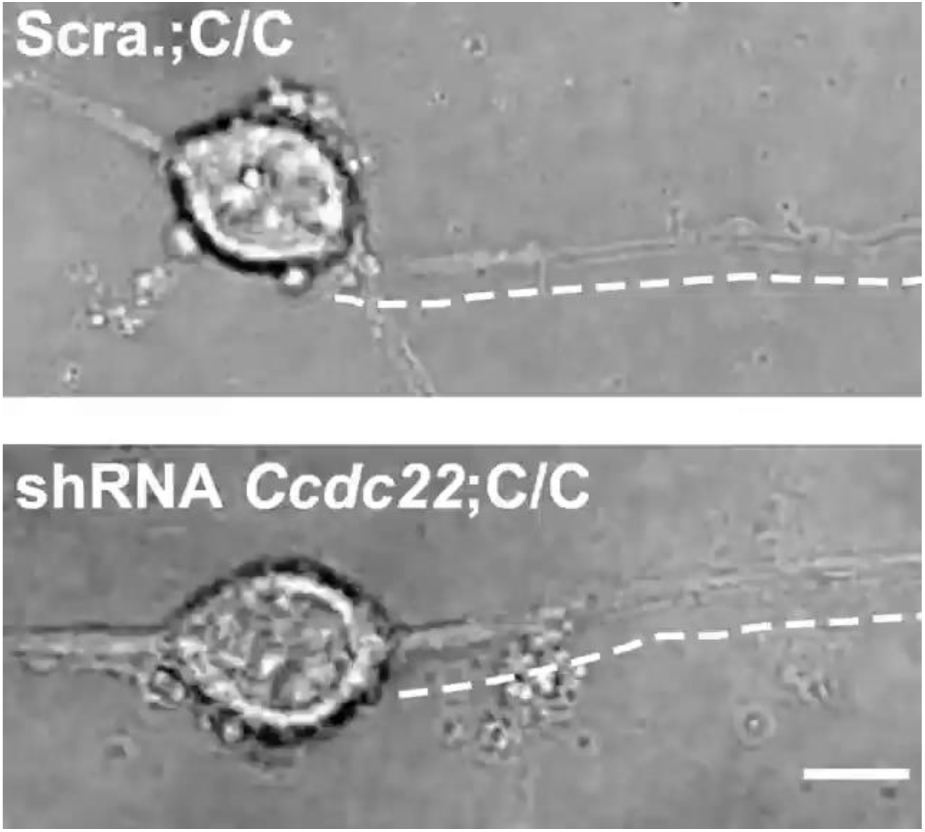
*Ccdc22* is required for integrity of neurite. Cultured cortical neurons (C/C genotype) infected with scrambled (Scra.) shRNA (negative control) or shRNA2 against *Ccdc22* were photodamaged at the soma and imaged by time-lapse imaging. Note that neurites were labeled by dotted lines and broken neurites were labeled by arrowheads. Scale bar, 5 µm.

**Movie S4.**
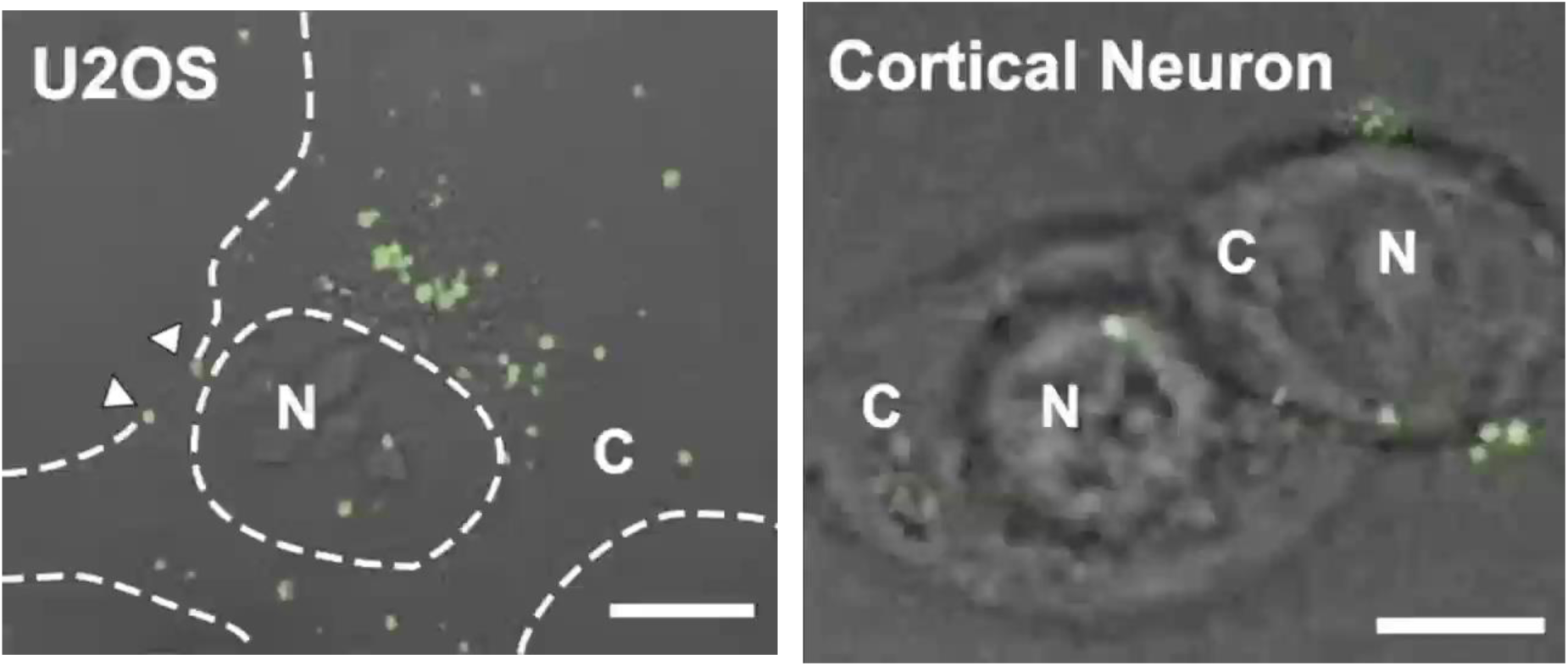
Extracellular applications of EGFP-TRIM72-containing exosomes in U2OS cells and cultured cortical neurons. Exosomes isolated from culture medium of 293FT cells expressing EGFP-TRIM72 were extracellularly applied to U2OS cells (Left). Dotted lines indicate the circumference of the nucleus (N) and the cell body (C). Arrowheads indicate EGFP+ exosomes that are in proximity to the cell surface. Exosomes expressing EGFP-TRIM72 extracellularly were applied to cultured cortical neurons (Right). Nucleus (N) and cytoplasm (C). Scale bar, 5 µm.

**Movie S5.**
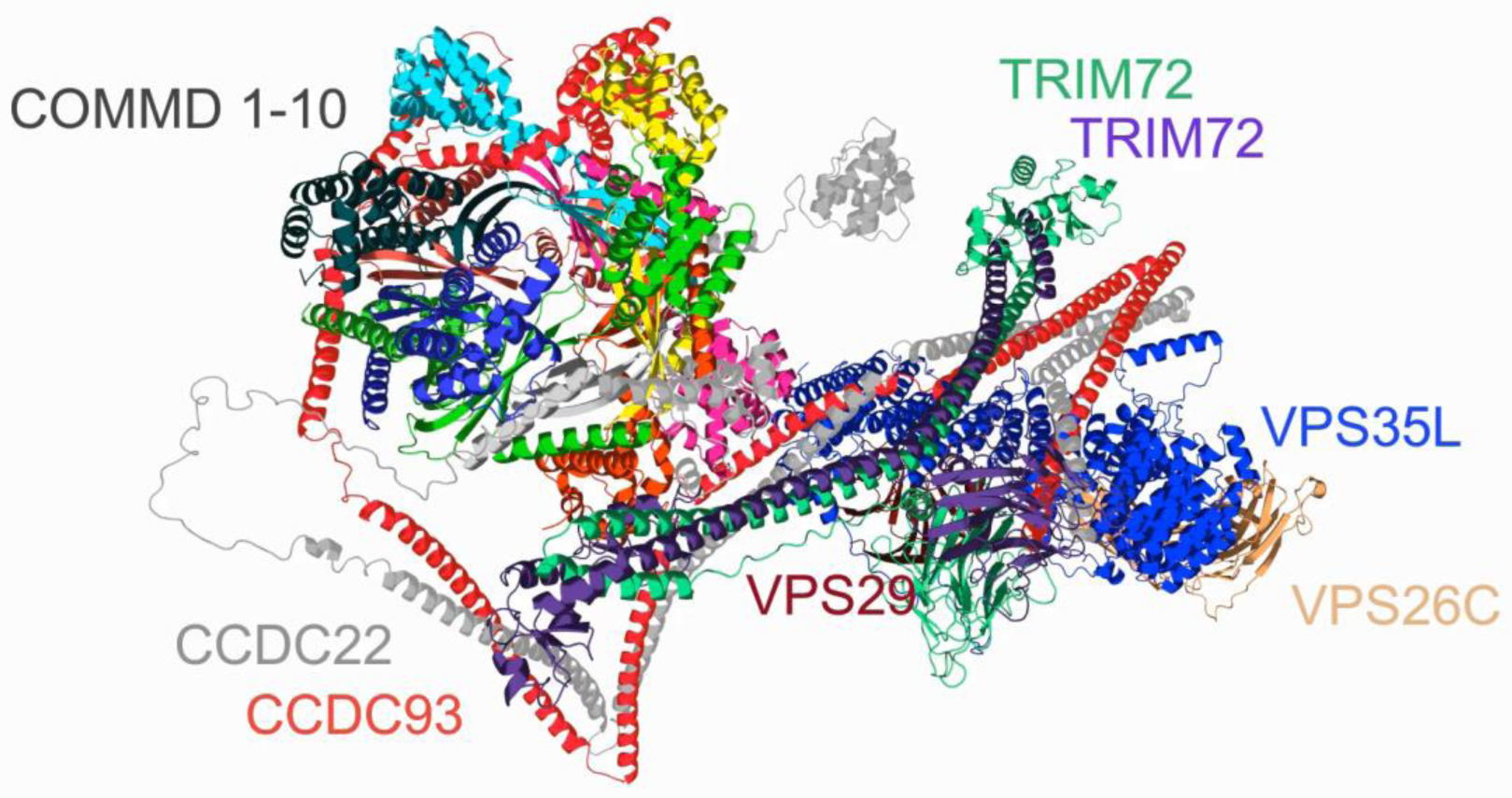
A 3D illustration of Commander/TRIM72 complex. A 3D model illustrates how TRIM72 interacts with CCDC22 to form Commander/TRIM72 complex, which facilitates vesicle recognition and membrane repair. The PRYSPRY domains of a TRIM72 dimer provides additional two sites for membrane anchoring (also see our working model in **Figure S34**).

